# Ancient *Yersinia pestis* genomes from across Western Europe reveal early diversification during the First Pandemic (541–750)

**DOI:** 10.1101/481226

**Authors:** Marcel Keller, Maria A. Spyrou, Christiana L. Scheib, Andreas Kröpelin, Brigitte Haas-Gebhard, Bernd Päffgen, Jochen Haberstroh, Albert Ribera i Lacomba, Claude Raynaud, Craig Cessford, Peter Stadler, Kathrin Nägele, Gunnar U. Neumann, Jessica S. Bates, Bernd Trautmann, Sarah Inskip, Joris Peters, John E. Robb, Toomas Kivisild, Michael McCormick, Kirsten I. Bos, Michaela Harbeck, Alexander Herbig, Johannes Krause

## Abstract

The first historically documented pandemic caused by *Yersinia pestis* started as the Justinianic Plague in 541 within the Roman Empire and continued as the so-called First Pandemic until 750. Although palaeogenomic studies have previously identified the causative agent as *Y. pestis*, little is known about the bacterium’s spread, diversity and genetic history over the course of the pandemic.

To elucidate the microevolution of the bacterium during this time period, we screened human remains from 20 sites in Austria, Britain, Germany, France and Spain for *Y. pestis* DNA and reconstructed six new genomes. We present a novel methodological approach assessing SNPs in ancient bacterial genomes, facilitating qualitative analyses of low coverage genomes from a metagenomic background. Phylogenetic analysis reveals the existence of previously undocumented *Y. pestis* diversity during the 6^th^–7^th^ centuries, and provides evidence for the presence of multiple distinct *Y. pestis* strains in Europe. We offer genetic evidence for the presence of the Justinianic Plague in the British Isles, previously only hypothesized from ambiguous documentary accounts, as well as southern France and Spain, and that southern Germany seems to have been affected by at least two distinct *Y. pestis* strains. Four of the reported strains form a polytomy similar to others seen across the *Y. pestis* phylogeny, associated with the Second and Third Pandemics. We identified a deletion of a 45 kb genomic region in the most recent First Pandemic strain affecting two virulence factors, intriguingly overlapping with a deletion found in 17^th^–18^th^-century genomes of the Second Pandemic.

**Significance Statement:** The first historically reported pandemic attributed to *Yersinia pestis* started with the Justinianic Plague (541–544) and continued for around 200 years as the so-called First Pandemic. To date, only one *Y. pestis* strain from this pandemic has been reconstructed using ancient DNA. In this study, we present six new genomes from Britain, France, Germany and Spain, demonstrating the geographic range of plague during the First pandemic and showing microdiversity in the Early Medieval Period. Moreover, we detect similar genome decay during the First and Second Pandemic (17^th^ to 18^th^ century) that includes the same two virulence factors, thus providing an example of potential convergent evolution of *Y. pestis* during large scale epidemics.

## Introduction

*Yersinia pestis*, the causative agent of plague, is a Gram-negative bacterium that predominantly infects rodents and is transmitted by their ectoparasites such as fleas. As a zoonosis, it is also able to infect humans with a mortality rate of 50–100 % without antibiotic treatment (1), manifesting as bubonic, septicaemic or bubonic plague. After the pathogen spread worldwide at the end of the 19^th^ century in the so-called Third Pandemic that started in 1855 in Yunnan, China, it established new local foci in Africa and the Americas in addition to the ancient foci that exist in Central and East Asia. Today, *Y. pestis* causes sporadic infections every year and even local recurrent epidemics such as documented in 2017 in Madagascar (2).

Although recent palaeogenetic analyses have been able to reconstruct an ancient form of *Y. pestis* that infected humans as early as in prehistoric times (2,800 to 1,700 BCE (3–5)) the First Pandemic (541–750) is the earliest historically recorded pandemic that has been clearly attributed to *Y. pestis* (6, 7), starting with the fulminant Justinianic Plague (541–544). It was later followed by the Second Pandemic, which started with the Black Death of 1347–1353 (8, 9) and persisted in Europe until the 18^th^ century (10–12).

First attempts in the 2000s aimed to amplify *Y. pestis-*specific DNA fragments from burials of the 6^th^ century (13–15). Although some early studies are controversial due to methodological limitations (16) and proved inconsistent with later work (17), more recent studies have been successful in reconstructing and authenticating whole *Y. pestis* genomes from two early medieval burial sites in modern-day Bavaria, Germany (6, 7).

These genomic investigations identified a previously unknown lineage associated with the First Pandemic that was found to be genetically identical in both sites and falls within the modern diversity of *Y. pestis*. Moreover, this lineage is distinct from those associated with the Second Pandemic that started ca. 800 years later, indicating two independent emergence events.

Although these studies have unequivocally demonstrated the involvement of *Y. pestis* in the First Pandemic, the published genomes represent a single outbreak, leaving the genetic diversity of that time entirely unexplored. Here, we assess the diversity and microevolution of *Y. pestis* during that time by analysing multiple and mass burials in a broader temporal and spatial scope than previously attempted. After screening 167 samples from 20 archaeological sites, we were able to reconstruct six new genomes with higher than 5-fold mean coverage from Britain, France, Germany and Spain. Furthermore, we identified a large deletion in the most recent First Pandemic strain that affects the same region as a deletion observed in late Second Pandemic strains, suggesting similar mechanism of pathogen adaptation in the waning period of the two separate pandemics.

## Results

### Screening and Capture

We used a previously described quantitative PCR assay (18) that targets the *Y. pestis*-specific *pla* gene on the pPCP1 plasmid to test 145 teeth from a minimum of 96 individuals from 19 sites (Table S1). All 19 PCR-positive extracts were subsequently turned into double-stranded libraries and enriched for *Y. pestis* DNA following an in-solution capture approach (19). Whereas some samples reached up to 9.6-fold chromosomal mean coverage after whole genome capture, four of the PCR-positive samples yielded a coverage of lower than 0.1-fold. Since the qPCR assay can amplify non-specific products and subsequent capture can enrich for environmental DNA that sporadically maps to the *Y. pestis* reference, it is crucial to differentiate between samples that show low DNA preservation and those that are false positives.

False positive samples are unlikely to show similar mapping success on all genetic elements when compared to true positive samples. Therefore, mapping to all three plasmids was used in combination with a statistical outlier detection for verification of low coverage genomes. Ratios of reads mapping to the *Y. pestis* chromosome and the three individual plasmids were determined, and samples were authenticated by calculating the Mahalanobis distance to detect outliers (χ^2^=9.210, df=2, p=0.01; Table S2). Two samples, DIR002.A and PEI001.A, were classified as outliers: despite having chromosomal coverage, they had no or only a few reads mapping to the plasmids and were therefore considered as *Y. pestis* negative. The remaining 17 samples come from four sites in Germany (Dittenheim [DIT], n=3; Petting, n=3; Waging [WAG], n=1; Unterthürheim [UNT], n=5), one in Spain (Valencia [VAL], n=1) and one in France (Lunel-Viel [LVC], n=6) (Table 1, Fig. 1).

**Table 1:**
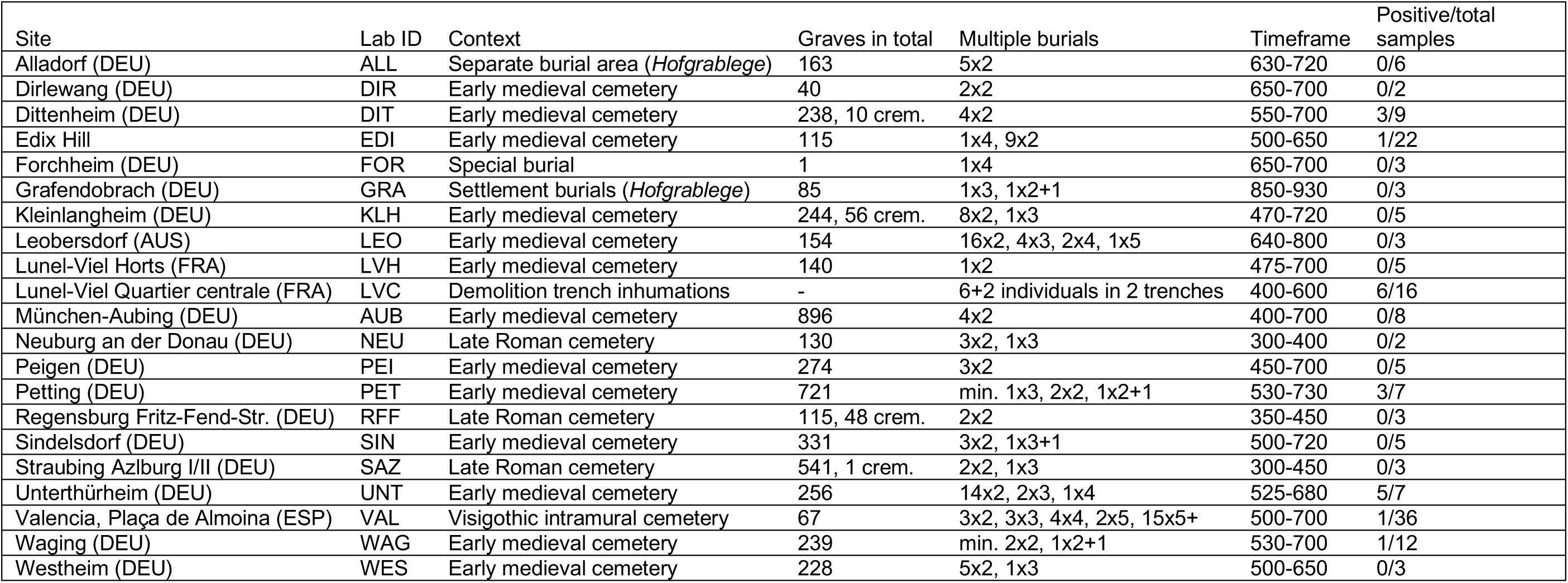
List of all sites that were tested with country in brackets (AUS=Austria, DEU=Germany, ESP=Spain, FRA=France). The number of graves is counting multiple burials as single graves; cremations are counted separately. Multiple burials are listed as number of graves times number of individuals (5×2 translates to 5 double burials, 1×2+1 to one double burial associated with a single burial). Detailed site descriptions are given in the SI, a table of all screened samples in Table S1.

**Fig. 1:**
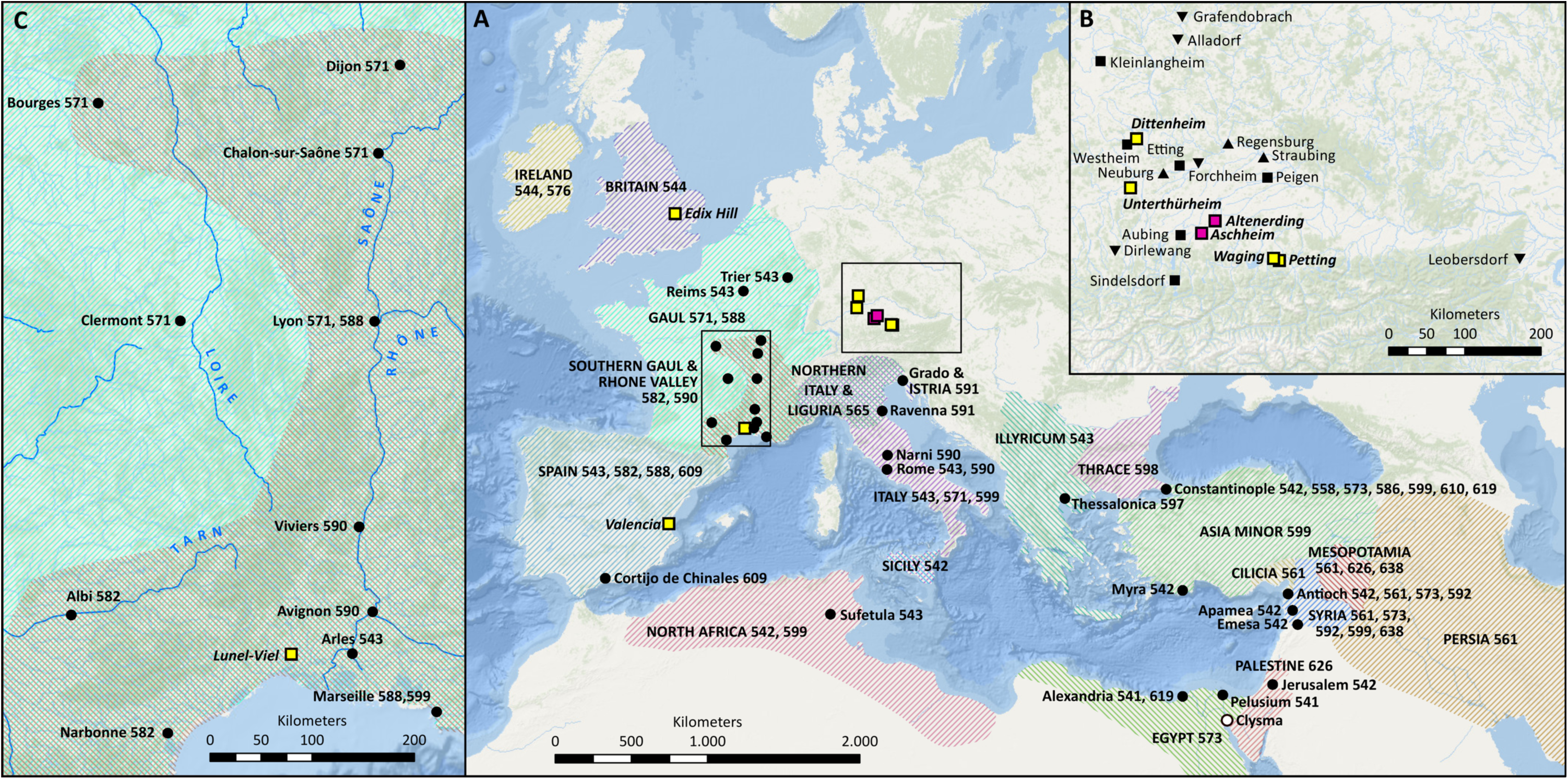
Geographic extent of the First Pandemic and sampled sites. A: Map of historically documented occurrences of plague (regions shaded, cities depicted by circles, both with respective years of occurrence) between 541 and 650 in Europe and the Mediterranean basin. All sources are given in the SI. Sites with genomic evidence for *Y. pestis* are shown as pink (previously published) and yellow squares (presented here). B: Enlarged rectangular space of A (right) showing all sites in Germany and Austria that were included in this study. Sites tested negative are depicted in black upward-pointing triangles (burials dating before 541), squares (dating around 541–544) downward-pointing triangles (dating after 544). C: Enlarged inset of A (left) shows reported occurrences in France and main rivers.

After mapping to the chromosome, seven genomes showed a higher than 5-fold mean coverage and were used for downstream analyses. These were DIT003.B (9.4-fold), VAL001.B (9.6-fold), PET004.A (5.6-fold) as well as UNT003.A and UNT004.A (7.6-fold and 5.2-fold respectively) (Table S3). Six positive samples of the individuals LVC001, LVC005 and LVC006 were merged to yield a mean coverage of 6.7-fold for the site of Lunel-Viel. For the phylogenetic analysis, we omitted the lower-covered UNT004.A after assuring that there are no conflicting positions with UNT003.A that derives from the same archaeological site.

For the 22 samples from Edix Hill, Britain, only shotgun sequencing data was available and, therefore, pathogen DNA screening was performed using the metagenomic tool MALT (19). This analysis revealed six putatively *Y. pestis*-positive samples after visual inspection of aligned reads in MEGAN (20) (Table S4). The sample EDI001.A had more than 9000 reads assigned to *Y. pestis* and was sequenced to a greater depth without enrichment to yield a mean chromosomal coverage of 9.1-fold.

### SNP Evaluation

In the context of ancient pathogen DNA, there are three possible sources for false positive SNPs: First, DNA damage such as deamination of cytosine to uracil can lead to misincorporation of nucleotides during sample processing (21). Second, the mapping of closely related environmental species to the reference sequence of the target organism is likely, especially for conserved regions of the genome (22). Third, mapping of short reads is more prone to mismapping and calling of false positive SNPs generated at sites of genome rearrangement. Whereas the first source can be circumvented via *in vitro* protocols like UDG treatment (23), the latter two can be reduced but not eliminated with strict mapping parameters and exclusion of problematic regions (24) as applied here. A fourth source for false SNP assignments could result from multiple genetically distinct strains that would lead to a chimeric sequence. The later was not observed in our data (Fig. S1) and this phenomenon might be limited to chronic infections with pathogens such as *Mycobacterium tuberculosis,* where mixed infections have been previously documented (25).

The retrieval of genomes that span a wide geographic area gives us the opportunity to assess *Y. pestis* microdiversity present in Europe during the First Pandemic. Given that our genomes are of relatively low genomic coverage, we critically evaluated uniquely called and shared SNPs among the First Pandemic genomes in order to accurately determine their phylogenetic position. This analysis was performed for all genomes retrieved from UDG-treated libraries with higher than 5-fold mean coverage, including the previously published high-quality Altenerding genome (17.2-fold mean coverage).

For this, we developed the tool ‘SNPEvaluation’ and defined three different criteria, all applying for a 50 bp window surrounding the SNP: (A) Comparing the mean coverage after BWA mapping with high and low stringency and excluding all SNPs that showed a higher coverage under low stringent mapping than in high stringent mapping. In metagenomic datasets, reads of related species map frequently to conserved regions in the reference genome. When the position is not covered by reads from the target organism (*Y. pestis*) but the genomic region is similar enough in other environmental organisms so that their reads can map, they might mimic a SNP in *Y. pestis* when the contaminant species carries a different allele in that position. (B) Excluding all SNPs for which heterozygous calls were identified in the surrounding regions. Heterozygous calls accumulate in conserved regions due to the above-described effect. (C) Excluding all SNPs within regions that include positions that lack genomic coverage. Variants in genome architecture often appear as gaps in mapped data and are likely to cause mapping errors, potentially resulting in false positive SNPs.

This evaluation was applied to all SNPs identified as unique to the First Pandemic lineage, totalling between one and 15 per genome, respectively (Table S5). 17 authentic derived chromosomal SNPs and an additional one detected on the pMT1 plasmid were found across all seven genomes (Table S6). The Altenerding genome (AE1175) as well as the genomes of Unterthürheim (UNT003.A, UNT004.A) and Dittenheim (DIT003.B) appear identical after SNP evaluation at all positions, with the exception of one SNP which is covered in both UNT samples, and by only one read in DIT003.B, but not covered in Altenerding. The genomes from Petting (PET004.A), Valencia (VAL001.B) and Lunel-Viel (LVC00_merged) appear distinct, each occupying a unique branch comprised of two, three and ten unique SNPs, respectively (Fig. 2B and Table S6). One additional SNP was found on the pMT1 plasmid in the Valencia genome (VAL001.B). An analysis of the Aschheim genome as well as a SNP effect analysis is presented in the SI.

**Fig. 2:**
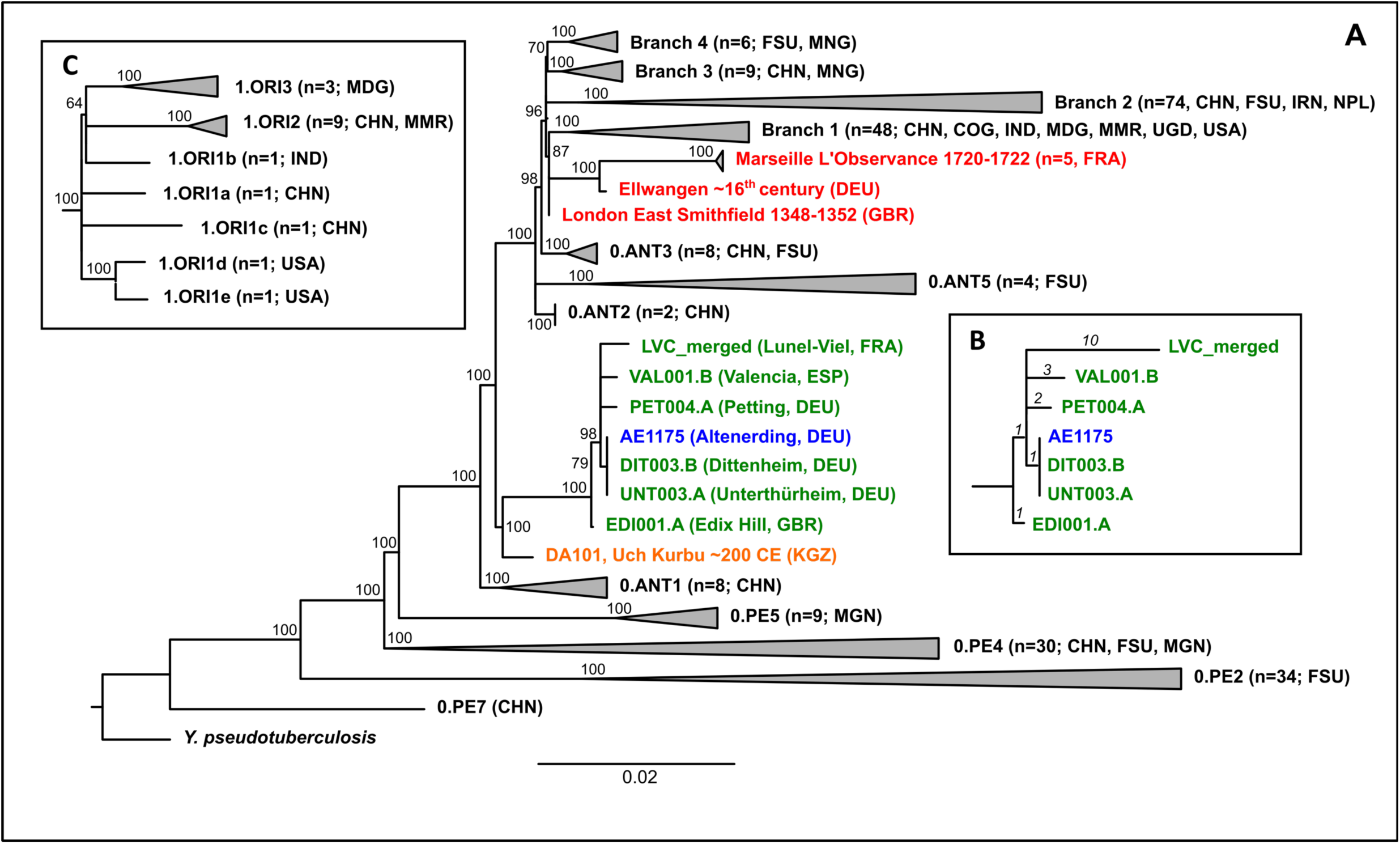
Phylogenetic tree. A: Maximum Likelihood tree with full SNP alignment (6496 positions) of 233 modern *Y. pestis* and one *Y. pseudotuberculosis* genome, nine published (2^nd^ to 3^rd^ century Tian Shan in orange; Altenerding in blue; Second Pandemic in red) and six genomes presented here (green) with country given in brackets (DEU=Germany, ESP=Spain, FRA=France, GBR=Great Britain). Numbers and origins of modern genomes are given in brackets (CHN=China, COG=Congo, FSU=Former Soviet Union, IND=India, IRN=Iran, MDG=Madagascar, MMR=Myanmar, MNG=Mongolia, NPL=Nepal, UGA=Uganda). Numbers on nodes are showing bootstrap values (1000 iterations). B: Detailed, manually drawn tree of the First Pandemic genomes showing all remaining SNP positions after SNP evaluation (number of SNPs given in italics). C: Detailed tree of the 1.ORI clade within Branch 1, showing the polytomy.

Since the Altenerding genome shows the highest coverage (6), all SNPs previously presented as unique SNPs for this genome were evaluated as potentially shared SNPs when they appeared in at least one of the new genomes – excluding the position shared exclusively with DIT003.B and UNT004.A. We applied the exact same parameters as for the unique SNPs, but also considered positions with less than 3-fold coverage (Table S7). Only SNPs that pass all three criteria of our SNP evaluation in at least half of the analysed genomes (i.e., four out of seven) were accepted as true shared SNPs, reducing the number from 53 identified in a previous study (6) to 43.

The Waging sample (WAG001.A) had a genomic coverage too low for inclusion in our phylogenetic analysis. Since it was the only sample giving evidence for *Y. pestis* presence at this site, it was assessed for all SNPs that were either shared or unique in the other First Pandemic genomes. Visual inspection revealed seven of the 43 shared SNPs to be present in the WAG001.A genome at low coverage (<3-fold), but none of the unique ones. For both shared and unique SNPs, no conflicting positions were found. This strain could, therefore, be attributed to the First Pandemic lineage without, however, resolving its exact position (Table S5).

The genome reconstructed from the non-UDG library of EDI001.A was not considered in the presented SNP analysis, since damaged sites interfere with the defined SNP evaluation criteria. With a mean coverage of 9.1, we accept the two relevant positions (one ancestral and one derived SNP) as true positive after visual inspection. Regardless, metrics determined in the SNP evaluation are still reported for comparison.

### Phylogenetic analysis

A set of 233 modern *Y. pestis* genomes (Table S8) as well as seven Second Pandemic genomes, including a representative of the Black Death strain (London) and six post-Black Death genomes (16^th^-century Ellwangen (11); 18^th^-century Marseille (12)), and an ancient genome from Tian Shan (DA101, 2^nd^ to 3^rd^ century (26)) were used for phylogenetic analyses alongside our First Pandemic genomes presented here (Table S3) and the previously published genome of Altenerding. The *Y. pseudotuberculosis* isolate IP32953 (27) was used as an outgroup.

Our maximum likelihood tree (28) constructed from the full SNP alignment reveals that all of the genomes presented here occupy positions on the same lineage (Fig. 2A, Fig. S2). This confirms their authenticity and is congruent with previous association of this lineage to the First Pandemic (541–750). In addition, the previously reported genome from Altenerding (2148) is identical to the new genomes from Dittenheim (DIT003.B) and Unterthürheim (UNT003.A). Moreover, the genomes of Petting (PET004.A), Valencia (VAL001.B) and Lunel-Viel (LVC_merged) seem to diverge from the Altenerding cluster through a polytomy (Fig. 2C, bootstrap support 98 %). The British genome of EDI001, however, branches off one SNP ancestral to this polytomy (bootstrap support 100 %) and possesses one unique SNP. This is remarkable, since the British Isles are one of the most remote places where the First Pandemic has been suspected to have reached in relation to the presumed starting point in Egypt.

### Virulence factor and deletion analysis

We screened for the presence/absence of 80 chromosomal and 42 plasmid associated virulence genes (29, 30) in all First Pandemic genomes with higher than 5-fold coverage (Fig. 3 and Fig. S3). Only the filamentous prophage was consistently found absent in all presented genomes. This is expected, since it has integrated into the genome of only a number of modern Branch 1 genomes (31). Reduced coverages for a set of virulence factors can be seen in the Altenerding (AE1175) and Ellwangen genomes due to a capture bias, since the capture probe set in the respective studies was designed on the basis of *Y. pseudotuberculosis* rather than of *Y. pestis* (6, 11).

**Fig. 3:**
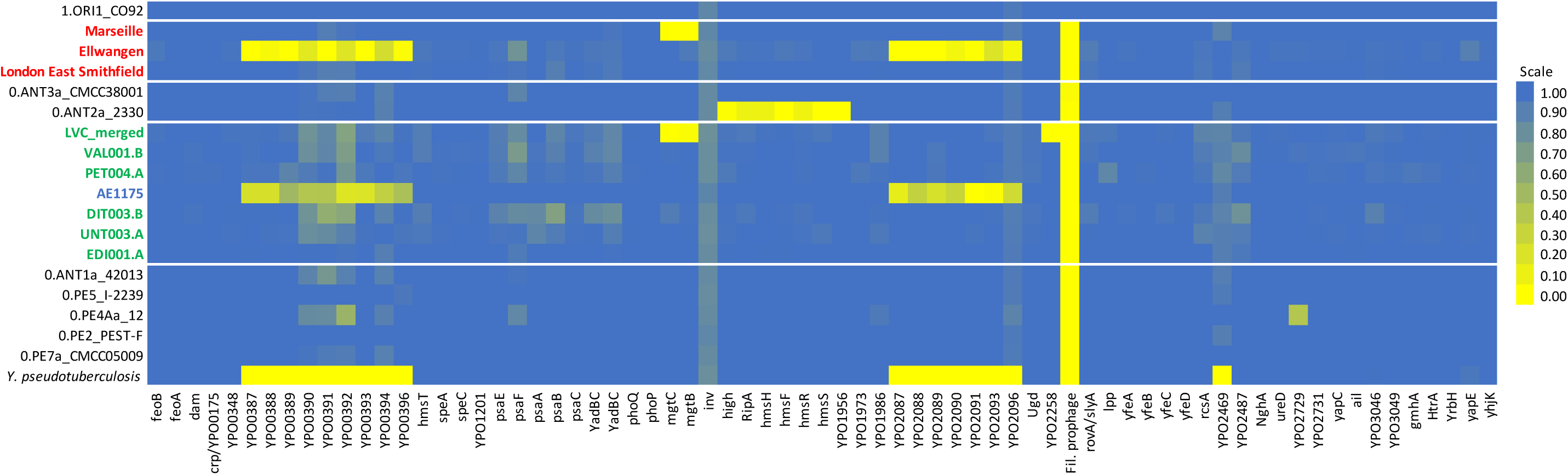
Heatmap showing the percentage of coverage of chromosomal virulence factors. First Pandemic genomes (blue and green) and Second Pandemic genomes (red) are shown in combination with selected strains of main clades of modern *Y. pestis* diversity on Branch 0 as well as the reference genomes of *Y. pseudotuberculosis* and *Y. pestis* (CO92).

Intriguingly, the most derived First Pandemic genome from Lunel-Viel shows a deletion of two chromosomal virulence factors, *mgtB* and *mgtC* (Fig. 3). These magnesium transporters are part of the PhoPQ regulon, which is important for survival of *Y. pestis* in the magnesium-deficient environment of macrophages. However, functional studies on *mgtB* hint at an important role during macrophage invasion rather than intracellular survival (32).

A second deletion was observed for the gene YPO2258, categorized as a potential virulence factor based on the presence of a frame shift mutation in the avirulent 0.PE2_Microtus91001 strain (30). Its inactivation in the 2.ANT1_Nepal516 strain, isolated from a human patient, nevertheless indicates that this gene is not essential for virulence in humans (33).

Further exploration of the deletion of the two neighbouring genes *mgtB* and *mgtC* revealed that they are part of a ca. 45 kb deletion (positions 1,883,402 to 1,928,869 in the CO92 reference), affecting 34 genes including multiple motility (*motA, motB*) and chemotaxis genes (*cheA, cheB, cheD, cheR, cheW, cheY, cheZ*) (Fig. S4). On the downstream end, the deletion is flanked by an IS100 insertion element. A potential upstream insertion element might be undetectable at our current resolution due to a genome rearrangement in the reference genome CO92. This is in agreement with previous findings concerning the highly abundant IS100 element in *Y. pestis*, responsible not only for disruptions of multiple genes caused by homologous recombination (27), but also for the loss of the 102 kb long *pgm* locus containing a high-pathogenicity island in several strains (34). To address the specificity of this deletion to the 6^th^–7^th^ century strain from France, we also investigated the presence of the two virulence factors in all other modern and ancient strains in this study. Intriguingly, a similar deletion affecting the same region including *mgtB* and *mgtC* was observed in the late Second Pandemic genomes from London New Churchyard, (1560-1635 (35)) and Marseille L’Observance (1720-1722 (12)). However, a full deletion of this 45 kb region was not found in any of the other ancient or modern genomes. Therefore, the deletion appeared independently in the course of both the First and Second Pandemics.

## Discussion

### Identifying *Y. pestis* DNA in low complexity specimens

In total, we screened 145 samples from 19 sites in France, Germany and Spain for *Y. pestis* with a qPCR assay (18) and 22 additional samples from Edix Hill, Britain, with the metagenomic tool MALT (19). While the most promising sample of Edix Hill was directly sequenced to greater depth to reach a chromosomal mean coverage of 9.1-fold, all qPCR positive samples of the other sites were turned into UDG libraries and subsequently enriched for *Y. pestis*, resulting in mean coverages ranging from 0.01 to 9.6-fold.

The validation of ancient genomes with relatively low coverage as presented here is challenging since the DNA extracted from archaeological remains results in metagenomic data and the differentiation between target organism DNA and environmental background can be difficult. The identification of *Y. pestis* DNA based on PCR targeting the *pla* locus on the pPCP1 plasmid has theoretically been shown to be problematic (36), leading to discussions about false positive results (16). However, assignment to *Y. pestis* based on reads retrieved from shotgun sequencing and mapping to a reference genome also can be challenging in case of extremely low genomic coverage (3, 4). Since all the presented genomes, except the one from Edix Hill, are derived from DNA libraries specifically enriched for *Y. pestis* DNA and are thus biased towards the target organism, a previously suggested competitive mapping approach (3) would not be suitable. Instead, we considered the relative number of mapping reads to the plasmids and chromosome to identify false positive samples from captured data. We were able to verify that 17 out of 19 samples were positive for *Y. pestis* with as few as 4000 reads mapping to the chromosome. Since the three plasmids pCD1, pMT1 and pPCP1 were already present in the early divergent Neolithic and Bronze Age strains (3, 4) and loss of plasmids has only been observed sporadically in attenuated strains (37), this method could be reliably applied to data stemming from other branches in the *Y. pestis* phylogeny.

### Analysing microdiversity with low coverage genomes

Reliable SNP calling is crucial for the phylogenetic analysis of verified low coverage genomes and can be challenging when dealing with ancient pathogen DNA stemming from metagenomic contexts. This has been demonstrated on *Y. pestis* genomes (6), but previously applied visual inspections are time-consuming and not easily reproducible.

Here, we present a novel approach for SNP authentication using a semi-automated SNP evaluation. We selected three criteria for our evaluation to assess the likelihood of mismapping. We excluded all SNPs that (A) had higher coverage when mapped with less strict parameters, (B) had ‘heterozygous’ positions in close proximity or (C) were flanked by gaps. With these filters, we tolerate a loss of sensitivity to increase specificity, which is critical for detection and characterization of microdiversity. Moreover, the tool ‘SNPEvaluation’ that was newly developed for this analysis offers a highly flexible framework for the assessment of VCF files and can be utilized also for a variety of analyses on different organisms.

### Phylogenetic analysis

We were able to confidently reconstruct six new genomes from the First Pandemic in Britain, France, Germany and Spain, providing insights into the microdiversity of *Y. pestis* in Europe between the 6^th^ and 7^th^ centuries.

Our presented genomes add diversity to a phylogenetic lineage that was previously shown to contain two identical 6^th^ century genomes of southern Germany (Aschheim and Altenerding (6, 7)). It diverges between the 0.ANT1, 0.ANT2 and 0.ANT5 clades in the main *Y. pestis* phylogeny and shares a short branch with a 2^nd^-to 3^rd^-century genome from the Tian Shan mountains (26). Intriguingly, a single diversification event gave rise to the published as well as three of the presented additional genomes, each defined by 1 to 10 derived SNPs. Similar polytomies can be detected in other parts of the phylogeny of *Y. pestis* that have been related to human epidemics (38): one gave rise to Branches 1 to 4 (including ancient Second Pandemic genomes, Fig. 2A) and is dated to 1142-1339 (38), shortly before the European Black Death. To date it is unknown if this event was restricted to a rodent reservoir, or if it was already associated with a human epidemic. A second polytomy gave rise to the 1.ORI clade, which includes strains related to the worldwide spread of plague during the Third Pandemic in the 19^th^ century (Fig. 2C).

Within the First Pandemic lineage, the genomes that derive from this polytomy display variable terminal branch lengths (1-10 SNPs), which are likely concurrent with their different ages (see below). Given that *Y. pestis* is a pathogen that can cover large geographic distances by accumulating little to no genetic diversity (11), it is challenging to elucidate the geographic origin for this diversification event. A first hypothesis suggests an origin of this diversification event within the historically recorded geographic range of the First Pandemic, i.e. either in Europe, the Mediterranean basin, or the Middle East. Our current data may lend some credibility to this scenario for two reasons: First, we identify four European strains with short genetic distances from this polytomy, the shortest of which is identified in three locations in rural Bavaria, and second, we identify an almost direct ancestor of this polytomy to be present in Europe during the 6^th^ century, represented by a genome from Britain. Alternatively, the bacterium may have been recurrently introduced to the affected regions from a single remote reservoir.

The hypothesis of a single introduction would require the establishment of a local reservoir, since we have to assume that at least the genome recovered from Lunel-Viel is not directly associated with the initial outbreak in 541–544 but rather with a subsequent one (see below). Possible locations for reservoirs during the First Pandemic have been suggested in the Iberian peninsula and the Levant (39). There is also a growing body of evidence for the presence of black rats (*Rattus rattus)* in Europe in late Antiquity and the Early Medieval Period (40, 41), suspected to represent the main reservoir species during the Second Pandemic (40).

Such a scenario would be congruent with the Second Pandemic, where the phylogeny of ancient genomes is in line with a single introduction and subsequent persistence in a local host (11, 35, 42), although this hypothesis was challenged by an alternative scenario claiming multiple introductions on the basis of climatic data (43). Similar to the European Second Pandemic lineage (11, 12), strains emerging from the First Pandemic lineage have so far been recovered solely from ancient DNA of European plague burials, suggesting that the lineage either went extinct or persists in a yet unsampled reservoir.

### Origin of the Justinianic Plague

Based on available data, it has been suggested that the most parsimonious location for the divergence event that gave rise to the First Pandemic lineage is Central Asia (26). All published genomes of the branches 0.ANT1, 0.ANT2 and 0.ANT5 that frame the First Pandemic lineage in the phylogenetic tree were sampled in the autonomous Xingjiang region in north-western China or in Kyrgyzstan (38, 44). In addition, an ancient 2^nd^-to 3^rd^-century *Y. pestis* genome from the Tian Shan mountains in Central Asia (26) branches off basal to all the First Pandemic genomes. The resulting claim that the Huns might have brought plague to Europe is however unsubstantiated due to the gap of more than three centuries prior to the onset of the First Pandemic.

Since the long shared branch of the First Pandemic genomes (43 SNPs) does not have any known extant descendants, this strain might have been maintained in a now extinct reservoir after its emergence in Central Asia. The first outbreak is reported in Pelusium, Egypt; an introduction from either Africa or Asia was presumed, given the sudden and dramatic onset of the pandemic. Previous assumptions of an African origin were mainly based on a single deeply diverging 0.PE strain ‘Angola’ (45) and the reports of the Byzantine historian Evagrius Scholasticus, who wrote in his *Ecclesiastical History* that the plague began in “Ethiopia”. However, there are legitimate doubts about the characterization of the ‘Angola’ genome as a genuine African strain (24, 46) and the account of Evagrius has been assessed critically with historical and philological methods (47, 48). For an Asian origin, the sea route via the Red Sea and the Indian Ocean is a plausible scenario since India was well connected by marine traffic with the early Byzantine Empire (39). A suggested alternative scenario would require overland transport from the Eurasian Steppe via Iran to the Red Sea that is, so far, not supported by any data (49). In conclusion, we interpret the current data as insufficient to resolve the origin of the Justinianic Plague as a human epidemic.

### Archaeological and historical context

Here, we present the first genomic evidence for the First Pandemic reaching the British Isles in the 6^th^ century. This genome was recovered from a burial on the site of Edix Hill, close to Cambridge (Roman *Duroliponte*) and near a Roman road running north from London (*Londinium*) toward Lincoln (*Lindum Colonia*) via Braughing, all of which were Roman settlements. Based on archaeological dating in combination with its rather basal position within the clade, this genome is likely related to the very first occurrence of plague in Britain suggested for 544 (see SI). The close proximity to the trade center of ancient London supports that the strain was introduced by sea communications, e.g., with Brittany, following the outbreak in central Gaul in 543 (Fig. 1, (50)). Interestingly, the genome was recovered from a single burial, underlining that in small settlements, plague-induced mortality crises need not always involve a radical change in mortuary practice towards multiple or mass burials. The fact that three of the five additional Edix Hill individuals that appeared positive for plague in the MALT screening were buried in two simultaneous double burials nevertheless suggests that multiple burials in normal cemeteries are indeed a good indicator for epidemic events (17).

In addition, we were able to reconstruct two genomes from the Mediterranean basin, where the historical records are more explicit about the presence of plague during the First Pandemic. Regarding Spain, the radiocarbon dating of the *Y. pestis*-positive individual from Valencia (432-610) would include the first outbreak reported for Spain in 543 in a contemporary chronicle (see SI). The three unique SNPs identified in this genome, which separate it from the identified polytomy may suggest its association with a later outbreak. Intriguingly, a canon of a church council held in 546 in Valencia dealing with burial practices for bishops in case of sudden death was recently connected with plague by philological and contextual analysis (51). Later outbreaks within the relevant timeframe are documented in Spain’s Visigothic kingdom, e.g., in 584 and 588 by Gregory of Tours, and by a funerary inscription dated 609 at Cortijo de Chinales 35 km northeast of Malaga (31).

The second Mediterranean genome from Lunel-Viel in southern France likely represents another outbreak, since it forms an independent strain which derives from the same polytomy as the Spanish and German genomes. The radiocarbon dates for the inhumations give an interval of at least 567-618 (youngest lower and oldest upper boundary, Table S9) overlapping with documented outbreaks in 571, 582, 588, 590 and possibly 599–600 in southern France (see Fig.1A and C). Lunel-Viel’s broader vicinity includes Arles, the seaport city of Marseille and the Rhône mouth. Close to important coastal and fluvial shipping routes as well as Roman roads that facilitated the spread of plague (39), Lunel-Viel could have been affected by all five recorded epidemics. The initial outbreak, documented for Arles ca. 543, falls outside of some of the radiocarbon intervals. This is consistent with the phylogenetic analysis that shows a higher accumulation of SNPs in this genome. Thus the victims at Lunel-Viel can most likely be attributed to one of the subsequent outbreaks.

Moreover, within Bavaria, Germany we detected *Y. pestis* in four sites (Dittenheim, Petting, Unterthürheim, Waging) in addition to the two previously published sites (Altenerding (6), Aschheim (7)). Two of the reconstructed genomes were identical to Altenerding and Aschheim, suggesting that these four can be attributed to the same epidemic event. Some of the radiocarbon intervals of these sites fall even slightly before the onset of the First Pandemic, suggesting an association of this outbreak directly with the Justinianic Plague. Regarding the Edix Hill genome, this would in turn necessitate the accumulation of one (Edix Hill) to two (Altenerding cluster) SNPs within the onset of the First Pandemic between 541–544.

Intriguingly, the genome of Petting, Bavaria falls not with the Altenerding cluster but in a distinct phylogenetic position. Since this strain also branches off from the common node with the other Bavarian strain as well as the French and Spanish genomes, this shows the presence of two independent strains and, therefore, presumably two independent epidemic events in early medieval Bavaria. This is striking, since we lack any historical records of the First Pandemic affecting southern Germany. The radiocarbon dates for the Bavarian sites are inconclusive and do not allow for a clear temporal separation of the two events. The higher number of accumulated SNPs nevertheless suggests a younger date for the epidemic represented by Petting. Further phylogeographic analyses are presented in the SI.

### Deletion analysis

The analysis of virulence factors revealed a deletion of a ca. 45 kb region in the most derived and putatively most recent genome thus far identified for the First Pandemic. This deletion contained two previously described virulence factors involved in host cell invasion and intracellular growth (*mgtB* and *mgtC*). Intriguingly, a similar deletion covering the same genomic region was detected in the most derived available Second Pandemic genomes from London New Churchyard (1560-1635) and Marseille (1720-1722). Genome decay by deletion or pseudogenization is a well-known trait of *Y. pestis* and has contributed to its distinct ecology and pathogenicity (53). Both deletions from the First and Second Pandemic are observed in genomes recovered from human victims. Therefore, it is reasonable to assume that the deletion may not have reduced the bacterium’s virulence. Moreover, it affects a number of cell surface proteins – remnants of the motile lifestyle of non-pestis *Yersiniae* (54) – so the deletion might have even facilitated immune evasion.

Because none of the investigated modern strains harboured this specific deletion, this possible case of convergent evolution might be an adaptation to a distinct ecological niche in Europe or the Mediterranean basin since an ancient local reservoir is the most parsimonious hypothesis for both historical pandemics (12, 42).

### Concluding remarks

Our study succeeds in offering new insights into the first historically documented plague pandemic, complementing the limited power of conventional historical, archaeological or palaeoepidemiological research. Moreover, we show the potential of palaeogenomic research for understanding historical and modern pandemics by a comparative approach on genomic features throughout millennia. Facing the problem of low coverage genomic data with a high environmental background – a notorious challenge in ancient DNA research –, we have developed new approaches to facilitate the authentication and confident phylogenetic placement of such genomes.

In the future, more extensive sampling of putative plague burials will help to draw a more comprehensive picture of the onset and persistence of the First Pandemic, especially on sites in the eastern Mediterranean basin, where not only is the Justinianic Plague reported to have started, but where also the 8^th^ century outbreaks also clustered. This will contribute to the comparative exploration of *Y. pestis’* microevolution and human impact in the course of past and present pandemics.

## Material and Methods

### Sites and Samples

The acquisition and selection of samples followed two approaches: Focussing on Bavaria, we concentrated on one region, where the two previously reconstructed *Y. pestis* genomes attributed to the Justinianic Plague had been found (6, 7). Additionally, given the absence of robust genetic evidence from the Mediterranean basin, which historical records depict as the epicenter of the pandemic, and the controversial presence of plague on the British Isles during the Justinianic Plague, we extended our screening to three sites with multiple burials in a broader geographical scope on the Mediterranean coast in France and Spain and inland Britain. Table 1 gives an overview of all tested sites.

For the first focus, we collected samples of 79 individuals from 46 burials belonging to 16 archaeological sites in Bavaria, Germany and one site in Austria (See Fig. 1B). The dating of the burials spans the 4^th^ to 10^th^ century, including also burials dating before (8 individuals on 3 sites) and after (17 individuals on 5 sites) the Justinianic Plague (541–544). Since mass graves that could be indicative of an epidemic are unsurprisingly rare for the small settlements associated with early medieval cemeteries in Bavaria, we followed the approach of the previous successful studies (6, 7, 17): we systematically screened multiple burials, i.e., where two or more individuals were found in a context indicating a simultaneous burial, such as a common grave pit and articulated remains on the same level. Single burials were sporadically tested, if the context suggested a close connection to a multiple burial. Burials with indications of a violent death of the interred were excluded, since a coincidental acute infection with *Y. pestis* seems unlikely.

Within the Mediterranean basin, we tested inhumations from Valencia, Spain and Lunel-Viel (Hérault), France. A contemporary chronicler records that bubonic infection devastated Spain during the first phase of the Justinianic Plague (541–544), and new interpretation of a contemporary record argues that it reached Valencia presumably before 546 (51). Further textual references, including an epitaph dating to 609, document later Iberian outbreaks (52) (See Fig. 1). In the Visigothic levels of the *Plaça de l’Almoina* in Valencia, several collective burials in an intramural cemetery were interpreted as possible plague burials (52, 55).

The historical evidence for the First Pandemic in France is more substantial, mainly based on the contemporary bishop and historian Gregory of Tours (56). He reports several plague outbreaks spanning from ca. 543 in the province of Arles through 588 in Marseille to 590 in Avignon (See Fig. 1C). The site of Lunel-Viel, around 30 km southwest of the ancient Roman city of Nîmes and less than 100 km from the mentioned cities, revealed eight exceptional inhumations in demolition trenches unrelated to the nearby contemporary cemeteries (57).

For the British Isles, the historical evidence for plague presence in the 6^th^ century is controversial. Unlike later outbreaks in 7^th^-century Britain that are reported, e.g., by Bede, the identification of a disease occurring in the 540s and called *blefed* in Irish chronicles as bubonic plague is mainly based on coincidence with the Continental European outbreaks and thus uncertain. The same is true for Britain, where a great mortality (*mortalitas magna*) is reported in the *Annales Cambriae* (see SI). For this study, we screened 22 individuals from the Anglo-Saxon cemetery of Edix Hill, well-connected to the Roman road network and Roman towns, and characterized by a number of multiple burials.

For the screening, one tooth (preferentially molar) per individual was used for every individual of a multiple burial, if available. For a number of individuals, additional teeth were tested, if sequencing the first gave a weak positive. For the collective burials from Valencia, a clear attribution to individuals was not assured, so multiple teeth were sampled per feature number, if possible. Detailed site descriptions can be found in the SI, including a table with all screened samples (Table S1).

### Sample Preparation, DNA Extraction, qPCR and MALT Screening

The sample preparation and DNA extraction for samples from Austria, France, Germany and Spain was done in the ancient DNA facilities of the ArchaeoBioCenter of the University of Munich, Germany, and the Max Planck Institute for the Science of Human History in Jena, Germany.

All teeth were cut along the cementoenamel junction and the surface of the pulp chamber was drilled out with a dental drill from the crown and in some cases the root, aiming for 30 to 50 mg of bone powder. DNA was extracted based on the protocol published in (58): The powder was suspended in 1 ml of extraction buffer (0.45 M EDTA pH 8.0, and 0.25 mg/ml Proteinase K in UV-irradiated HPLC water) and incubated at 37 °C overnight on a rotor. After centrifugation, the supernatant was mixed with 10 ml binding buffer (5 M guanidinium hydrochlorid, 40 % isopropanol and 90 mM sodium acetate) to bind the DNA on a silica column of either the MinElute purification kit (Qiagen) or the High Pure Viral Nucleic Acid Kit (Roche). After purification with washing buffer of the respective kit, the DNA was eluted in 100 µl TET buffer (10 mM Tris-HCl, 1 mM EDTA pH 8.0, 0.05 % Tween20).

All extracts were tested with the qPCR assay targeting a 52 bp region on the pPCP1 plasmid published in (18) with minor changes (0.75 mg/ml BSA, additional 5 % DMSO, EVA green instead of SYBR green, annealing for 30 s, elongation for 30 s, gradient from 60 to 90 °C). All samples showing an amplification with a melting peak between 74 and 80 °C were captured for *Y. pestis*.

The samples of Edix Hill, UK were prepared in the ancient DNA facility of the University of Cambridge, Department of Archaeology. Root portions of teeth were removed with a sterile drill wheel. These root portions were briefly brushed with 5 % w/v NaOCl using a UV-irradiated toothbrush that was soaked in 5 % w/v NaOCl for at least 1 min between samples. Roots were then soaked in 6 % w/v bleach for 5 min. Samples were rinsed twice with ddH_2_O and soaked in 70 % Ethanol for 2 min, transferred to a clean paper towel on a rack inside the glove box, UV irradiated for 50 min on each side, and then allowed to dry. They were weighed and transferred to clean, UV-irradiated 5 ml or 15 ml tubes for chemical extraction. Per 100 mg of each sample, 2 ml of EDTA Buffer (0.5 M pH 8.0) and 50 µl of Proteinase K (10 mg/ml) were added. Tubes were rocked in an incubator for 72 h at room temperature. Extracts were concentrated to 250 µl using Amplicon Ultra-15 concentrators with a 30 kDa filter. Samples were purified according to manufacturer’s instructions using the Minelute™ PCR Purification Kit with the only change that samples were incubated with 50 µl Elution Buffer at 37 °C for 10 min prior to elution.

### Library Preparation

Of putatively positive extracts in the qPCR screening, 50 µl were turned into Illumina double-stranded DNA libraries with initial USER treatment (New England Biolabs) to remove post-mortem damage in form of deaminated Cytosines by consecutive incubation with uracil-DNA-glycosylase (UDG) and endonuclease VIII(23). To enhance the efficiency of subsequent double indexing, UDG-treated libraries were quantified by qPCR using IS7/IS8 primer and split for a maximum of 2×10^10^ DNA molecules. Every library was indexed with a unique index combination in a 10-cycle amplification reaction using *Pfu Turbo Cx Hotstart DNA Polymerase* (Agilent) (59, 60). The amplification products were purified using the MinElute DNA purification kit (Qiagen) and eluted in TET (10 mM Tris-HCl, 1 mM EDTA pH 8.0, 0.05 % Tween20). For the capture, the indexed libraries were amplified to 200-300 ng/µl using *Herculase II Fusion DNA Polymerase* (Agilent) and purified a second time as described.

The non-UDG library preparation for all Edix Hill samples was conducted using a protocol modified from the manufacturer’s instructions included in the NEBNext® Library Preparation Kit for 454 (E6070S, New England Biolabs, Ipswich, MA) as detailed in (61). DNA was not fragmented and reactions were scaled to half volume, adaptors were made as described in (59) and used in a final concentration of 2.5 µM each. DNA was purified on MinElute columns (Qiagen, Germany). Libraries were amplified using the following PCR set up: 50 µl DNA library, 1x PCR buffer, 2.5 mM MgCl_2_, 1 mg/ml BSA, 0.2 µM in PE 1.0, 0.2 mM dNTP each, 0.1 U/µl HGS Taq Diamond, and 0.2 µM indexing primer. Cycling conditions were: 5’ at 94 °C, followed by 18 cycles of 30 seconds each at 94 °C, 60 °C, and 68 °C, with a final extension of 7 min at 72 °C. Amplified products were purified using MinElute columns and eluted in 35 µl EB. Samples were quantified using Quant-iT™ PicoGreen® dsDNA kit (P7589, Invitrogen™ Life Technologies) on the Synergy™ HT Multi-Mode Microplate Reader with Gen5™ software.

### In-Solution Capture

For the in-solution capture, a probe set was generated using a fragment size of 52 bp and a tiling of 1 bp with the following genomes as templates: CO92 chromosome (NC_003143.1), CO92 plasmid pMT1 (NC_003134.1), CO92 plasmid pCD1 (NC_003131.1), KIM 10 chromosome (NC_004088.1), Pestoides F chromosome (NC_009381.1) and *Y. pseudotuberculosis* IP 32953 chromosome (NC_006155.1). The capture was performed as previously described (62) on 96-well plates with a maximum of two samples pooled per well and all blanks with unique index combinations in one well.

### Sequencing and Data Processing

All captured products were sequenced either on a Illumina NextSeq500 or HiSeq4000 platform at the Max Planck Institute for the Science of Human History in Jena, Germany. Edix Hill libraries were sequenced on Illumina NextSeq500 at the University of Cambridge Biochemistry DNA Sequencing Facility and the FastQ files were processed on the Estonian Biocenter server. De-multiplexed reads were processed with the EAGER pipeline (63) starting Illumina adapter removal, sequencing quality filtering (minimum base quality of 20) and length filtering (minimum length of 30 bp). Sequencing data of paired end and single end sequencing were concatenated after adapter removal and merging. The same was done for samples from the same individual (DIT004) and all data from Lunel-Viel (LVC) due to low genomic coverage. The sequencing results are shown in Table S3.

All Edix Hill samples were screened using MALT (19) against a reference set including full bacterial and viral genomes with 85 % identity, the strong positive sample EDI001.A was then sequenced deeper to a coverage of 9.1-fold without enrichment.

After clipping of 3 bases on each end with fastx_trimmer of the FASTX toolkit (https://github.com/agordon/fastx_toolkit) to remove the majority of damaged sites for the non-UDG library of EDI001.A, the sample was processed along the UDG treated libraries. Mapping against reference genomes of CO92 (chromosome NC_003143.1, plasmid pMT1 NC_003134.1, plasmid pCD1 NC_003131.1, plasmid pPCP1 NC_003132.1) was done with BWA using stringent parameters (-n 0.1, −1 32). Reads with low mapping quality were removed with Samtools (-q 37) and duplicates were removed with MarkDuplicates. For the plasmids, a merged reference was used, consisting of the CO92 reference of pCD1 (NC_003131.1), pMT1 (NC_003134.1) and pPCP1 (NC_003132.1, with base pairs 3,000 to 4,200 masked (18)), to avoid overestimation of coverage due to homologous regions. For the verification of positive qPCR results, we normalized the number of reads mapping to each plasmid with reads mapping to the chromosome and calculated the Mahalanobis distance for each sample to detect outliers. Based on this, we excluded the samples PEI001.A and DIR002.A as false positives (Table S2). The raw data of the Aschheim and Altenerding genomes were processed identically, however considering only the A120 sample for Aschheim (6, 7).

### SNP Calling and Evaluation

All genomes recovered from UDG-libraries with higher than 5-fold mean coverage including the Altenerding genome were assessed in the SNP analysis. Additionally, the sample WAG001.A was evaluated to explore its phylogenetic position, since it was the only positive sample of the relevant site.

The UnifiedGenotyper within the Genome Analysis Toolkit was used for SNP calling and creating VCF files for all genomes, using ‘EMIT_ALL_SITES’ to generate calls for all positions in the reference genome. For the subsequent analyses, 233 previously published modern *Y. pestis* genomes (Table S8), one genome from 2^nd^ to 3^rd^ century Tian-Shan mountains (DA101 (26)) one genome representing the Black Death from London East Smithfield (8291-11972-8124 (12)), and six Second Pandemic genomes (Ellwangen, Marseille L’Observance OBS107, OBS110, OBS116, OBS124, OBS137 (11, 12)) were taken along together with *Y. pseudotuberculosis* (IP32953) as an outgroup. Previously identified problematic regions (24, 38) as well as regions annotated as repeat regions, rRNAs, tRNAs and tmRNAs were excluded for all following analyses. MultiVCFAnalyzer v0.85 (64) was used for generating a SNP table with the following settings: Minimal coverage for base call of 3 with a minimum genotyping quality of 30 for homozygous positions, minimum support of 90 % for calling the dominant nucleotide in a ‘heterozygous’ position. All positions failing these criteria would be called ‘N’ in the SNP table. For the SNP evaluation, all ‘N’ positions of unique SNPs within the First Pandemic lineage were re-evaluated, replacing ‘N’ by ‘0’ for not covered and lower case letters for homozygous positions with max. 2-fold coverage. To test for possible mixed infections of elevated contamination, all SNPs not passing the 90 % threshold were plotted (Fig. S1).

For the evaluation of unique and shared SNPs of First Pandemic genomes retrieved from non-UDG libraries, we used the newly developed tool ‘SNPEvaluation’ (https://github.com/andreasKroepelin/SNP_Evaluation) and a comparative mapping, using BWA with high stringent (-n 0.1, −1 32) and low stringent (-n 0.01, −1 32) mapping parameters, allowing for more mismatches in the latter. SNPs were called true positive when meeting the following criteria within a 50 bp window: (A) the ratio of mean coverage of low stringent to high stringent mapping is not higher than 1, (B) no ‘heterozygous’ positions, (C) no non-covered positions (Table S5). SNP evaluation on the plasmids was done using the same criteria after mapping to the individual references as described above. For the SNP effect analysis, the remaining unique true SNPs were compared to the genome annotations of the CO92 *Y. pestis* reference genome (Table S6).

Shared SNPs (Table S7) were evaluated with the same criteria with minor modifications: The minimum threshold for calling a position was set to 1 read covering and SNPs were called true positive, if the SNP passed the criteria in more than half of the genomes under examination.

The genome of EDI001.A from Edix Hill was only included in the presented SNP evaluation for comparison, since it is derived from a non-UDG library and damaged sites interfere with both the comparative mapping and the count of heterozygous positions.

The Aschheim genome was evaluated separately (Table S8) but with the same criteria. As previously addressed (6), the enormously high number of false positive SNPs might not be explained solely by contamination by soil bacteria or sequencing errors but additionally by PCR or capture artefacts.

### Phylogenetic Analyses

For the phylogenetic analyses we aimed for one high coverage genome per site to minimize missing data in the SNP alignment, excluding the genome of UNT004.A after assuring no conflicting positions with UNT003.A in the SNP evaluation. A maximum likelihood tree (RAxML 8 (28) using the GTR substitution model, Fig. 2A, for full tree see Fig. S2) was generated without exclusion of missing and ambiguous data (full SNP alignment), resulting in a total number of 6496 SNPs. Robustness of all trees was tested by the bootstrap methods using 1000 pseudo-replicates.

A detailed tree of the First Pandemic lineage was drawn manually based on the performed SNP evaluation, excluding all false positive SNPs (Fig. 2B).

### Analysis of virulence factors and genome decay

The presence/absence analysis for genes was performed with BEDTools (65) by calculating the percentage across each gene (3). Since gene duplications can affect the mapping quality, the mapping quality filter of BWA was set to 0 (-q = 0) to generate a bam-file as input. For the heatmap of virulence factors (Fig. 3), a collection of proven and putative virulence genes (29, 30) was evaluated. The more extensive analysis on genome decay was based on the annotation file for the reference genome CO92 (54) by extracting all regions annotated as ‘gene’.

For the exact determination of the start and end positions of deletions, mapping with BWA_MEM was performed (66).

### Radiocarbon Dating

At least one individual per burial was sampled for radiocarbon dating for all burials that tested positive for *Y. pestis*, assuming simultaneity of interment for the multiple burials. Samples were dated at the CEZ Archaeometry gGmbH, Mannheim, Germany. The raw radiocarbon dates were calibrated with IntCal13 (67) in OxCal v4.3.2 (68). All raw and calibrated dates are given in Table S9; Fig. S5 shows the respective probability distributions. Some of the intervals completely pre-date the onset of Justinianic Plague (541) which could be explained by a marine or freshwater reservoir effect (69, 70) or human bone collagen offset (71). In the absence of C/N isotope data and a well-established method for addressing the human bone collagen offset, we report calibrated dates without any correction.

### Cartography

All maps were generated in ArcGIS 10.4.1 (ESRI) using the ‘World Ocean Basemap’ without references. The sources for all historical occurrences are given in the SI. The mapped regions in Fig. 1 are primarily based on the Digital Atlas of Roman and Medieval Civilizations (DARMC; https://darmc.harvard.edu) maps “Provinces AD303-324” for the western Europe and “Provinces ca. AD500” for eastern Europe, Middle East and Africa. The provinces in Fig. S6 are based on a georeferenced map by Rettner and Steidl (72), the Roman roads are combined from Rettner and Steidl and the DARMC map “Roman Roads”. The main rivers in Figs. 1C and 3 were taken from Natural Earth (ne_10m_river_lake_centerlines, http://www.naturalearthdata.com), based on data provided by the European Commission, Joint Research Centre, Institute for Environment and Sustainability (JRC IES).

## Supporting information

## Author contributions

M.McC., M.H., A.H. and J.K. designed the study. M.K., M.A.S, C.L.S., K.N., G.U.N. and J.S.B. performed laboratory work. A.K. developed the new analytical tool. M.K. and M.A.S. performed data analyses. B.T. and S.I. performed anthropological examination. C.L.S. and T.K. provided genomic data. B.H.-G., B.P., J.H., A.R.iL., C.R., P.S., J.P., J.E.R., and M.H. identified and provided access to archaeological material. B.H.-G., B.P., J.H., A.R.iL., C.R., C.C., P.S. and M.McC. provided archaeological and historical information. M.K., M.A.S, M.McC. and A.H. wrote the paper with contributions from all authors.

## Acknowledgements

We are grateful to Aditya K. Lankapalli, Aida Andrades Valtueña and all members of the Department of Archaeogenetics Max Planck Institute for the Science of Human History for support and fruitful discussions, and Raphaela Stahl, Marta Burri, Cäcilia Freund, Franziska Aron, Antje Wissgott and Guido Brandt for their assistance in the lab. We thank the staff of the SAPM for support during sample collection and Ronny Friedrich at the CEZ Archaeometry gGmbH, Mannheim for providing additional information on radiocarbon dates. Furthermore we thank Henry Gruber for his correspondence. This study was supported by the European Research Council starting grant APGREID (to J.K.) and by the Justinianic Pandemic Working Group at MHAAM/SoHP.

## Data availability

The raw sequencing data of the relevant plague positive samples will be available on the European Nucleotide Archive under project accession number PRJEB29991 upon publication.

