## Supplementary material for "Ancient *Yersinia pestis* genomes from across Western Europe reveal early diversification during the First Pandemic (541–750)"

**This PDF file includes:**

Supplementary Information text
- SNP Evaluation of the Aschheim Genome and SNP Effect analysis
- Phylogeographical Analysis
- Archaeological Context Information
- Sources for mapping plague outbreaks between 541 and 650 CE

Figs. S1 to S6
Tables S1 to S9
References for SI reference citations

### Supplementary Information Text

#### SNP Evaluation of the Aschheim Genome and SNP Effect Analysis

The Aschheim genome was evaluated separately, given its peculiarly high number of false positive SNPs described previously (1). Our systematic evaluation verified previous classifications: all SNPs potentially unique to Aschheim that passed the criteria show a coverage lower than 5-fold, which was the threshold of their SNP calling. However, the high number of presumably shared SNPs that did not pass our stricter criteria underlines again the high 'heterozygosity' of the genome (see Table S8) that might be explained not only by contamination by soil bacteria or sequencing errors but presumably also by PCR and capture artefacts, as previously discussed (1). Therefore, the Aschheim genome was excluded from subsequent analyses.

Of the 17 unique chromosomal SNPs that were detected among all new genomes, 10 are non-synonymous in coding regions of (hypothetical) proteins (Table S6). The genome of VAL001.B shows non-synonymous SNPs in the genes *tyrP*, YPO1985 and YPO2588. TyrP is a transcriptional regulator for the metabolism of aromatic amino acids and was identified as a virulence factor crucial for the infection of mice (2). YPO1985 was identified as a glycosyl transferase gene inactivated in the avirulent strain 91001 and thus might be as well a virulence factor (3). The gene YPO2588 codes for an ABC transport protein. An additional non-synonymous SNP was detected on the pMT1 plasmid in the putative DNA-binding protein YPMT1.59C. For the genome PET004.A only one non-synonymous SNP was identified located on the hypothetical protein YPO3510. The genome of LVC_merged shows six non-synonymous SNPs: in the genes *marC*, a multidrug resistance protein; *phrB,* coding a 3',5'-cyclic-nucleotide phosphodiesterase; *tyrA*, a bifunctional chorismite mutase/prephenate dehydrogenase; the phosphoenolpyruvate carboxylase *ppc* and the two hypothetical proteins YPO2238 and YPO4112. The gene *ppc* has been shown to be connected with the type three secretion system that is essential for pathogenicity in *Y. pestis* by injection of Yops (*Yersinia* outer proteins) into host cells of the innate immune system (4).

#### Phylogeographic Analyses

The new genomes and radiocarbon dates combined suggest an association of the British genome as well as the polytomy giving rise to the four lineages with the early phase of the First Pandemic or even the Justinianic Plague itself (541-544). The accumulation of one (EDI001) or two (Altenerding cluster) SNPs from the basal node of all genomes could have happened on the way from Egypt to western Europe. The fact that the pandemic reportedly spread from Pelusium along the Mediterranean coastline in two independent waves, one heading west to Alexandria and the other east to Palestine, could explain the early branching event (5). Strikingly, such a diversification during the onset of a pandemic has not been found yet for the Black Death (1348-1352), where the two genomes from London East Smithfield (6) and Barcelona were found to be identical (7). Besides differing mutation rates, this might be due to differences in propagation speed between the 6th and 14th century, related to changes in human mobility by land and sea: a significantly slower or less direct transmission over large distances would allow the pathogen to acquire more substitutions.

The lineages found in Bavaria could have spread there by a ‘western route’ from Gaul, by a ‘southern route’ from Italy or by an ‘eastern route’ from Illyricum, which were all affected by plague in or around 543. The presence of plague in the British Isles even suggests a fourth ‘northern route’ upstream along the Rhine river. The ‘western’ and ‘southern route’ would have necessitated overland transport via the Roman road network that connected all of the relevant sites with the Mediterranean coastlines and was still functional in the 6th century (Fig. S6). The ‘southern route’ would have required crossing the Alps via different passes that had been used since Antiquity (8). Navigation along the Danube could have facilitated the ‘eastern route’. The importance of rivers for the spread of plague has already been shown exemplarily for the Rhône during the First Pandemic (9) and for the Black Death (10). However, attempts to prove the preferential spread via rivers during Second Pandemic have recently been criticized (11, 12).

The site of Petting is geographically situated only 100 km southeast of Aschheim and Altenerding (Fig. S6). However, it was located in the Roman province *Noricum ripense* whereas the sites with the distinct uniform lineage were situated in *Raetia secunda* (Aschheim, Altenerding, Unterthürheim) or close by (Dittenheim). Although the administrative system of the Western Roman Empire had broken down by the mid-6th century, its political borders continued to be influential, not least because of the ecclesiastical system of dioceses that followed them. It is possible that these ancient boundaries influenced the spread of the two epidemic outbreaks in modern-day Bavaria. Since the river Inn separated *Raetia secunda* and *Noricum ripense*, this might suggest that rivers could serve as physical barriers to the spread of plague where river transport was negligible. This in turn would rather suggest the ‘eastern route’ or the ‘southern route’ for Petting as described above.

Complementing the previous results from Aschheim and Altenerding, our new data from Unterthürheim and Dittenheim underline the epidemic extent of this plague outbreak in early medieval Bavaria, totalling 16 individuals with genomic evidence for *Y. pestis* and an additional five PCR-positive individuals in Aschheim (13). Far from the urban centres of the time and any recorded outbreak of plague, the new molecular evidence stands in strong contrast to Durliat’s claim that the Justinianic Plague was merely an urban phenomenon. Instead, we view this data as being in line with ancient statements by Procopius, John of Ephesos and Paul the Deacon who reported that the countryside of the Levant and Italy were severely impacted.

**Archaeological Context Information**

*The following site descriptions present condensed information on all sites examined in this study. The classification of multiple burials follows McCormick (2015). Sex and age determination is based only on morphological examination. For the age classification, the German system* (14) *is used as follows: Infans I (0-6 years old), infans II (7-12 years old), juvenile (13-20 years old), adult (20-40 years old), mature (40-60 years old), senile (more than 60 years old).*

<u>Alladorf (ALL; Markt Turnau, Landkreis Kulmbach, Germany):</u>
The Carolingian cemetery of Alladorf, dating roughly between 630 to 720, revealed 163 graves with remains of 276 individuals. However, the total size of the cemetery is unknown, since the excavation did not reach the borders of the burial area. Leinthaler classified three burials as double burials (type 1): 179/180 (ALL001, early adult female; ALL002, infans I), 184/185 (ALL003, infans II; ALL004, late adult to early mature male) and 203/204 (ALL005, late adult male; ALL006, infans I). The two burials 188/189 (infans I; late adult to early mature male) and 208/209 (infans I; early adult female) were classified as double burials with unclear simultaneity.
(15, 16)

<u>Dirlewang (DIR; Landkreis Unterallgäu, Germany):</u>
The Alemannic site of Dirlewang is a very small cemetery with 40 excavated burials and an expected number of 55 burials in total. It dates to the Late Merovingian period, from 650-700, based on the archaeological finds. Two double burials (type 1) were found on this site, 33/34 (DIR001, juvenile to early adult female; juvenile male) and 38/39 (adult male; DIR002, adult female). Graves 18 and 19 (mature male; early adult male) did not share the same grave pit but were buried very close to each other, indicating a connection. Burials 30/31, 36/37 and 2 were classified as non-simultaneous successive or additive double burials (type 2).
(17)

<u>Dittenheim (DIT; Landkreis Weißenburg-Gunzenhausen, Germany):</u>
The early medieval cemetery of Dittenheim recruited from a settlement on the site of the modern village. Only 6.5 km south of the *limes*, the settlement was probably well connected to the remaining Roman infrastructure. 2.5 km Southeast of the cemetery, remains of a Germanic fortification dating to the Migration Period were found, known as *Gelbe Bürg*. This structure went out of use around 500, just before the region fell under the rule of Franks.
The cemetery was known already by 1937. The first excavation campaign in 1968 revealed the first 114 burials. Later campaigns took place in 1971 (burials 115-164) and 1972 (165-244). The excavations probably reached the borders of the cemetery. However, some burials might have been lost due to erosion from a nearby river and from plowing. Besides the early medieval burials, older settlement traces from the Linear Pottery culture, Pre-Roman Iron Age and the imperial Roman period as well as one Bronze Age burial were found on the site.

The cemetery of Dittenheim, dating from the middle of the 6th century to the end of the 7th century, revealed 238 graves containing 244 individuals and 10 cremations from the same period. Other remarkable finds were three horse burials and one circular pit. Four burials were classified as double burials type 1: 8A/B (DIT007, mature female; mature male; possibly an additive burial), 18A/B (DIT003, mature male; DIT004, adult female), 22A/B (DIT005, infans I; DIT006, adult female) and 188A/B (DIT001; DIT002, both infans II). There were also three exceptional single burials in prone position (180, 201, 204) deviating from the contemporary burial rites.

(18)

<u>Edix Hill (EDI; civil parish Barrington, Cambridgeshire, United Kingdom):</u>

The Anglo-Saxon cemetery of Edix Hill, close to Barrington and Orwell was initially discovered in the 19th century. Excavations between 1989 and 1991 revealed part of an inhumation cemetery comprising 149 individuals in 115 graves, dating to between 500 and 625/650, although one burial was believed to be possibly Iron Age. It is estimated that there may originally have been around 300 burials in total with a complete cross-section of the population by age and sex suggesting that the burials relate to a community of 50–65 people spanning around 150 years. The initial dating of the cemetery was based primarily upon artefact typologies of the various grave goods and seriation by correspondence analysis, with burials broadly divided into earlier and later groups and some evidence for spatial patterning over time. Subsequent radiocarbon dating has broadly confirmed and refined the dating of the cemetery. Palaeopathological evidence suggested the presence of tuberculosis, leprosy and cancers. The human remains are currently held by Cambridgeshire County Council, who generously provided access to the material.

The cemetery contained a total of 18 multiple burials, of which 10 were classifiable as type 1: they comprise one quadruple burial with at least two of the individuals buried simultaneously and eight simultaneous double burials. Four double burials are of unclear simultaneity, another four and a triple burial are clearly understandable as type 2 (non-simultaneous). The site was initially not sampled for the purpose of plague screening, so samples from both multiple and single burials were screened. This includes the single graves 76 (Sk405, EDI001; juvenile), 69 (Sk359, EDI002, early adult female), 78 (Sk424, EDI003, juvenile), 46 (Sk146, EDI010, early mature male), 60 (Sk183, EDI011, adult female), 63 (Sk198, EDI012, early adult male), 83 (Sk436, EDI014, early adult female), 90 (Sk458, EDI017, senile female), 95 (Sk530, EDI018, late mature female), 97 (Sk551, EDI019, early adult male), 99 (Sk576, EDI020, mature male), 100 (Sk578, EDI021, early mature male), 105 (Sk592, EDI022, late mature female), the non-simultaneous double burial grave 66 (Sk322A, EDI013, at least adult male; SK322B, early adult female) and triple burial grave 18 (Sk42A1, infans I; Sk42A2, early adult with indet. sex); Sk42B, EDI009, early adult female), the simultaneous double burials grave 96 (Sk547A, adult female; Sk537B, EDI004, infans II), 106 (Sk626A, EDI005, early adult female; Sk626B, EDI006, early adult male), 9 (Sk13A, EDI007, mature to senile male; juvenile male) and 84 (Sk440A, EDI015, adult female; Sk440B, infans I) as well as the complex burial grave 2, with two individuals buried simultaneously (Sk3B, adult to mature male; Sk3C, EDI007, early adult

male) and two more individuals buried later (Sk3A1 and Sk3A2, both adult and possibly female and male respectively).
(19, 20)

Forchheim (FOR; Gemeinde Pförring, Landkreis Eichstätt, Germany):
Within the vestiges of a late medieval settlement, a quadruple burial was found without any context suggesting a larger burial ground. Two mature women (FOR002, individual 2), a mature man (FOR003, ind. 3) and a male juvenile (FOR001, ind. 1) were buried partially overlapping but all in East-West orientation. Remarkably, one of the women (2) was buried in the prone position. Grave goods such as a sax and a glass bead necklace date the burial to the second half of the 7th century. A belt buckle was indicative of clothed inhumation.
(21)

Grafendobrach (GRA; Gemeinde Kulmbach, Landkreis Kulmbach, Germany):
The Carolingian cemetery of Grafendobrach revealed 85 burials; the excavations 1975-1976 did not reach the edges of the burial ground. The various garment artefacts found in the graves point to the late 9th to early 10th century. Besides several secondary burials (type 2), there were two remarkable graves: complex 83/84/85 consisted of two adjacent stone cists holding two women (83, GRA002, early mature; 85, GRA003 early adult) and an infant (84, ca. 1 year old) buried at the feet of 83. A single burial of a man (42, GRA001, early senile) was covered with stones, on top of which two neonates (43, 44) were found without individual burial pits.
(22)

Kleinlangheim (KLH, Gemeinde Großlangheim, Landkreis Kitzingen, Germany):
The Frankish cemetery of Kleinlangheim dates to the late 5th to early 8th century and was completely excavated from 1962 to 1969. It contained 244 burials and a remarkable number of 56 cremations; unlike other contemporary cemeteries, the graves showed signs of grouping, potentially indicating family groups. Nine multiple burials type 1 were identified, eight of them double burials: 35/36 (KLE005, adult male; late mature male), 41/42 (late adult male with skull fracture; juvenile to early adult), 147/148 (min. adult; neonate), 171/172 (two infans I), 208/209 (infans I; early adult female), 272/273 (late adult male; KLE004, senile male), 211 (KLE001, mature male; second individual cremated) and 218/219 (KLE002, mid-adult to mature male, KLE003, late adult to early mature female). The triple burial 100-102 contained remains of two infants (1-2 years old; 4-6 years old) and one mature individual of indeterminable sex.
(23)

Leobersdorf (LEO, Bezirk Baden, Austria):
The Avar cemetery of Leobersdorf was excavated between 1977 and 1983 and was dated to 640-800. In total, 154 burials containing remains of 171 individuals were excavated with an exceptionally high number of 25 multiple burials. 16 burials were identified as double burials, of which 9 contained each an adult and a subadult (16: adult female and infans I; 57: mature male and infans I; 74: adult female and infant; 93: mature male and infans I; 100: mature male

and juvenile female; 104: adult female and infans I; 114: adult male and infans I; 119: late mature male and infans II; 140: adult female and infans II). Four consisted of two adults (35: mature male and female; 86: adult female and mature male; 114: adult female and male; 144: senile and adult male) and two held only subadults (23: male juvenile and infans II; 103: two juvenile females; 145: infans I and II). The remains of infant individuals often were identified only after the morphological examination following the excavation.

Three burials were identified as triple burials (21: senile male, mature female and infant; 67: adult female and two infants; 82: juvenile male, LEO001; adult female, LEO002; infans I, LEO003). A fourth (99) was interpreted as a double burial (mature male and infant) with a secondary burial (female).

Two burials contained four individuals (105: mature female, juvenile female, infans I and neonates; 134: adult female, mature male, infans I and II). Burial 79 contained the remains of five individuals, identified as the burial of a mature female with secondary burials of four additional individuals (senile female and male, juvenile female and infans II) that had probably been buried on the same spot previously, and were reburied after the mature female.

(24, 25)

Lunel-Viel (LVH, LVC; Arrondissement Montpellier, Département Hérault, France):

The site of Lunel-Viel is equidistant (25 km) from the modern and Roman towns of Montpellier and Nîmes at the intersection of two paved roads that were active in late antiquity: a later Roman secondary road running about 2 km southeast and parallel to the Roman *Via Domitia* that connected Iberia to Italy, and a road connecting the *Via Domitia* to the coastal lagoon *l'Étang d'or*, part of the complex of lagoons that included that of Lattes (anc. *Lattara*), famous already in antiquity for its fishery (Pliny the Elder, Natural History, 9.29-32). A brief Roman-era occupation in the 2nd century BCE was followed by continuous settlement beginning ca. 50-80 CE, although the ancient settlement area was abandoned in the 7th century, and the exact location of the new dwelling zones between the 7th and the 10th century remains unknown. Three inhumation cemeteries received the deceased of this community in succession from the 4th century. The earliest, at *Le Verdier,* functioned from the end of the 3rd century down to the beginning of the 6th century, and yielded 340 burials. The second, known as *Les Horts*, received burials of the late 5th to 7th centuries; 140 burials were excavated out of an estimated original total of ca. 200. The third is associated with the church of St. Vincent; burials began there ca. 520 and continued down to the 17th century; 97 have been excavated.

The cemetery *Les Horts* contained one clear case of a double burial, likely simultaneous (type 1), of two adult individuals (38A/B) placed head facing toes in a sarcophagus; the head and upper body of 38B were destroyed, along with the feet of 38A and part of the sarcophagus, during later leveling operations. Disarticulated remains of a third poorly preserved adult (38C, LVH001) were found pushed into the eastern end of the sarcophagus. Grave goods likely stemming from the original burial (one buckle, one plated buckle, one fibula, one pin and one clasp) date the burial of 38C to 475–550.

Outside the cemeteries, anomalous (deviant) burials of the remains of eight individuals were found during the excavation of the Gallo-Roman structures, called the *Quartier central.*

Robbing of foundation stones in antiquity had left empty trenches which were subsequently used for the summary interment of these individuals. A pin buckle, a spindle whorl and a knife found with the remains of the presumably clothed individuals date the interment to the 5th to 6th century. The positions and postures of the individuals clearly deviate from the burial customs seen in the contemporaneous cemeteries of *Les Horts* and *Le Verdier*: two adult women (3A: LVC003, adult; 3B: LVC004, mature) were placed under limestone slabs, with one resting her head on the thighs of the other. Westward of this group, a woman (1: LVC001, early adult) and another adult individual (2: LVC002, late adult to early mature, sex indeterminable) were found; the latter however had been disturbed by later agricultural activities. Whereas these individuals appear to have been laid rather carefully on the ground, the two men adjacent to the west (4: LVC005, mature, 5: LVC006, early adult) seem to have been carelessly dropped into the trench, the latter with his arms stretched over his head. Two more individuals, a young adolescent woman (6, LVC007) and an infant (7, sex indeterminable), were found in a second trench to the east. However, all individuals were interred more or less in supine East-West oriented position, as expected for early medieval Christian burials.
(26)

München-Aubing (AUB, Stadt München, Germany):
The Baiuvarian cemetery of München-Aubing was excavated in two phases, 1938 and 1960-1963. It was used from the 5th to 7th century. A total of 896 graves was excavated. Four double burials (type 1) have been identified: 854/855 (AUB008, adult male; late mature male) 809/810 (AUB006; AUB007, two adult males), 724/725 (AUB004, adult male; AUB005, mature male; with possible later manipulation) and 676/677 (AUB002, mature female; AUB003, male individual). A third individual (675, infans I) was buried later on top of the double burial 676/677.
(27)

Neuburg an der Donau (NEU; Landkreis Neuburg-Schrobenhausen, Germany):
The site of Neuburg an der Donau was occupied at least from the first half of the 2nd century by a Roman military base associated with an urn cemetery with around 130 cremations. The burial ground *Seminargarten* is related to a later occupation period during the 4th century. From an estimated number of 150 burials, this site revealed 130 burials with 133 individuals. One burial was identified as a double burial (32A/B, adult female and juvenile with indeterminable sex), one as a triple burial (34A/B/C: mature male; NEU001, mature male; NEU002, adult male). Both are part of Zone 1, dating to 330-360. Two additional burials (14, 28) were suggested to contain “mother and child” (neonate/infant).
(28, 29)

Peigen (PEI; Markt Pilsting, Landkreis Dingolfing-Landau, Germany):
The early medieval cemetery of Peigen dating to the mid-5th to the 7th century revealed around 274 burials. Among these, three could be identified as double burials. Double burial 18

contained an adult woman (18-1, PEI001) and an early adult woman (18-2, PEI002), buried in opposite orientation. Double burial 63 contained an adult woman (63-1, PEI004) and a senile man (63-2, PEI005). The burial of a late mature male (109-1) and a child (infans I, 109-2) form the third clear double burial. The burials 62-1 of a juvenile male and 62-2 of a late mature female (PEI003) were adjacent but at different depth and therefore of questionable simultaneity (type 2).
(30)

Petting (PET; Landkreis Traunstein, Germany):
Although located in Germany today, Petting falls in a region that was in ancient times associated to the Roman city and later ecclesiastical center of *Iuvavum* (today Salzburg, Austria) and has therefore a different settlement history than other parts of southern Bavaria. The cemetery of Petting was discovered in 1991 and excavated within the following two years. A comprehensive publication of the site of Petting is still pending, not least because the archaeological findings were not available for research until the ownership situation was clarified in 2010. Studies so far concentrated on individual burials or specific artefacts.
The cemetery was completely excavated, counting 721 burials. Only 24 burials were lost due to destruction prior to the excavation: around 50 % of the burials fell victim to ancient grave robbers. At least four multiple burials were observed in the cemetery, all presumably type 1. The double burial 172/173 contained a female and a male individual, the double burial 377/378 a late adult to mature and a late adult to early mature male (PET003/PET004), the triple burial 342/343/344 at least one early adult male (PET001) and an infans II (PET002). The double burial 630/631 of an early adult female (PET006) and an early mature female (PET007) was associated with a single burial of an infans II (PET005) on top.
(31, 32)

Regensburg Fritz-Fend-Straße (RFF; Stadt Regensburg, Germany):
Regensburg, *Castra Regina,* was an important Roman military base with a civilian settlement of the province *Raetia*. The extensive adjacent cemetery *Kumpfmühl* was excavated in several campaigns following the first excavation in 1872 by Dahlem. He documented 827 burials and cremations but left the vast majority of features undocumented, which led Schnurbein to the assumption that the first excavation uncovered around 3000 cremations and 2000 burials, mainly dating to the late 3$^{rd}$ to mid 4$^{th}$ century. More features were lost in this area due to later construction works without archaeological survey. An excavation in 1999 revealed 100 additional cremations and 50 burials. The excavations of *Fritz-Fend-Straße* and *Im Güterbahnhof* took place in 2011 and revealed 48 cremations/115 burials and 161 cremations/116 burials, respectively. The site *Fritz-Fend-Straße* contained two exceptional double burials of type 1: In burial 30, the second individual (30-1: RFF001, late adult to early mature male) was laid in prone position and opposite orientation on top of the first (30-2), who was buried in supine position. In 53, the two individuals, a juvenile female (53-2: RFF002) and an early mature male (53-3, RFF003), were buried in crouched position facing in the same

direction but in opposite orientation. An archaeological or radiocarbon date for *Fritz-Fend-Straße* has not yet been published, but a similar date as for *Kumpfmühl* can be assumed. (33)

Sindelsdorf (SIN; Landkreis Weilheim-Schongau, Germany):

The early medieval cemetery of Sindelsdorf consists of 331 burials with remains of 354 individuals dating from 500 to 720. It included three double burials type 1 (25/26 senile male and mature male; 51/52, senile and mature male (SIN006, SIN007); 154/155, infant and senile female) and a burial group of three individuals with one buried later on top (163: infant; 164: SIN002, late adult to early mature male; 165: SIN003, adult female; 162: SIN001, senile male). The double burial 25/26 with a later burial on top (24, mature female) may be identified as a probable homicide due to a perimortem skull fracture (26). (34)

Straubing Azlburg I/II (SAZ; Stadt Straubing, Germany):

Straubing, called *Sorviodurum* by the Romans, was an important *castrum* on the Danubian border of *Raetia*. The Late Roman cemeteries – around 200 m apart from each other – were excavated in 1981 (Azlburg I) and 1984 (Azlburg II). However, the borders of the cemeteries were not reached on all sides, so the original dimensions remain unclear. They were in use at roughly the same time, approximately between 300 and 450. Azlburg I contained 107 graves with 111 individuals and one cremation. The burial 54-1/-2/-3 was identified as a triple burial (SAZ002, male juvenile; SAZ003, male juvenile; third individual min. adult). The double burial 16/17 contained the remains of two children (SAZ001, infans I; neonate). Azlburg II contained additional 434 graves with 45 individuals. The double burial 5a/b of two adult men was interpreted as the burial of two soldiers. (35)

Unterthürheim (UNT; Gemeinde Buttenwiesen, Landkreis Dillingen an der Donau, Germany):

The town of Unterthürheim is probably the site of the early medieval settlement that used the burial ground *Buttenwiesen*. Traces of an older Imperial Roman settlement (1$^{st}$ to 5$^{th}$ century) were found on the *Thürlesberg* hill, around two kilometers from the cemetery; it was near the Roman roads *Via Iuxta Danuvium* (following the Danube) and the *Via Claudia Augusta* (connecting *Augusta Vindelicum*/Augsburg with Italy) which intersect at the fort *Submuntorium*, around 10 km Northeast of Unterthürheim.

The Alemannic cemetery has been known since 1889, when a local resident discovered the first six graves. Additional burials were found between 1943 and 1966. The first professional excavation took place in 1968 (graves 1-42), later excavations occurred between 1969 and 1972 (graves 43-178) and in 1979 (graves 180-238). All of them were rescue excavations, which explains the scattered trenches and limited dimensions: The excavations are thought to have reached the southern and northern border of the cemetery, but the extent to the West and East remain unknown. In total, 256 burials were excavated of which 230 were preserved well enough for more detailed examination. They are archaeologically dated to 525 to 680.

Lüdemann lists in total 14 double burials, two triple burials and one quadruple burial. The triple burials 120/122 and 124/125 as well as the double burials 167/168, 186/186a and 217/217a were classified as secondary burials (type 2). Burials 12 (mature with indeterminable sex, subadult) 63/64 (UNT006, adult female; UNT007, infans I), 79/80 (adult female, infans I), 116 (mature female; UNT001, infans II), 140/141 (mature male, early adult female), 145/146 (adult to early mature female, male) and 189 (adult females, neonates) presumably simultaneous double burials (type 1). 65/66 (infans I, infans II) and 190/190a (adult to early mature female, early adult male) were classified by Grünewald as possible secondary burials (type 2). For the burials 97/98 and 147/149, the classification is unclear.

The quadruple burial 131-134 (UNT004, infans I; UNT005, adult male; infans I; min. adult male) was associated with two single burials: 129 (UNT002, infans I) and 130 (UNT003, mature male).

(25, 36)

<u>Valencia Plaça de l'Almoina (VAL; Ciudad de Valencia, Spain):</u>

The site of Valencia, "Plaça de l'Almoina" is an intramural cemetery that was founded in the 5th century and remained in use during the Visigothic period in the 6th and 7th centuries. It was excavated from 1985 to 1999. Contrary to the earlier Roman practice, the necropolis was not located outside the city walls but close to a shrine commemorating the martyrdom of St. Vincent, and adjacent to the cathedral. Many burials still follow the later Roman tradition that made use of *tegulae* and amphoras. Among the later burials, there are several monumentalizing collective tombs that were probably used as elite family tombs for successive burials.

The excavated areas of the necropolis contained a number of multiple burials of differing types and dates, including 15 or 16 collective slab tombs that appear to reflect successive burials (type 2: details in McCormick 2016, 1024n15). At the moment of sampling, the human remains were comingled and only sorted by find numbers (corresponding the numbers VAL001-009), so the assignment to individuals is not possible for this site. Four or five features seemed to be of type 1, i.e., reflecting at least in part simultaneous burials. The quadruple burial tomb 4 was covered with *tegulae* and contained four individuals who were piled one on top of another in an east-west orientation (two samples were taken, VAL008). Tomb 28 was stone-lined and just north of an apse identified as a memorial shrine to the martyrdom of St. Vincent; it contained a minimum of 21 individuals, from whom sixteen samples were taken (VAL005, VAL006, VAL007). Tomb 50, just southwest of Tomb 28 (and actually under the wall of that apse), may well have been a continuation of Tomb 28; it contained at least seven individuals, from whom seven samples were taken (VAL003, VAL004). Tombs 28 and 50 are lower than and earlier than the wall of the apse. The apse itself seems well dated to the late sixth or early seventh century, based on pottery excavated in a pit under the apse pavement. Multiple tomb 40 was a pit burial containing four supine individuals (five samples, VAL002) oriented east-west in the entry from the east into the apse of the memorial shrine. Tomb 41 is south of the apse structure which might indicate a privileged burial space. It is a pit that seems to have used pre-existing Roman walls to the south and east for two sides; bricks were simply placed without mortar to form the other two sides of the burial space, which might suggest a hasty or improvised burial.

It contained the very disturbed remains of at least 15 individuals, including 4 subadults from which we took seven samples (VAL001).
(25, 37–39)

Waging (WAG; Landkreis Traunstein, Germany):
The early medieval cemetery of Waging, close to the site of Petting, was discovered and excavated in 1987/1988. The excavators deduced a discontinuity between the previous Roman settlement (until 300) and the early medieval colonization related to the burial ground that was in use between 530 and 700. Similar to the site of Petting, the resettlement was most likely dominated by *Iuvavum* (modern Salzburg, Austria) in the Southeast.
Although the 239 burials were subsequently analyzed and the artefacts went into a local exhibition after extensive conservation measures, the site is still not published in a comprehensive way. At least three burials were identified as multiple burials: the double burial 200/201 of an early mature male and infans II (WAG002; WAG003) were located on top of the burial of a potentially male infans II (WAG001). The double burial 37 contained remains of a potentially female infans II (WAG004) and a juvenile to early adult individual (WAG005). Burial 39 was identified as a double burial of a potentially female infans II (WAG006) and a late mature female (WAG007).
(31, 40)

Westheim (WES; Landkreis Weißenburg-Gunzenhausen, Germany):
The Merovingian cemetery of Westheim-Mehlbuck, excavated between 1979 and 1985, revealed remains of 255 individuals in 228 graves and dates to the 6th to mid-7th century. Five burials were clearly identified as double burials: 17a/b, 26a/b, 36a/b, 52a/b, 106a/b and 128a/b. Six additional burials were interpreted as presumably double burials of a child with a parent: 172, 190, 202, 206, 208 and 210. The triple burial I, uncovered during a previous excavation in the 1910s, held the remains of a late adult to early mature male (1-1, WES001), a late mature female (1-3, WES003) and of a third individual (1-2, WES002, min. adult, probably female). Burial 13a/b was reported by Lüdemann as a quadruple. However, according to Reiss, the burial contained only disturbed remains of two males (adult, mature).
(25, 41)

**Sources for mapping plague outbreaks between 541 and 650**

Clysma as possible starting point

    Tsiamis et al. (2009); Harper (2017) 215-218

541: Pelusium

    Prokopios *BP,* 2.22, 6
    See also: Stathakopoulos (2004) no. 102

541: Alexandria

    Prokopios, *BP*, 2.22, 6
    John of Ephesos, *Fragment E,* 77, 80
    Chron. Seert in *PO* 7, 185
    Michael the Syrian 2 (235-238)
    See also: Stathakopoulos (2004) no. 104

542: Jerusalem

    Cyril of Scythopolis, *Vita* of Kyriakos 10
    See also: Stathakopoulos (2004) no. 105

542: Antioch

    *Vita* of Symeon Stylites Iunior, 69
    See also: Stathakopoulos (2004) no. 107

542: Apamea

    Evagrios, *Hist. eccl.* 4.29
    See also: Stathakopoulos (2004) no. 108

542: Emesa

    Zacharias Rhetor, *Fragment ch. IX*
    Leontios of Neapolis, *Vita* of Symeon Salos, 151
    See also: Stathakopoulos (2004) no. 109

542: Myra

    *Vita* of Nicholas of Sion, 52
    See also: Stathakopoulos (2004) no. 110

542: Constantinople

    Prokopios, *BP,* 2.22, 2.23
    John of Ephesos, *Fragment E,* 74- 93
    John Malalas, 482
    Theophanes, *Chronographia,* AM 6034
    See also: Stathakopoulos (2004) no. 111

542: North Africa

    Victor of Tunnuna, *Chronica* ad a. 542 (201)
    See also: Stathakopoulos (2004) no. 114

542: Sicily

    *Byzantina Siciliae,* 133
    See also: Stathakopoulos (2004) no. 115

543: Sufetula

    Epigraphic material from Sufetula nos. 1-4 (277-280)

See also: Stathakopoulos (2004) no. 117

543: Italy, Illyricum

Marcellinus Comes ad a. 543 (107)

See also: Stathakopoulos (2004) no. 116

543: Gaul, Arles, Reims, Trier

Gregory of Tours, *Lib. hist.* 4.5, 6.15, 6.33

Gregory of Tours, *Lib. vitae patrum* 6.6, 17.4

Gregory of Tours, *Liber in gloria confessorum* 78

543: Spain

*Victoris Tunnunensis Chronicon*, *Consularia Caesaraugustana*, ad a. p.c. Basili II

See also: Kulikowski (2007) 150-151

543-544: Rome

*Inscriptiones Christianae Urbis Romae.* 1.1452, 2.4287, 7.17624, 2.5088, 2.4289, 8.20839, 2.5087, 2.5087, 2.5087.

See also: Stathakopoulos (2004) no. 118

544: Ireland, Britain

*Annals of Tigernach*, 137, 198

Adomnán, *Life of St Columba,* 348

See also: E. Phillimore (1888); A. Dooley (2007), 216; J. Maddicott (2007), 173-174

558: Constantinople

Agathias, *Hist.,* 5.10

John Malalas, *Chron.,* 18.127 (489)

Theophanes, *Chron.,* AM 6050

Agapios, *Kitab al-'Unwan*

See also: Stathakopoulos (2004) no. 134, Harper (2017) no. 1

561 – 62: Cilicia, Syria, Mesopotamia, Persia, Antioch

Theophanes, *Chron.,* AM 6053

Vita Symeon Stylites Iunior, 126- 29

Chron. ad a. 640

Chron. *Seert* in *PO* 7, 185- 86

Amr ibn Matta, 42 – 43

See also: Stathakopoulos (2004) no. 136, Harper (2017) no. 2

565: Liguria, Northern Italy

Paul the Deacon, *Hist. Langobardorum,* 2.4

See also: Stathakopoulos (2004) no. 139, Harper (2017) no. 3

571: Italy, Gaul, Bourges, Chalon-sur-Saône, Clermont, Dijon, Lyon

Marius of Avenches, a. 571

Gregory of Tours, *Lib. hist.* 4.31-32

See also: Stathakopoulos (2004) no. 144, Harper (2017) n. 4

573 – 74: Constantinople, Egypt, Syria and Antioch

John of Biclaro, a. 573

Agapios, *Kitab al-'Unwan*

John of Nikiu, 94.18
Chron. ad a. 846
Michael the Syrian, 10.8 (346)
See also: Stathakopoulos (2004) no. 145, Harper (2017) no. 5

576: Ireland
For secondary sources see:
Dooley (2007), 219
Woods (2003)

582- 84: Southwestern Gaul, Narbonne, Spain
Gregory of Tours, *Lib. hist. 6.*14, 6.33
See also: Harper (2017) no. 6

586: Constantinople
Agapios, *Kitab al-'Unwan*
See also: Harper (2017) no. 7

588: Gaul, Lyon, Marseille, Spain
Gregory of Tours, *Lib. hist.* 9.21-22
See also: Harper (2017) no. 8

590-91: Rome, Narni, Rhône Valley, Ravenna, Grado, Istria, Avignon, Viviers
Gregory of Tours, *Lib. hist.* 10.1, 10.23
Gregory the Great, *Dial.* 4.18, 4.26, 4.37; *Ep*. 2.2
Paul the Deacon, *Hist. Langobardorum,* 3.24, 4.4
*Liber pontificalis ,* 65
See also: Stathakopoulos (2004) nos. 151, 154; Harper (2017) nos. 9, 10

592: Syria, Palestine, Antioch
Evagrius, *Hist. eccl.* 4.29
I. Palaestina Tertia, nos. 68-70
Hassan ibn Thabit in Conrad (1981) 154
See also: Stathakopoulos (2004) no. 155, Harper (2017) no. 11

597: Thessalonica and countryside
*Mir. Demetr.,* 3 & 14
See also: Stathakopoulos (2004) no. 156, Harper (2017) no. 12

598: Thrace
Theophylact Simocatta, *Hist.* 7.15.2
See also: Stathakopoulos (2004) no. 159, Harper (2017) no. 13

599-600: Constantinople, Asia Minor, Syria, North Africa, Italy, Marseille
Michael the Syrian, 10.23 (387)
Chron. ad a. 1234
Gregory the Great, *Ep.* 9.232, 10.20
Paul the Deacon, *Hist. Langobardorum,* 4.14
Elias of Nisibis, a. 911
Fredegar, *Chron.* 4.18

See also: Stathakopoulos (2004) no. 160, Harper (2017) no. 14, Biraben and LeGoff (1975), 75

609: Cortijo de Chinales, Spain

CIL II 7.677

See also: Harper (2017) no. 15

McCormick (2016) 327

619: Constantinople, Alexandria

*Mirac. sanct. Artemii,* 34

See also: Stathakopoulos (2004) no. 173, Harper (2017) no. 17

626-28: Palestine, Mesopotamia

Michael the Syrian, 11.3 (409)

Arabic sources in Conrad (1981) 159-163

See also: Stathakopoulos (2004) nos. 177, 178; Harper (2017) no. 18

638-639: Palestine, Syria, Mesopotamia

Michael the Syrian, 11.8 (423)

Elias of Nisibis, (AH 18)

Chron. ad a. 1234,76 (AH 18)

Arabic sources in Conrad (1981) 167ff.

See also: Stathakopoulos (2004) no. 180, Harper (2017) no. 20

Primary sources:

Adomnan, *Life of St Columba = Adomnán's Life of St. Columba.* Ed. and. tr. A.O. Anderson and M. O. Anderson (Oxford, 1991)

*Annals of Tigernach = The Annals of Tigernach*. Ed. W. Stokes. *Revue Celtique* 16-17 (1895-96). Reprint, 2 vols. (Dyfed 1993).

Agapios = *Kitab al-'Unvan.* Ed. A. Vasiliev. In *PO* 5.4, 7.4; 8.3; 11.1 (1910, 1911, 1912, 1915).

Agathias = *Agathiae Myrinei Historiarum Libri Quinque.* Ed. R. Keydell. *CFHB* 2 (Berlin 1967).

Chron. ad a. 640 = *The Seventh Century in the West-Syrian Chronicles*. Ed. A. Palmer. *Translated texts for historians 15* (Liverpool 1993).

Chron. ad a. 846 = *Chronicon ad a. 846*. Ed. I. B. Chabot, E. W. Brooks. CSCO 4, Syr. 4 (Louvain 1907)

Chron. ad a. 1234 = *Anonymi auctoris Chronicon ad annum Christi 1234 pertinens.* Ed. J.-B. Chabot. CSCO 81, 82, 109; Syr. 36, 37, 56 (Paris 1916, 1920, 1937).

Chron. Seert = *Chronicle of Seert (Histoire Nestorienne).* Ed. A. Scher. in *PO* 4, 213-313; 5, 217-344; 7, 93-203.

*Victoris Tunnunensis Chronicon = Victoris Tunnunensis Chronicon cum reliquiis ex consularibus Caesaraugustanis et Iohannis Biclarensis Chronicon*. Ed. C. Cardelle de Hartmann. Corpus Christianorum, Series Latina 173A (Turnhout 2001).

Cyril of Scythopolis = *Kyrillos von Skytopolis.* Ed. E. Schwartz. TU 49.2 (Leipzig 1939).

Elias of Nisibis = *Opus Chronologicum.* Ed. E. W. Brooks. CSCO 63, Syr. 23 (Louvain 1964).

Evagrius *Hist. eccl.* = *Historia ecclesiastica.* Eds. J. Bidez, L. Parmentier. *The Ecclesiastical History of Evagrius* (London, 1898).

Fredegar, *Chron.* = Fredegar, *Chronica*. Ed. B. Krusch, SRM, 2.128.5-6 (Hannover 1888).

Gregory of Tours, *Lib. hist.* = *Libri historiarum X.* Eds. B. Krusch, W. Levison. MGH SRM, 1.1 (Hannover 1937-1951).

Gregory of Tours, *Liber in Gloria confessorum* = *Liber in Gloria confessorum*. B. Krusch, MGH, SRM, 1.2 (Hannover 1885).

Gregory of Tours, *Lib. vitae partum* = *Liber vitae partum.*.Ed. B. Krusch, MGH, SRM, 1.2 (Hannover 1885).

Gregory the Great, *Dial.* = *Dialogorum libri iv.* Ed. A: de Vogüé. SC 251, 260, 265 (Paris 1978-80).

Gregory the Great, *Ep.* = *Registrum epistularum.* Ed. D. Norberg. CC 140-40A (Turnhout 1982).

John of Biclaro = *Chronica.* Ed. T. Mommsen. MGH AA 9, 2, 213-214 (Berlin 1894).

John of Ephesos *Fragment E* = *Historiae Ecclesiasticae fragmenta.* Eds. W. J. van Douwen, J. P. N. Land. *Verhandelingen der koniglijke Akademie van Wetenschapen Af deeling Letterkunde* 18, 197-264 (Amsterdam 1899).

John Malalas = *Ioannis Malalae chronographia*. Ed. I. Thurn (Berlin 2000).

John of Nikiu = *The Chronicle of John, Bishop of Nikiu.* Ed. R. H. Charles (London 1916).

Leontios of Neapolis, *Vita of Symeon Salos* = *Das Leben des heiligen Narren Symeon*. Ed. L. Rydén. *Acta Universitatis Upsalensis. Studia Graeca Upsaliensia* 4, 151 (Stockholm, Göteborg, Upsala 1963).

*Liber pontificalis* = *Liber pontificalis.* Ed. T. Mommsen. MGH Gesta Pont. Rom. 1 (Hannover 1898).

Marius of Avenches = *Chronica.* Ed. T. Mommsen. MGH AA 9, 2, 225-240 (Berlin, 1894).

Marcellinus Comes = *Chronicon ad annum DXVIII.* Ed. T. Mommsen, MGH AA 11, 60-108 (Hannover 1894).

Michael the Syrian = *Chronique de Michel le Syrien, patriarche Jacobite d'Antioche (1166-1199).* Ed. J.-B. Chabot (Paris 1899-1924).

Mir. Demetr. = *Les plus anciens recueils de Miracles de Saint Démétrius et la pénétration des Slaves dans les Balkans.* P. Lemerle. 2 vols. (Paris 1979-1981).

Mirac. sanct. Artemii = *Miracula Sancti Artemii.* Ed. A. Papadopoulos-Kerameus. *Varia Graeca Sacra*. 34, 52 (Saint Petersburg 1909).

Paul the Deacon = *Historia Langobardorum*. Ed. G. Waitz. SS rer. Langobard. (Hannover 1878).

Prokopios *BP* = *Persian Wars*. Eds. J. Haury, G. Wirth, *Opera*. Vol. 1 (Leipzig 1962-1964).

Theophanes = *Theophanis chronographia.* Ed. C. de Boor (Leipzig 1883).

Theophylact Simocatta, *Hist.* = *Theophylacti Simocattae historiae.* Ed. C. de Boor (Leipzig 1997).

Victor of Tunnuna = *Chronica a. CCCCXLIV-DLXVII.* Ed. T. Mommsen, MGH AA 11, 184-206 (Berlin 1894).

*Vita* of Nicholas of Sion = *The Life of Saint Nicholas of Sion.* Eds. I. Ševčenko, N. P. Ševčenko (Brookline, 1984).

*Vita* of Symeon Stylites Iunior = *La vie ancienne de S. Syméon Stylite le jeune (521-592).* Ed. P. van den Ven (Brussels 1962).

Zacharias Rhetor = *Historia Ecclesiastica.* Ed. E. W. Brooks, CSCO Syr. 3.6 (Louvain, 1924).

Inscriptions:

Byzantina Siciliae = *Byzantina Siciliae*. Ed. G. Manganaro. *Minima Epigraphica et Papyrologica* 4, 133 (2001).

*Inscriptiones Christianae Urbis Romae = Inscriptiones Christianae Urbis Romae septimo seculo antiquiores, Nova Series* (= ICUR NS) Eds. I.B. de Rossi et al. 10 vols. (Rome 1922-92).

CIL = *Corpus Inscriptionum Latinarum* (Berlin 1863-).

Epigraphic material from Sufetula = Nouvelles recherches d'archéologie et d'épigraphie chrétienne à Sufetula (Byzacène). N. Duval. *Mélanges d'archéologie de l'École française de Rome* 68 (1956).

I. Palaestina Tertia = *Inscriptions from Palaestina Tertia*. Eds. Y. Meimaris, K. Kritikakou (Athens 2005).

Secondary sources:

L. I. Conrad: *The Plague in the Early Medieval Near East* (PhD, Princeton University 1981).

A. Dooley: The Plague and Its Consequences in Ireland. In: *Plague and the End of Antiquity: The Pandemic of 541-750.* Ed. L. K. Little (Cambridge 2007).

J.-N. Biraben, J. Le Goff: The Plague in the Early Middle Ages. In: *Biology of Man in History.* Ed. R. Forster and O. Ranum, trans. E. Forster and P. M. Ranum (Baltimore 1975).

K. Harper: *The Fate of Rome. Climate, Disease & the End of an Empire.* (Princeton, Oxford 2017)

M. Kulikowski: Plague in Spanish Late Antiquity. In: *Plague and the End of Antiquity: The Pandemic of 541-750.* Ed. L. K. Little (Cambridge 2007).

J. Maddicott: Plague in Seventh-Century England. In: *Plague and the End of Antiquity: The Pandemic of 541-750.* Ed. L. K. Little (Cambridge 2007).

M. McCormick: Tracking mass death during the fall of Rome's empire (II): a first inventory of mass graves. *Journal of Roman Archaeology* 29 (2016):1004-1046.

E. Phillimore: The Annales Cambriae and Old-Welsh Genealogies from Harlein MS 3859. *Y Cymmrodor* 9 (1888):141–83.

D. C. Stathakopoulos: *Famine and Pestilence in the Late Roman and Early Byzantine Empire.* (Burlington 2004).

C. Tsiamis et al.: The Red Sea and the Port of Clysma. A Possible Gate of Justinian's Plague. *Gesnerus* 66 (2009):209–217.
D. Woods: *Acorns, the Plague and the 'Iona Chronicle.' Peritia* 17–18 (2003–4): 495–502

### Supplementary Figures

**Fig. S1:** Heterozygosity plots for the six genomes considered in the phylogenetic trees. The y axis displays percentage of allele frequency, the reference calls (0 %) and alternative calls (100 %) were excluded for scaling.

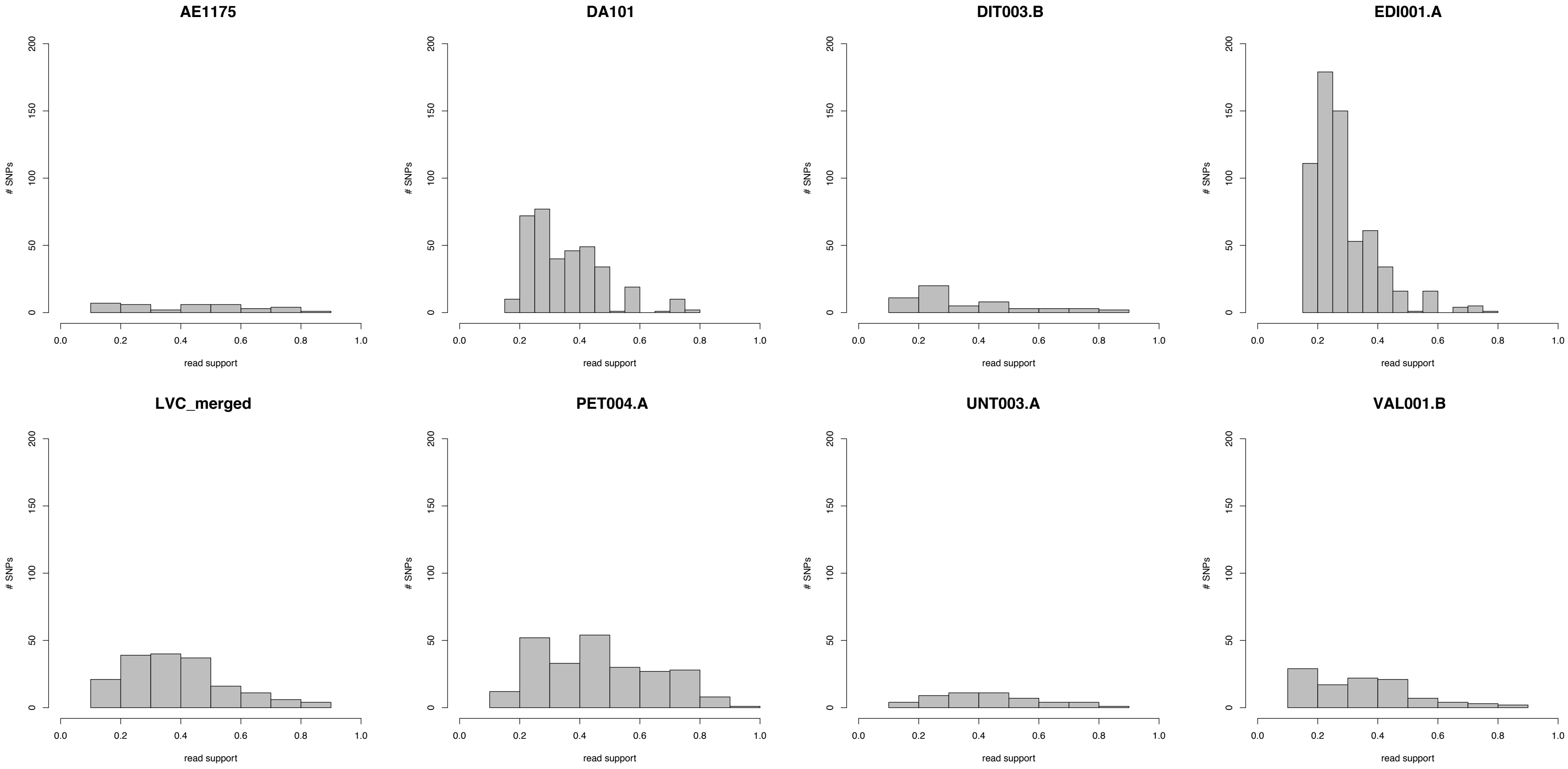

**Fig. S2:** Maximum Likelihood tree with full SNP alignment (6496 positions) of 233 modern *Y. pestis* and one *Y. pseudotuberculosis* genome, nine published (3rd century in orange; Altenerding in blue; Second Pandemic in red) and six newly reconstructed genomes (green). Numbers on node are showing bootstrap values (1000 iterations).

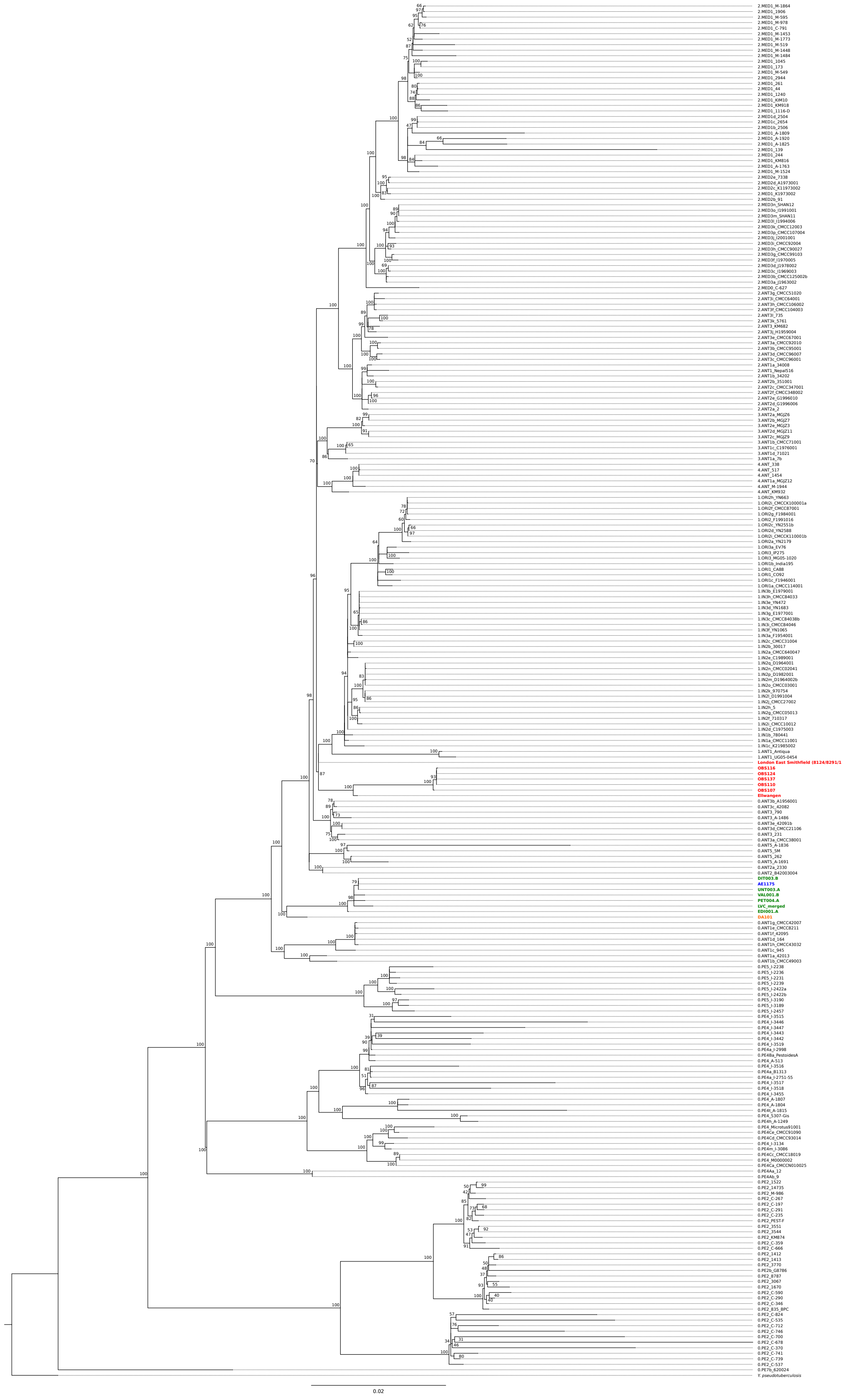

**Fig. S3:** Heatmap showing the percentage of coverage of plasmidal virulence factors in all analyzed First Pandemic genomes.

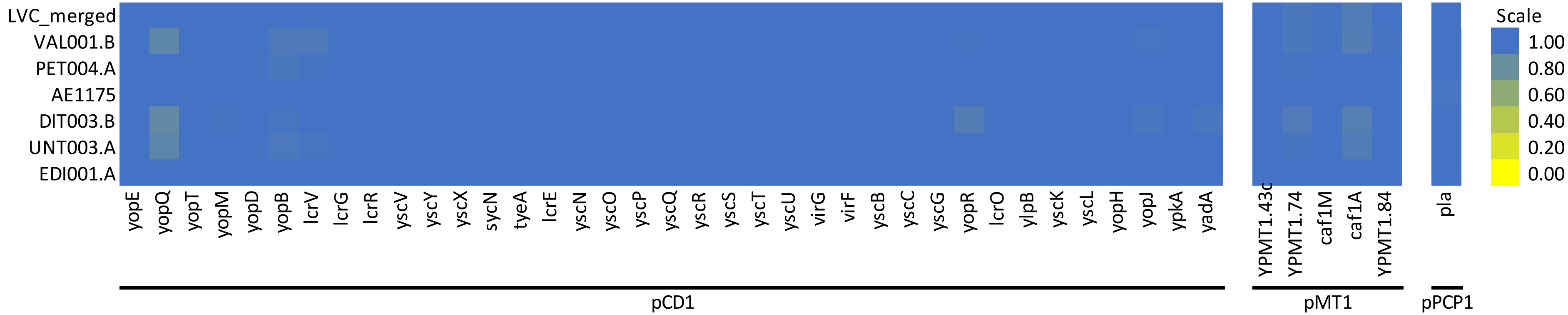

**Fig. S4:** Heatmap showing the percentage of coverage of the ca. 45 kb region of the chromosome deleted in LVC_merged and Marseille. First Pandemic genomes (blue and green) and Second Pandemic genomes (red) are shown in combination with selected strains of main clades of modern Y. pestis diversity on Branch 0 as well as the reference genomes of *Y. pseudotuberculosis* and *Y. pestis* (CO92).

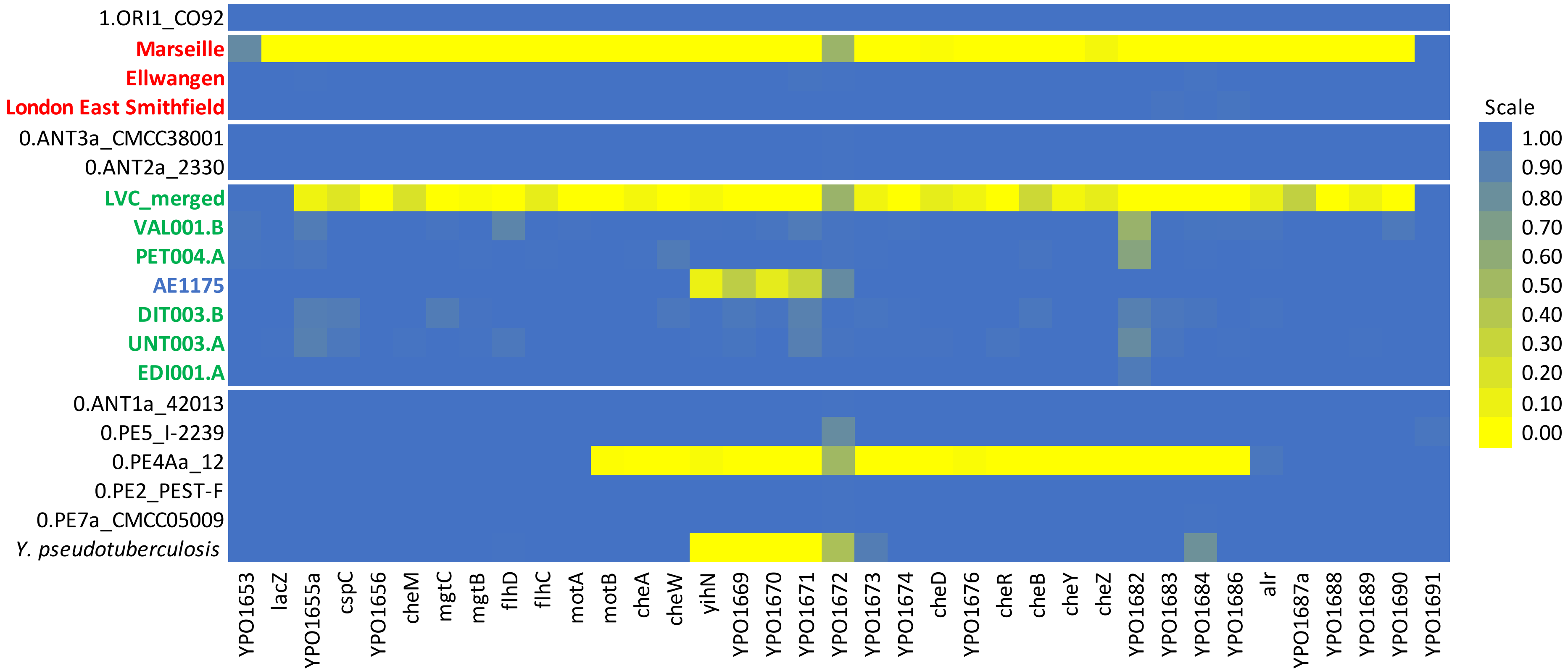

**Fig. S5:** Probability plots for all radiocarbon dates given in Table S9, cal AD 2σ.

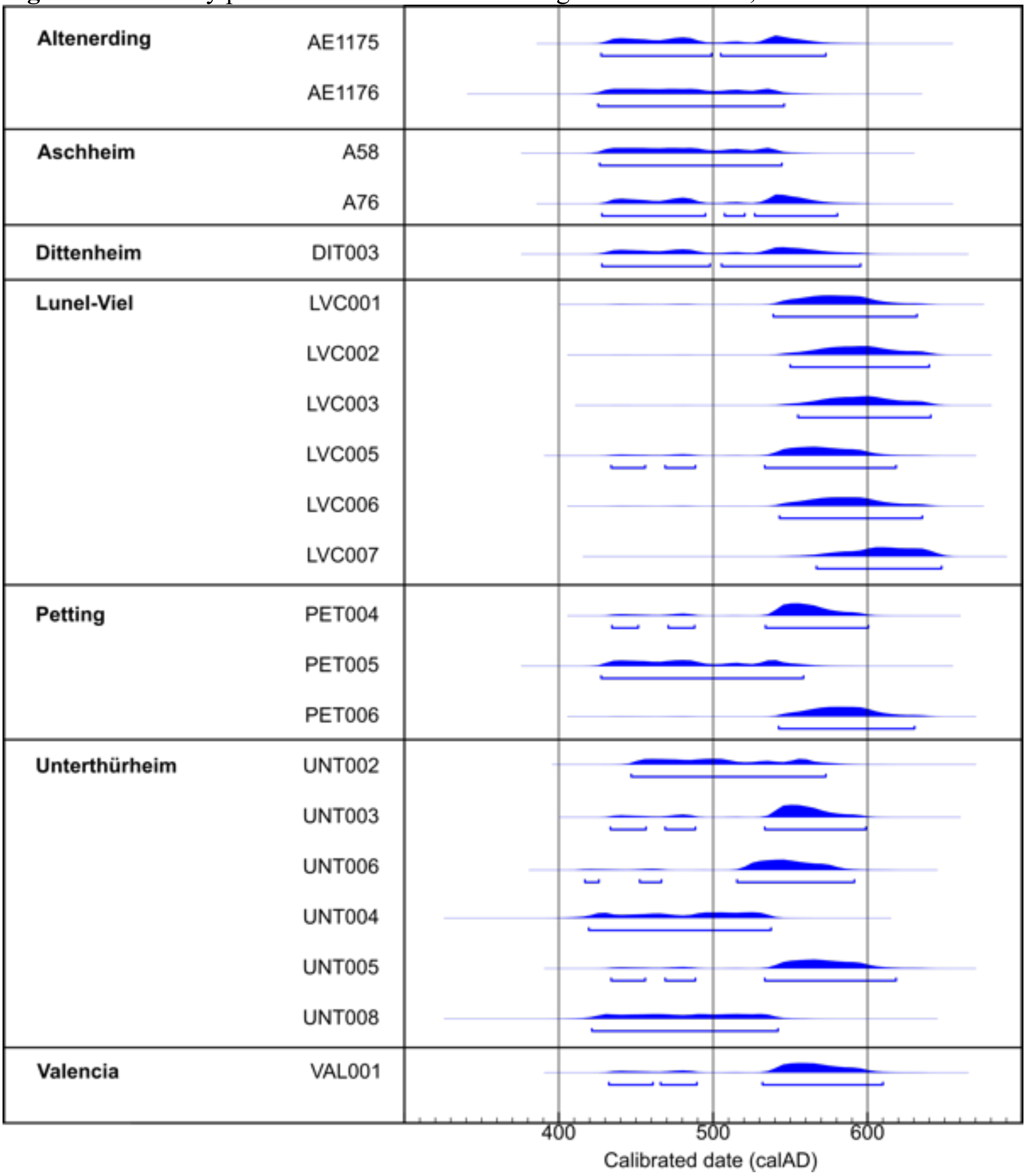

**Fig. S6:** Historical geography of the tested sites in Germany and Austria. A: Detailed map of all tested sites in Germany and Austria in relation to main rivers (blue), the Roman road network (orange) and the Roman provinces in their last state before the fall of the Western Roman Empire (shaded regions); all sources are given in the methods section of the main text. The sites of published genomes are depicted by pink squares, the sites for the genomes presented here are represented by yellow squares. Sites tested negative are depicted in black upward-pointing triangles (burials dating before 541), squares (dating around 541–544), downward-pointing triangles (dating after 544). B: Relation of the detailed map (rectangular space) to the sites in Britain (Edix Hill), France (Lunel-Viel) and Spain (Valencia) presented in this study.

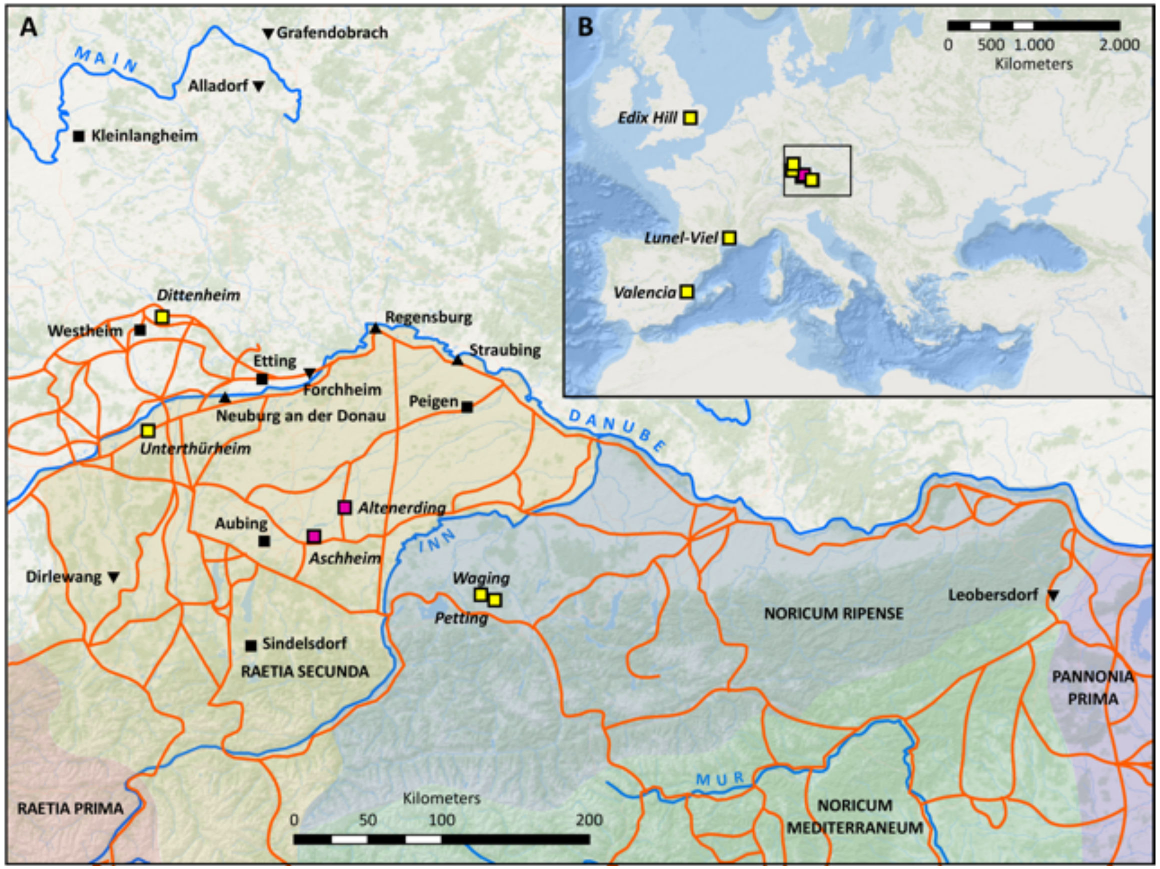

### Supplementary Tables

**Table S1:** Table with all tested individuals and samples. ‘Group’ gives the archaeological IDs of simultaneously buried individuals or individual from the same collective burial (Valencia: for details see site description). ‘Dating’ gives the range of the archaeological dating CE. ‘Total’ list the total number of individuals, further broken down to adults (>20 ya) and subadults (<20 ya). ‘Arch. ID’ and ‘Lab ID ind.’ are showing the individual ID with the morphological ‘sex’(f=female, m=male, ?=indeterminable) and ‘age’ (i=infans 1, ii=infans 2, j=juvenile, a=adult, m=mature, s=senile, (a)=at least adult, e=early, l=late). Teeth are noted in the FDI system or letters (i=Incisor, c=Canine, P=premolar, M=molar, lower case letters for deciduous teeth, indet=indeterminable). The qPCR screening results for *Y. pestis* are given in the ‘qPCR’ column, the ‘Seq’ column shows if samples were considered as positive after sequencing, ‘pos?’ refers here to a positive hit in the MALT screening (see Table S4).

| Site | Group | Dating | Total | Adult | Subadult | Arch. ID | Lab ID ind. | Sex | Age | Features | Teeth | Lab ID sample | qPCR | Seq |
|---|---|---|---|---|---|---|---|---|---|---|---|---|---|---|
| Alladorf | 179/180 | 700-900 | 2 | 1 | 1 | 179 | ALL001 | f | ea | | 47 | ALL001.A | neg | |
| | | | | | | 180 | ALL002 | ? | i | | 64 | ALL002.A | neg | |
| | 184/185 | 700-900 | 2 | 1 | 1 | 184 | ALL003 | ? | ii | | 85 | ALL003.A | neg | |
| | | | | | | 185 | ALL004 | m | la-em | | 28 | ALL004.A | neg | |
| | 203/204 | 700-900 | 2 | 1 | 1 | 203 | ALL005 | m | la | | 18 | ALL005.A | neg | |
| | | | | | | 204 | ALL006 | ? | i | May be a later burial | 74 | ALL006.A | neg | |
| Dirlewang | 33/34 | 650-700 | 2 | 1 | 1 | 33 | DIR001 | f | j-a | | 51 | DIR001.A | neg | |
| | 38/39 | 650-675 | 2 | 2 | 0 | 39 | DIR002 | f | a | | 17 | DIR002.A | pos | neg |
| Dittenheim | 188(A+B) | 550-700 | 2 | 0 | 2 | 188A | DIT001 | ? | i | | 37 | DIT001.A | neg | |
| | | | | | | 188B | DIT002 | ? | i | | 37 | DIT002.A | neg | |
| | 18(A+B) | 500-600 | 2 | 2 | 0 | 18A | DIT003 | m | m | | 18 | DIT003.A | neg | |
| | | | | | | | | | | | M | DIT003.B | pos | pos |
| | 18(A+B) | 500-600 | 2 | 2 | 0 | 18B | DIT004 | f | a | | 17 | DIT004.A | pos | pos |
| | | | | | | | | | | | M | DIT004.B | pos | pos |
| | 22(A+B) | 600-700 | 2 | 1 | 1 | 22A | DIT005 | ? | i | | 85 | DIT005.A | neg | |
| | | | | | | | DIT006 | f | a | | 38 | DIT006.A | neg | |
| | 8(A+B) | 500-700 | 2 | 2 | 0 | 8A | DIT007 | f | m | | 18 | DIT007.A | neg | |
| Edix Hill | Grave 76 | 500-550 | 1 | 0 | 1 | Sk405 | EDI001 | ? | j | | M | EDI001.A | - | pos |
| | Grave 69 | 500-550 | 1 | 1 | 0 | Sk359 | EDI002 | f | ea | | M | EDI002.A | - | pos? |
| | Grave 78 | 500-550 | 1 | 0 | 1 | Sk424 | EDI003 | ? | j | | M | EDI003.A | - | pos? |
| | Grave 96 | 500-550 | 2 | 1 | 1 | Sk547B | EDI004 | ? | ii | Simultaneous double burial | M | EDI004.A | - | pos? |
| | Grave 106 | 500-550 | 2 | 2 | 0 | Sk626A | EDI005 | f | ea | Simultaneous double burial | M | EDI005.A | - | pos? |
| | | | | | | Sk626B | EDI008 | m | ea | Simultaneous double burial | M | EDI008.A | - | pos? |
| | Grave 2 | 550-650 | 4 | 4 | 0 | Sk3C | EDI009 | m | ea | Sk3B and Sk3C simultaneous | M | EDI009.A | - | neg |
| | Grave 9 | 500-550 | 2 | 1 | 1 | Sk13A | EDI010 | m | m-s | Simultaneous double burial | M | EDI010.A | - | neg |
| | Grave 18 | 550-650 | 3 | 2 | 1 | Sk42B | EDI006 | f | ea | Nonsimultaneous double burial | M | EDI006.A | - | neg |
| | Grave 46 | 500-550 | 1 | 1 | 0 | Sk146 | EDI011 | m | em | | M | EDI011.A | - | neg |
| | Grave 60 | 550-650 | 1 | 1 | 0 | Sk183 | EDI012 | f | a | | M | EDI012.A | - | neg |
| | Grave 63 | 500-650 | 1 | 1 | 0 | Sk198 | EDI013 | m | ea | | M | EDI013.A | - | neg |
| | Grave 66 | 500-650 | 2 | 2 | 0 | Sk322A | EDI014 | m | (a) | Nonsimultaneous double burial | M | EDI014.A | - | neg |

| | | | | | | | | | | | | | | |
|---|---|---|---|---|---|---|---|---|---|---|---|---|---|---|
| | Grave 83 | 500-550 | 1 | 1 | 0 | Sk436 | EDI015 | f | ea | | M | EDI015.A | - | neg |
| | Grave 84 | 500-550 | 2 | 1 | 1 | Sk440A | EDI016 | f | a | Simultaneous double burial | M | EDI016.A | - | neg |
| | Grave 88 | 500-650 | 1 | 1 | 0 | Sk453A | EDI017 | m | ea | | M | EDI017.A | - | neg |
| | Grave 90 | 500-650 | 1 | 1 | 0 | Sk458 | EDI007 | f | s | | M | EDI007.A | - | neg |
| | Grave 95 | 550-550 | 1 | 1 | 0 | Sk530 | EDI018 | f | lm | | M | EDI018.A | - | neg |
| | Grave 97 | 550-650 | 1 | 1 | 0 | Sk551 | EDI019 | m | ea | | M | EDI019.A | - | neg |
| | Grave 99 | 500-650 | 1 | 1 | 0 | Sk576 | EDI020 | m | m | | M | EDI020.A | - | neg |
| | Grave 100 | 500-650 | 1 | 1 | 0 | Sk578 | EDI021 | m | ea | | M | EDI021.A | - | neg |
| | Grave 105 | 500-650 | 1 | 1 | 0 | Sk592 | EDI022 | f | lm | | M | EDI022.A | - | neg |
| Forchheim | 1-4 | 650-700 | 4 | 2 | 1 | 1 | FOR001 | m | j | | 27 | FOR001.A | neg | |
| | | | | | | 2 | FOR002 | f | m | Prone position | 27 | FOR002.A | neg | |
| | | | | | | 3 | FOR003 | m | m | | 47 | FOR003.A | neg | |
| Grafendobrach | 42/43/44 | 850-930 | 3 | 1 | 2 | 42 | GRA001 | m | es | | 46 | GRA001.A | neg | |
| | 83/84/85 | 850-93 | 3 | 2 | 1 | 83 | GRA002 | f? | em | May be a later burial | 16 | GRA002.A | neg | |
| | | | | | | 85 | GRA003 | f | ea | | 16 | GRA003.A | neg | |
| Kleinlangheim | 211/212 | 630-720 | 2 | min 1 | ? | 211 | KLH001 | m | m | Second burial: cremation | 47 | KLH001.A | neg | |
| | 218/219 | 560-630 | 2 | 2 | 0 | 218 | KLH002 | m | ma-m | | 17 | KLH002.A | neg | |
| | | | | | | 219 | KLH003 | f | la-em | | 23 | KLH003.A | neg | |
| | 272/273 | 670-720 | 2 | 2 | 0 | 273 | KLH004 | m | s | | 33 | KLH004.A | neg | |
| | 35/36 | 630-720 | 2 | 2 | 0 | 35 | KLH005 | m | a | | 37 | KLH005.A | neg | |
| Leobersdorf | 82(A-C) | 770-800 | 3 | 1 | 2 | 82A | LEO001 | m | j | | 36 | LEO001.A | neg | |
| | | | | | | 82B | LEO002 | f | a | | 27 | LEO002.A | neg | |
| | | | | | | 82C | LEO003 | ? | i | | 55 | LEO003.A | neg | |
| Lunel-Viel (Les Horts) | 38(A-C) | 475-550 | 3 | 3 | 0 | 38C | LVH001 | ? | (a) | Double burial with remains of a third individual | M | LVH001.A | neg | |
| | | | | | | | | | | | M | LVH001.B | neg | |
| | | | | | | | | | | | M | LVH001.C | neg | |
| | | | | | | | | | | | I | LVH001.D | neg | |
| | | | | | | | | | | | I | LVH001.E | neg | |
| Lunel-Viel (Quartier centrale) | 1-5 | 400-600 | 6 | 6 | 0 | 1 | LVC001 | f | ea | | M | LVC001.A | neg | |
| | | | | | | | | | | | M | LVC001.B | pos | pos |
| | | | | | | | | | | | 26 | LVC001.C | pos | pos |
| | | | | | | 2 | LVC002 | ? | la-em | | M | LVC002.A | neg | |
| | | | | | | | | | | | M | LVC002.B | neg | |
| | | | | | | 3A | LVC003 | f | a | | M | LVC003.A | neg | |
| | | | | | | | | | | | M | LVC003.B | neg | |
| | | | | | | 3B | LVC004 | f | a | | M | LVC004.A | neg | |
| | | | | | | | | | | | M | LVC004.B | neg | |
| | | | | | | 4 | LVC005 | m | m | | M | LVC005.A | pos | pos |
| | | | | | | | | | | | M | LVC005.B | pos | pos |
| | | | | | | | | | | | 43 | LVC005.C | pos | pos |
| | | | | | | 5 | LVC006 | m | ea | | M | LVC006.A | pos | pos |
| | | | | | | | | | | | M | LVC006.B | neg | |
| | 6/7 | 400-600 | 2 | 0 | 2 | 6 | LVC007 | f | j-ea | | M | LVC007.A | neg | |
| | | | | | | | | | | | M | LVC007.B | neg | |
| München-Aubing | 675/676/677 | 650-700 | 3 | 2 | 1 | 676 | AUB002 | f | m | | 13 | AUB002.A | neg | |
| | | | | | | 677 | AUB003 | m | ? | | 16 | AUB003.A | neg | |

| | | | | | | | | | | | | | | |
|---|---|---|---|---|---|---|---|---|---|---|---|---|---|---|
| | 724/725 | 400-700 | 2 | 2 | 0 | 724 | AUB004 | m | a | | 46 | AUB004.A | neg | |
| | | | | | | 725 | AUB005 | m | m | | 31 | AUB005.A | neg | |
| | 809/810 | 400-700 | 2 | 2 | 0 | 809 | AUB006 | m | a | | 16 | AUB006.A | neg | |
| | | | | | | 810 | AUB007 | m | a | | 36 | AUB007.A | neg | |
| | 854/855 | 650-700 | 2 | 2 | 0 | 854 | AUB008 | m | a | | 47 | AUB008.A | neg | |
| | 675/676/677 | 650-700 | 3 | 2 | 1 | 675 | AUB009 | ? | i | | 85 | AUB009.A | neg | |
| Neuburg an der Donau | 34(N/M/S) | 330-360 | 3 | 3 | 0 | 34M | NEU001 | m | m | | 13 | NEU001.A | neg | |
| | | | | | | 34S | NEU002 | m | a | | 37 | NEU002.A | neg | |
| Peigen | 18(1-2) | 600-700 | 2 | 2 | 0 | 18-1 | PEI001 | f | a | Opposite orientation | 37 | PEI001.A | pos | neg |
| | | | | | | 18-2 | PEI002 | f | ea | Opposite orientation | 47 | PEI002.A | neg | |
| | 62(1-2) | 600-650 | 2 | 1 | 1 | 62-2 | PEI003 | f | lm | | 38 | PEI003.A | neg | |
| | 63(1-2) | 580-620 | 2 | 2 | 0 | 63-1 | PEI004 | f | a | | 47 | PEI004.A | neg | |
| | | | | | | 63-2 | PEI005 | m | s | | 45 | PEI005.A | neg | |
| Petting | 342/343/344 | 530-730 | 3 | 2 | 1 | 342 | PET001 | m | ea | | 37 | PET001.A | neg | |
| | | | | | | 343 | PET002 | ? | ii | | 74 | PET002.A | neg | |
| | 377/378 | 570-600 | 2 | 2 | 0 | 377 | PET003 | m | la-m | | 26 | PET003.A | neg | |
| | | | | | | 378 | PET004 | m | la-em | | 36 | PET004.A | pos | pos |
| | 628 | 530-730 | 1 | 0 | 1 | 628 | PET005 | ? | ii | Only a few remains, directly above 630/631 | 37 | PET005.A | pos | pos |
| | 630/631 | 570-600 | 2 | 2 | 0 | 630 | PET006 | f | ea | | 46 | PET006.A | pos | pos |
| | | | | | | 631 | PET007 | f | em | | 46 | PET007.A | neg | |
| Regensburg | 30(1-2) | 350-450 | 2 | 2 | 0 | 30-1 | RFF001 | m | la-em | Prone position | 36 | RFF001.A | neg | |
| | 53(2/3) | 350-450 | 2 | 1 | 1 | 53-2 | RFF002 | f | j | Crouched inhumation | 26 | RFF002.A | neg | |
| | | | | | | 53-3 | RFF003 | m | em | Crouched inhumation | 46 | RFF003.A | neg | |
| Sindelsdorf | 162 | 630-720 | 1 | 1 | 0 | 162 | SIN001 | m | s | 162 directly above 163-165 | 48 | SIN001.A | neg | |
| | 163-165 | 630-720 | 3 | 2 | 1 | 164 | SIN002 | m | la-em | | 17 | SIN002.A | neg | |
| | | | | | | 165 | SIN003 | f? | m | | 47 | SIN003.A | neg | |
| | 51/52 | 630-670 | 2 | 2 | 0 | 52-1 | SIN006 | m | s | | 37 | SIN006.A | neg | |
| | | | | | | 52-2 | SIN007 | m | m | | 33 | SIN007.A | neg | |
| Straubing | 16/17 | 300-400 | 2 | 0 | 2 | 16 | SAZ001 | ? | i | | 85 | SAZ001.A | neg | |
| | 54a-c | 300-400 | 3 | 1 | 2 | 54a | SAZ002 | m? | j | | 26 | SAZ002.A | neg | |
| | | | | | | 54b | SAZ003 | m | j | | 27 | SAZ003.A | neg | |
| Unterthürheim | 116(1-2) | 525-680 | 2 | 1 | 1 | 116(-2) | UNT001 | ? | ii | | 16 | UNT001.A | neg | |
| | 129 | 550-600 | 1 | 0 | 1 | 129 | UNT002 | ? | i | Close to 131-134 | 54 | UNT002.A | pos | pos |
| | 130 | 550-600 | 1 | 1 | 0 | 130 | UNT003 | m | m | Close to 131-134 | 37 | UNT003.A | pos | pos |
| | 131/132/133/134 | 550-600 | 4 | 2 | 2 | 131 | UNT004 | ? | i | | 55 | UNT004.A | pos | pos |
| | | | | | | 132 | UNT005 | m | a | | 48 | UNT005.A | pos | pos |
| | 63/64 | 550-600 | 2 | 1 | 1 | 63 | UNT006 | f | a | | 36 | UNT006.A | pos | pos |
| | 63/64 | 550-600 | 2 | 1 | 1 | 64 | UNT007 | ? | i | | 84 | UNT007.A | neg | |
| Valencia | 41 | 500-700 | 15 | ? | min 4 | 1657 | VAL001 | ? | ? | Collective burial | M | VAL001.A | neg | |
| | | | | | | | | | | | M | VAL001.B | pos | pos |
| | | | | | | | | | | | P | VAL001.C | neg | |
| | | | | | | | | | | | m | VAL001.D | neg | |
| | | | | | | | | | | | P | VAL001.E | neg | |
| | | | | | | | | | | | m | VAL001.F | neg | |
| | 40 | 500-700 | 4 | ? | ? | 1728 | VAL002 | ? | ? | Burial on apse entrance | I | VAL002.A | neg | |

| | | | | | | | | | | | | | | |
|---|---|---|---|---|---|---|---|---|---|---|---|---|---|---|
| | | | | | | | | | | | M | VAL002.B | neg | |
| | | | | | | | | | | | C | VAL002.C | neg | |
| | | | | | | | | | | | indet. | VAL002.D | neg | |
| | | | | | | | | | | | indet. | VAL002.E | neg | |
| | 50 | 500-700 | 7 | min 2 | min 1 | 10039 | VAL003 | ? | ? | Collective burial next to apse | P | VAL003.A | neg | |
| | | | | | | | | | | | M | VAL003.B | neg | |
| | | | | | | | | | | | M | VAL003.C | neg | |
| | | | | | | | | | | | I | VAL003.D | neg | |
| | | | | | | | | | | | I | VAL003.E | neg | |
| | | | | | | | | | | | I | VAL003.F | neg | |
| | | | | | | 10346 | VAL004 | ? | ? | Collective burial next to apse | C | VAL004.A | neg | |
| | 28 | 500-700 | min 21 | min 13 | min 3 | 10307 | VAL005 | ? | ? | Collective burial next to apse | M | VAL005.A | neg | |
| | | | | | | | | | | | M | VAL005.B | neg | |
| | | | | | | | | | | | M | VAL005.C | neg | |
| | | | | | | | | | | | M | VAL005.D | neg | |
| | | | | | | | | | | | M | VAL005.E | neg | |
| | | | | | | | | | | | M | VAL005.F | neg | |
| | | | | | | | | | | | M | VAL005.G | neg | |
| | | | | | | | | | | | M | VAL005.H | neg | |
| | | | | | | | | | | | Indet. | VAL005.I | neg | |
| | | | | | | | | | | | indet. | VAL005.J | neg | |
| | | | | | | | | | | | M | VAL005.K | neg | |
| | | | | | | | | | | | indet. | VAL005.L | neg | |
| | | | | | | 10313 | VAL006 | ? | ? | Collective burial next to apse | M | VAL006.A | neg | |
| | | | | | | | | | | | m | VAL006.B | neg | |
| | | | | | | 10322 | VAL007 | ? | ? | Collective burial next to apse | M | VAL007.A | neg | |
| | 4 | 500-700 | 4 | 3 | 1 | 10066 | VAL008 | ? | ? | *Tegula* burial | M | VAL008.A | neg | |
| | | | | | | | | | | | 27 | VAL008.B | neg | |
| | 28 | 500-700 | min 21 | min 13 | min 3 | 10339 | VAL009 | ? | ? | Collective burial next to apse | 27 | VAL009.A | neg | |
| Waging | 199 | 600-700 | 1 | 1 | 0 | 199 | WAG001 | m | ii | Superposition (under 200/201) | 55 | WAG001.A | pos | pos |
| | | | | | | | | | | | M | WAG001.B | neg | |
| | 200/201 | 600-700 | 2 | 1 | 1 | 200 | WAG002 | m | em | | 36 | WAG002.A | neg | |
| | | | | | | | | | | | M | WAG002.B | neg | |
| | | | | | | 201 | WAG003 | ? | ii | | 85 | WAG003.A | neg | |
| | | | | | | | | | | | M | WAG003.B | neg | |
| | 37(1-2) | 600-700 | 2 | 1 | 1 | 37-1 | WAG004 | f? | ii | Superposition over 39, disturbed | 46 | WAG004.A | neg | |
| | | | | | | 37-2 | WAG005 | ? | j-ea | | 62 | WAG005.A | neg | |
| | 39(N/S) | 600-700 | 2 | 1 | 1 | 39N | WAG006 | f | ii | | 75 | WAG006.A | neg | |
| | | | | | | | | | | | m | WAG006.B | neg | |
| | | | | | | 39S | WAG007 | f | lm | | 43 | WAG007.A | neg | |
| | | | | | | | | | | | P | WAG007.B | neg | |
| Westheim | 1 | 500-650 | 3 | 2 | 1 | 1(-1?) | WES001 | m | la-em | | 27 | WES001.A | neg | |
| | | | | | | 1(-2?) | WES002 | f? | (a) | | 18 | WES002.A | neg | |
| | | | | | | 1(-3?) | WES003 | f | lm | | 26 | WES003.A | neg | |

**Table S2:** Mapping results of all UDG treated and captured samples for the chromosome and all three plasmids, giving the number of unique reads after clipping, merging, quality filtering and duplicate removal, and the mean coverage. In the last three columns, the ratio of mapping reads to the respective plasmid/chromosome is calculated for each sample.

| Sample | Chrom. reads | Chrom. coverage | pCD1 reads | pCD1 coverage | pMT1 reads | pMT1 coverage | pPCP1 reads | pPCP1 coverage | Ratio reads pCD1/chrom. | Ratio reads pMT1/chrom. | Ratio reads pPCP1/chrom. | Mahalanobis distance |
|---|---|---|---|---|---|---|---|---|---|---|---|---|
| DIR002.A | 8470 | 0.08 | 10 | 0.01 | 9 | 0.01 | 3 | 0.01 | 0.00118 | 0.00106 | 0.00035 | 11.50 |
| DIT003.B | 877150 | 9.41 | 31613 | 23.44 | 26800 | 14.12 | 7777 | 49.80 | 0.03604 | 0.03055 | 0.00887 | 0.83 |
| DIT004.A | 192557 | 1.65 | 7638 | 4.40 | 6383 | 2.74 | 2043 | 9.06 | 0.03967 | 0.03315 | 0.01061 | 0.98 |
| DIT004.B | 21824 | 0.22 | 849 | 0.58 | 935 | 0.46 | 220 | 1.12 | 0.03890 | 0.04284 | 0.01008 | 2.10 |
| LVC001.B | 40317 | 0.41 | 1731 | 1.19 | 1731 | 0.86 | 1176 | 6.33 | 0.04293 | 0.04293 | 0.02917 | 5.37 |
| LVC001.C | 182392 | 1.78 | 8385 | 5.54 | 6156 | 2.96 | 5694 | 28.43 | 0.04597 | 0.03375 | 0.03122 | 6.27 |
| LVC005.A | 162315 | 1.81 | 7300 | 5.56 | 5327 | 2.94 | 2022 | 12.14 | 0.04497 | 0.03282 | 0.01246 | 2.24 |
| LVC005.B | 3628 | 0.04 | 114 | 0.08 | 132 | 0.07 | 63 | 0.30 | 0.03142 | 0.03638 | 0.01736 | 1.36 |
| LVC005.C | 276945 | 2.68 | 8392 | 5.43 | 9594 | 4.55 | 6033 | 31.11 | 0.03030 | 0.03464 | 0.02178 | 3.27 |
| LVC006.A | 41524 | 0.43 | 1178 | 0.81 | 1673 | 0.85 | 277 | 1.44 | 0.02837 | 0.04029 | 0.00667 | 2.73 |
| PEI001.A | 1133 | 0.01 | 6 | 0.00 | 36 | 0.03 | 0 | 0.00 | 0.00530 | 0.03177 | 0.00000 | 9.36 |
| PET004.A | 515024 | 5.62 | 19971 | 14.86 | 17721 | 9.54 | 4442 | 26.78 | 0.03878 | 0.03441 | 0.00862 | 1.29 |
| PET005.A | 55916 | 0.49 | 2213 | 1.31 | 2395 | 1.04 | 589 | 2.62 | 0.03958 | 0.04283 | 0.01053 | 2.01 |
| PET006.A | 19144 | 0.20 | 645 | 0.46 | 746 | 0.38 | 209 | 1.09 | 0.03369 | 0.03897 | 0.01092 | 0.94 |
| UNT002.A | 14842 | 0.14 | 648 | 0.43 | 546 | 0.28 | 311 | 1.52 | 0.04366 | 0.03679 | 0.02095 | 1.23 |
| UNT003.A | 753340 | 7.57 | 21453 | 14.59 | 18224 | 9.13 | 5836 | 32.65 | 0.02848 | 0.02419 | 0.00775 | 0.87 |
| UNT004.A | 533323 | 5.19 | 20168 | 13.53 | 13211 | 6.45 | 4671 | 24.91 | 0.03782 | 0.02477 | 0.00876 | 2.72 |
| UNT005.A | 13669 | 0.15 | 578 | 0.40 | 448 | 0.24 | 171 | 0.90 | 0.04229 | 0.03277 | 0.01251 | 1.31 |
| UNT006.A | 7322 | 0.07 | 177 | 0.12 | 159 | 0.08 | 80 | 0.42 | 0.02417 | 0.02172 | 0.01093 | 1.34 |
| VAL001.B | 996679 | 9.62 | 27992 | 18.30 | 30071 | 14.29 | 5520 | 29.05 | 0.02809 | 0.03017 | 0.00554 | 0.77 |
| WAG001.A | 22904 | 0.23 | 733 | 0.50 | 539 | 0.26 | 351 | 1.93 | 0.03200 | 0.02353 | 0.01532 | 1.50 |

**Table S3:** Sequencing results for all verified *Y. pestis*-positive samples. The data of the Altenerding sample AE1175 was reprocessed without additional capture or sequencing. Data was merged in case multiple samples of the same individual were positive (DIT004, LVC001, LVC005) or when samples of multiple individuals of the same burial context were merged (LVC). *Without capture.

| Sample | Reads prior mapping after adapter clipping and merging | Unique reads mapping to *Y. pestis* reference | Endogenous DNA (%) | Cluster Factor | Mean Coverage | Coverage 1X (%) | Median frag. Length |
|---|---|---|---|---|---|---|---|
| AE1175 | 67364973 | 1164051 | 3.56 | 2.06 | 17.17 | 93.62 | 64 |
| DIT003.B | 12699975 | 877150 | 22.88 | 3.31 | 9.41 | 91.41 | 48 |
| DIT004.A | 65225088 | 192557 | 1.02 | 3.44 | 1.65 | 69.02 | 38 |
| DIT004.B | 4361688 | 21824 | 1.80 | 3.59 | 0.22 | 17.41 | 45 |
| DIT004 merged | 70153762 | 215192 | 1.06 | 3.46 | 1.88 | 72.67 | 39 |
| EDI001.A* | 544200122 | 914881 | 0.20 | 1.22 | 9.10 | 94.58 | 44 |
| LVC001.B | 3593715 | 40317 | 3.65 | 3.25 | 0.41 | 28.23 | 45 |
| LVC001.C | 16970946 | 182392 | 5.76 | 3.29 | 1.78 | 72.48 | 43 |
| LVC001 merged | 20564661 | 220514 | 3.56 | 3.32 | 2.17 | 76.42 | 43 |
| LVC005.A | 3299048 | 162315 | 14.12 | 2.87 | 1.81 | 70.13 | 50 |
| LVC005.B | 3634255 | 3628 | 0.27 | 2.75 | 0.04 | 3.09 | 44 |
| LVC005.C | 20373965 | 276945 | 7.44 | 3.61 | 2.68 | 80.19 | 43 |
| LVC005 merged | 27307268 | 430854 | 5.40 | 3.43 | 4.42 | 86.67 | 45 |
| LVC006.A | 4480909 | 41524 | 3.29 | 3.55 | 0.43 | 29.66 | 46 |
| LVC merged | 52352838 | 658940 | 4.50 | 3.57 | 6.71 | 89.89 | 45 |
| VAL001.B | 65109736 | 996679 | 4.28 | 2.80 | 9.62 | 91.35 | 43 |
| PET004.A | 52013500 | 515024 | 2.69 | 2.72 | 5.62 | 92.30 | 48 |
| PET005.A | 89496419 | 55916 | 0.34 | 5.51 | 0.49 | 34.62 | 39 |
| PET006.A | 48802311 | 19144 | 0.12 | 3.02 | 0.20 | 16.10 | 44 |
| UNT002.A | 88974245 | 14842 | 0.20 | 12.15 | 0.14 | 10.97 | 41 |
| UNT003.A | 37303508 | 753340 | 6.11 | 3.03 | 7.57 | 92.65 | 44 |
| UNT004.A | 26118776 | 533323 | 7.07 | 3.46 | 5.19 | 91.22 | 42 |
| UNT005.A | 3613234 | 13669 | 0.49 | 1.29 | 0.15 | 13.24 | 45 |
| UNT006.A | 91966889 | 7322 | 0.08 | 9.64 | 0.07 | 4.98 | 40 |
| WAG001.A | 74283932 | 22904 | 0.13 | 4.31 | 0.23 | 13.89 | 43 |

**Table S4:** Number of reads of EDI samples assigned to the *Y. pseudotuberculosis* complex/*Y. pestis* node in MALT with 85 % identity.

| Samples | Total # of reads | *Y. pseudotuberculosis* complex node | | *Y. pestis* node |
|---|---|---|---|---|
| | | assigned | summed | summed |
| EDI001.A | 14183537 | 26478 | 36623 | 9886 |
| EDI002.A | 10714470 | 44 | 58 | 14 |
| EDI003.A | 41364951 | 3485 | 4738 | 1204 |
| EDI004.A | 5792548 | 743 | 991 | 230 |
| EDI005.A | 9029341 | 21 | 33 | 4 |
| EDI006.A | 7838268 | 6 | 10 | 1 |
| EDI007.A | 44427907 | 21 | 27 | 1 |
| EDI008.A | 4544207 | 79 | 110 | 29 |
| EDI009.A | 9918621 | 12 | 25 | 2 |
| EDI010.A | 4767075 | 0 | 0 | 0 |
| EDI011.A | 8639291 | 4 | 7 | 0 |
| EDI012.A | 38492923 | 145 | 259 | 29 |
| EDI013.A | 17140889 | 7 | 8 | 1 |
| EDI014.A | 8189593 | 0 | 2 | 0 |
| EDI015.A | 10412796 | 5 | 10 | 1 |
| EDI016.A | 27869518 | 44 | 69 | 6 |
| EDI017.A | 9565231 | 10 | 16 | 0 |
| EDI018.A | 2040312 | 3 | 4 | 1 |
| EDI019.A | 10904548 | 2 | 6 | 2 |
| EDI020.A | 29162638 | 29 | 45 | 2 |
| EDI021.A | 10637817 | 4 | 9 | 1 |
| EDI022.A | 9585693 | 23 | 31 | 4 |

**Table S5:** Table with all non-shared SNPs that were called in the investigated First Pandemic genomes, sorted by genome (second cell in header). 'Position' refers to the position in the reference genome CO92, followed by the respective SNP call and the reference call. The classification as false or true positive is based on the preceding parameters regarding a 50 bp window surrounding the SNP: Only SNPs with no uncovered position, no heterozygous SNP and an equal mean coverage with high and low stringent mapping (ratio of 1.00) are classified as true positive SNPs. The 'comment' column with additional classification is followed by the respective calls in the other First Pandemic genomes (0=not covered, N=heterozygous, lower case letters=less than 3 reads coverage). The last column is giving the corresponding genomes, if the SNP appears as homoplastic in the SNP table.

| **Position** | **AE1175 call** | Reference call | Uncovered positions in 50 bp | Heterozygous SNPs in 50 bp | Mean coverage ratio LS/HS | Classification | Comment | DIT003.B | EDI001.A | LVC_merged | PET004.A | UNT003.A | UNT004.A | VAL001.B | WAG001.A | Homoplasy |
|---|---|---|---|---|---|---|---|---|---|---|---|---|---|---|---|---|
| 20769 | G | A | 0 | 1 | 1.00 | false positive | | 0 | a | 0 | A | 0 | a | 0 | 0 | 2.MED2e_7338 |
| 420208 | T | G | 20 | 0 | 1.00 | false positive | Feldman et al. 2016: Called as SNP but not called as unique due to homoplasy | 0 | 0 | 0 | 0 | 0 | 0 | 0 | 0 | 0.PE2_6904 |
| 549767 | C | T | 0 | 0 | 1.45 | false positive | Feldman et al. 2016 | C | T | c | c | C | C | c | 0 | |
| 2348782 | T | C | 5 | 0 | 1.00 | false positive | | 0 | t | 0 | 0 | t | 0 | t | 0 | |
| 2828136 | G | A | 0 | 3 | 1.39 | false positive | | N | N | 0 | 0 | a | a | g | 0 | |
| 2980096 | T | G | 0 | 1 | 1.38 | false positive | | t | g | g | 0 | 0 | 0 | g | 0 | 2.ANT2b_351001 |
| 2980097 | G | A | 0 | 1 | 1.37 | false positive | | g | a | a | 0 | 0 | 0 | a | 0 | 2.ANT2b_351001 |
| 3179828 | A | C | 0 | 0 | 1.00 | true positive | shared AE, DIT, UNT | A | C | C | C | A | A | C | 0 | |
| 3223354 | T | G | 0 | 1 | 1.13 | false positive | | 0 | g | 0 | 0 | 0 | g | 0 | 0 | 1.IN2i_CMCC10012 |
| 3223359 | A | C | 0 | 1 | 1.13 | false positive | | 0 | N | 0 | a | 0 | c | 0 | 0 | 1.IN2i_CMCC10012, 1.IN1a_CMCC11001 |
| 3225856 | A | C | 23 | 0 | 1.24 | false positive | | a | 0 | 0 | 0 | c | a | c | 0 | |
| 3588894 | C | A | 0 | 1 | 1.17 | false positive | | a | c | a | 0 | 0 | 0 | a | 0 | |
| 3750736 | A | G | 0 | 0 | 0.99 | true positive | Feldman et al. 2016, shared AE, DIT, LVC, UNT, VAL | a | G | a | A | A | A | A | a | |
| 3755861 | T | C | 0 | 0 | 1.00 | true positive | Feldman et al. 2016, potentially shared ancestral | 0 | T | t | t | t | t | 0 | 0 | |
| 4412624 | G | A | 0 | 0 | 1.00 | false positive? | Feldman et al. 2016, false positive in UNT003.A | 0 | A | 0 | G | G | g | g | 0 | |

| Position | **DIT003.B call** | Reference call | Uncovered positions in 50 bp | Heterozygous SNPs in 50 bp | Mean coverage ratio LS/HS | Classification | Comment | AE1175 | EDI001.A | LVC_merged | PET004.A | UNT003.A | UNT004.A | VAL001.B | WAG001.A | Homoplasy |
|---|---|---|---|---|---|---|---|---|---|---|---|---|---|---|---|---|
| 335336 | C | T | 0 | 1 | 1.00 | false positive | | N | N | 0 | 0 | N | N | c | 0 | 0.PE4Cd_CMCC93014, 2.ANT1_Nepal516, 0.PE4_Microtus91001, 0.PE5_6213, 0.PE4_6216, Y. pseudotuberculosis |
| 549767 | C | T | 0 | 0 | 1.17 | false positive | Feldman et al. 2016 | C | T | c | c | C | C | c | 0 | |
| 944177 | G | C | 6 | 0 | 1.00 | false positive | | N | C | c | c | N | N | N | 0 | |
| 944178 | A | G | 6 | 0 | 1.00 | false positive | | N | G | g | g | N | g | N | 0 | |
| 3179828 | A | C | 0 | 0 | 1.00 | true positive | shared AE, DIT, UNT | A | C | C | C | A | A | C | 0 | |
| 3225949 | G | A | 21 | 0 | 1.12 | false positive | | g | 0 | a | 0 | 0 | 0 | g | 0 | |
| 3890928 | G | C | 4 | 0 | 1.03 | false positive | | N | N | N | N | g | N | G | 0 | |
| 4232217 | T | C | 4 | 0 | 1.00 | false positive | | 0 | t | c | 0 | t | 0 | 0 | 0 | 0.PE2_1522 |

| Position | **EDI001.A** | Reference call | Uncovered positions in 50 bp | Heterozygous SNPs in 50 bp | Mean coverage ratio LS/HS | Classification | Comment | AE1175 | DIT003.B | LVC_merged | PET004.A | UNT003.A | UNT004.A | VAL001.B | WAG001.A | Homoplasy |
|---|---|---|---|---|---|---|---|---|---|---|---|---|---|---|---|---|
| 567757 | A | C | 0 | 0 | 1.00 | true positive | Probably shared ancestral | a | 0 | 0 | a | 0 | 0 | 0 | 0 | |
| 2801707 | A | G | 0 | 0 | 1.13 | true positive | unique EDI | G | G | G | G | G | g | G | 0 | |

| Position | **LVC_merged call** | Reference call | Uncovered positions in 50 bp | Heterozygous SNPs in 50 bp | Mean coverage ratio LS/HS | Classification | Comment | AE1175 | DIT003.B | EDI001.A | PET004.A | UNT003.A | UNT004.A | VAL001.B | WAG001.A | Homoplasy |
|---|---|---|---|---|---|---|---|---|---|---|---|---|---|---|---|---|
| 218506 | A | C | 0 | 1 | 3.32 | false positive | | C | C | C | C | C | C | C | 0 | |
| 481144 | T | C | 0 | 0 | 1.00 | true positive | unique LVC | C | C | C | C | C | C | C | 0 | |
| 712800 | A | C | 0 | 0 | 1.00 | true positive | unique LVC | C | C | C | C | C | C | C | 0 | |
| 1490914 | A | C | 0 | 0 | 1.00 | true positive | unique LVC | C | C | C | C | C | C | C | 0 | |
| 2515458 | T | C | 0 | 0 | 1.00 | true positive | unique LVC | C | C | C | C | C | C | C | 0 | |
| 2546208 | G | A | 10 | 0 | 1.27 | false positive | | A | A | A | A | A | A | A | 0 | |
| 2671168 | T | C | 0 | 0 | 1.21 | false positive | | C | C | C | C | 0 | C | C | 0 | |
| 2737457 | T | C | 0 | 0 | 1.00 | true positive | unique LVC | C | C | C | C | C | C | C | 0 | |
| 2924618 | A | G | 0 | 0 | 1.00 | true positive | unique LVC | G | G | G | G | G | G | G | 0 | |
| 3024601 | T | G | 0 | 0 | 1.00 | true positive | unique LVC | G | G | G | G | G | G | G | 0 | |
| 3393110 | G | A | 0 | 0 | 3.73 | false positive | | A | A | A | a | N | 0 | A | g | |
| 3665838 | C | T | 0 | 0 | 1.00 | true positive | unique LVC | T | T | T | T | T | T | T | 0 | |
| 4416487 | T | G | 0 | 0 | 1.00 | true positive | unique LVC | G | G | G | G | G | G | G | 0 | |
| 4417092 | T | G | 0 | 1 | 1.00 | false positive | | N | N | N | N | N | N | N | 0 | |
| 4563794 | A | G | 0 | 0 | 1.02 | false positive | | G | G | G | G | G | G | G | 0 | |
| 4633424 | A | G | 0 | 0 | 1.00 | true positive | unique LVC | G | G | G | G | G | G | G | 0 | |

| Position | **PET004.A call** | Reference call | Uncovered positions in 50 bp | Heterozygous SNPs in 50 bp | Mean coverage ratio LS/HS | Classification | Comment | AE1175 | DIT003.B | EDI001.A | LVC_merged | UNT003.A | UNT004.A | VAL001.B | WAG001.A | Homoplasy |
|---|---|---|---|---|---|---|---|---|---|---|---|---|---|---|---|---|
| 514923 | T | C | 0 | 1 | 1.66 | false positive | | C | C | C | C | C | C | C | c | |
| 1573356 | C | T | 0 | 1 | 2.90 | false positive | | T | T | T | N | T | T | N | N | |
| 1791420 | G | A | 0 | 0 | 1.00 | true positive | unique PET | A | a | A | 0 | A | A | A | 0 | |

| | | | | | | | | | | | | | | | | |
|---|---|---|---|---|---|---|---|---|---|---|---|---|---|---|---|---|
| 3750736 | A | G | 0 | 0 | 1.00 | true positive | shared AE, DIT, LVC, UNT, VAL | A | a | G | a | A | A | A | A | |
| 3848782 | A | T | 0 | 0 | 3.27 | false positive | | N | T | T | N | T | N | N | T | |
| 3921330 | T | C | 0 | 0 | 1.00 | true positive | unique PET | C | C | C | C | C | C | C | 0 | Y. pseudotuberculosis |
| 4198065 | G | A | 0 | 0 | 1.53 | false positive | | A | A | A | A | A | A | A | 0 | |
| 4198659 | C | A | 0 | 2 | 2.60 | false positive | | A | N | A | A | A | A | A | N | |
| 4210722 | T | A | 0 | 1 | 2.95 | false positive | | A | A | A | N | A | A | A | 0 | |
| 4412624 | G | A | 0 | 0 | 1.00 | false positive? | Feldman et al. 2016, false positive in UNT003.A | G | 0 | A | 0 | G | g | g | 0 | |
| 4644248 | T | C | 0 | 0 | 1.45 | false positive | | C | C | C | C | C | C | C | c | |
| **Position** | **UNT003.A call** | Reference call | Uncovered positions in 50 bp | Heterozygous SNPs in 50 bp | Mean coverage ratio LS/HS | Classification | Comment | AE1175 | DIT003.B | EDI001.A | LVC_merged | PET004.A | UNT004.A | VAL001.B | WAG001.A | Homoplasy |
| 549767 | C | T | 0 | 0 | 1.11 | false positive | | C | C | T | c | c | C | c | 0 | |
| 1480098 | T | G | 0 | 0 | 1.00 | true positive | probably shared AE, DIT, UNT | 0 | t | G | g | g | 0 | 0 | 0 | |
| 3179828 | A | C | 0 | 0 | 1.00 | true positive | shared AE, DIT, UNT | A | A | C | C | C | A | C | 0 | |
| 3750736 | A | G | 0 | 0 | 1.00 | true positive | shared AE, DIT, LVC, UNT, VAL | A | a | G | a | A | A | A | a | |
| 4412624 | G | A | 0 | 0 | 1.27 | false positive | | G | 0 | A | 0 | G | g | g | 0 | |
| **Position** | **UNT004.A call** | Reference call | Uncovered positions in 50 bp | Heterozygous SNPs in 50 bp | Mean coverage ratio LS/HS | Classification | Comment | AE1175 | DIT003.B | EDI001.A | LVC_merged | PET004.A | UNT003.A | VAL001.B | WAG001.A | Homoplasy |
| 225458 | G | T | 0 | 1 | 1.76 | false positive | | T | T | T | T | t | T | t | N | |
| 231350 | C | T | 24 | 1 | 1.58 | false positive | | T | T | T | N | N | T | T | g | |
| 3155190 | C | T | 0 | 1 | 1.91 | false positive | | T | T | T | T | T | T | N | 0 | |
| 3750736 | A | G | 0 | 0 | 1.00 | true positive | shared AE, DIT, LVC, UNT, VAL | A | a | G | a | A | A | A | a | |
| 3685814 | A | G | 0 | 0 | 1.59 | false positive | | G | g | G | G | g | g | G | G | |

| | | | | | | | | | | | | | | | |
|---|---|---|---|---|---|---|---|---|---|---|---|---|---|---|---|
| 3685844 | A | G | 5 | 0 | 1.75 | false positive | | G | g | G | G | G | g | g | G |
| 4199133 | A | C | 0 | 2 | 2.43 | false positive | | C | C | C | C | C | C | C | N |
| 4201886 | G | A | 0 | 1 | 1.75 | false positive | | G | G | G | G | G | G | G | G |

| **Position** | **VAL001.B call** | Reference call | Uncovered positions in 50 bp | Heterozygous SNPs in 50 bp | Mean coverage ratio LS/HS | Classification | Comment | AE1175 | DIT003.B | EDI001.A | LVC_merged | PET004.A | UNT003.A | UNT004.A | WAG001.A | Homoplasy |
|---|---|---|---|---|---|---|---|---|---|---|---|---|---|---|---|---|
| 718827 | C | T | 21 | 0 | 1.32 | false positive | | c | c | c | c | c | c | c | 0 | Branch 2 |
| 1361842 | T | C | 0 | 0 | 1.00 | true positive | unique VAL | C | C | C | C | C | C | C | 0 | |
| 2041294 | G | T | 22 | 0 | 1.00 | false positive | | T | T | T | 0 | T | t | t | 0 | |
| 2253178 | A | C | 0 | 0 | 1.00 | true positive | unique VAL | C | C | C | C | C | C | C | 0 | |
| 2303798 | C | T | 16 | 0 | 1.92 | false positive | | T | N | T | N | N | N | N | N | |
| 2732804 | G | C | 0 | 0 | 1.12 | false positive | | C | C | C | C | C | c | c | 0 | |
| 2911604 | G | A | 0 | 0 | 1.00 | true positive | unique VAL | A | A | A | A | A | A | A | 0 | |
| 3623563 | A | G | 0 | 1 | 1.07 | false positive | | N | N | N | N | 0 | N | a | 0 | |
| 3750736 | A | G | 5 | 0 | 1.00 | true positive? | shared AE, DIT, LVC, UNT, VAL | A | a | G | a | A | A | A | a | |
| 3890928 | G | C | 3 | 0 | 1.17 | false positive | | N | G | N | N | N | g | A | 0 | |
| 4256445 | G | A | 0 | 0 | 1.10 | false positive | | 0 | A | A | 0 | A | A | N | 0 | |
| 4635136 | A | C | 0 | 1 | 1.17 | false positive | | N | C | c | N | c | a | G | 0 | |
| 4648185 | A | G | 0 | 0 | 1.03 | false positive | | G | G | G | G | G | G | c | G | |

**Table S6:** True SNPs identified in the genomes of Unterthürheim (UNT), Dittenheim (DIT), Valencia (VAL), Petting (PET) and Lunel-Viel (LVC). For all identified unique SNPs including false positives, see Table S4. *This SNP appears in DIT in only one read (below our SNP calling threshold of min. 3 reads).

| Position on chromosome CO92 | Genome | Ref | SNP | SNP type | Codon change | AA change | Gene ID | Gene name | Function |
|---|---|---|---|---|---|---|---|---|---|
| 1480098 | DIT, UNT* | G | T | Intergenic | | | | | |
| 2801707 | EDI | G | A | Synonymous | acC/acT | T/T | YPO2493 | | Dioxygenase subunit alpha |
| 481144 | LVC | C | T | Synonymous | Ctg/Ttg | L/L | YPO0457 | creA | Hypothetical protein |
| 712800 | LVC | C | A | Synonymous | cgC/cgA | R/R | YPO0653 | glnE | Bifunctional glutamine-synthetase adenylyltransferase/deadenyltransferase |
| 1490914 | LVC | C | A | Upstream: 63 bases | | | YPO1324 | | Undecaprenyl pyrophosphate phosphatase |
| 2515458 | LVC | C | T | Non-synonymous | cCg/cTg | P/L | YPO2238 | | Hypothetical protein |
| 2737457 | LVC | C | T | Non-synonymous | gCg/gTg | A/V | YPO2437 | marC | multiple drug resistance protein MarC |
| 2924618 | LVC | G | A | Synonymous | gcC/gcT | A/A | YPO2603 | rodA | Cell wall shape-determining protein |
| 3024601 | LVC | G | T | Non-synonymous | atG/atT | M/I | YPO2696 | phrB | 3',5'-Cyclic-nucleotide phosphodiesterase |
| 3665838 | LVC | T | C | Non-synonymous | aAa/aGa | K/R | YPO3285 | tyrA | Bifunctional chorismate mutase/prephenate dehydrogenase |
| 4416487 | LVC | G | T | Non-synonymous | atG/atT | M/I | YPO3929 | ppc | Phosphoenolpyruvate carboxylase |
| 4633424 | LVC | G | A | Non-synonymous | aCg/aTg | T/M | YPO4112 | | Hypothetical protein |
| 1791420 | PET | A | G | Intergenic | | | | | |
| 3921330 | PET | C | T | Non-synonymous | cGc/cAc | R/H | YPO3510 | | Hypothetical protein |
| 1361842 | VAL | C | T | Non-synonymous | gGt/gAt | G/D | YPO1209 | tyrP | Tyrosine specific transport protein |
| 2253178 | VAL | C | A | Non-synonymous | Cgc/Agc | R/S | YPO1984 | | Hypothetical protein |
| 2911604 | VAL | A | G | Non-synonymous | Agt/Ggt | S/G | YPO2588 | | ABC transport protein |
| Position on pMT1 CO92 | Genome | Ref | SNP | SNP type | Codon change | AA change | Gene ID | Gene name | Function |
| 59982 | VAL | A | C | Non-synonymous | | I/C | YPMT1.59 C | | Putative DNA-binding protein |

**Table S7:** Table with all shared SNPs that were called in the First Pandemic genomes, giving the results of the SNP evaluation for each genome separately. Position is referring to the position in the reference genome CO92, followed by the reference call. For each genome, the call at the SNP position is given (0=not covered, N=heterozygous, lower case letters=less than 3 reads coverage). The classification as false or true positive is based on the preceding parameters regarding a 50 bp window surrounding the SNP: Only SNPs with no uncovered position, no heterozygous SNP and an equal mean coverage with high and low stringent mapping (ratio of 1.00) are classified as a candidates for true positive SNPs. SNPs that are supported by less than half of the genomes (<4, last column) are excluded as inconclusive (highlighted in yellow). The measurements EDI001.A are shown for completeness (grey) but not taken into consideration, since the method is not applicable to non-UDG libraries. Although the WAG001.A sample was not included in this analysis due to low coverage, all positions were checked for a rough phylogenetic classification. Similarly, the pre-Justinianic DA101 sample was checked for all positions as well.

| Position | Reference call | AE1175 | | | | DIT003.B | | | | EDI001.A | | | | UNT003.A | | | | UNT004.A | | | | PET004.A | | | | VAL001.B | | | | LVC_merged | | | | DA101 call | WAG001.A call | No. of genomes passing criteria |
|---|---|---|---|---|---|---|---|---|---|---|---|---|---|---|---|---|---|---|---|---|---|---|---|---|---|---|---|---|---|---|---|---|---|---|---|---|
| | | SNP Call | Uncovered positions in 50 bp | Heterozygous SNPs in 50 bp | Mean coverage ratio LS/HS | SNP Call | Uncovered positions in 50 bp | Heterozygous SNPs in 50 bp | Mean coverage ratio LS/HS | SNP Call | Uncovered positions in 50 bp | Heterozygous SNPs in 50 bp | Mean coverage ratio LS/HS | SNP Call | Uncovered positions in 50 bp | Heterozygous SNPs in 50 bp | Mean coverage ratio LS/HS | SNP Call | Uncovered positions in 50 bp | Heterozygous SNPs in 50 bp | Mean coverage ratio LS/HS | SNP Call | Uncovered positions in 50 bp | Heterozygous SNPs in 50 bp | Mean coverage ratio LS/HS | SNP Call | Uncovered positions in 50 bp | Heterozygous SNPs in 50 bp | Mean coverage ratio LS/HS | SNP Call | Uncovered positions in 50 bp | Heterozygous SNPs in 50 bp | Mean coverage ratio LS/HS | | | |
| 86824 | A | G | 0 | 0 | 1 | 0 | 43 | 0 | 1 | g | 0 | 1 | 1.33 | g | 0 | 0 | 1 | g | 0 | 0 | 1 | G | 0 | 0 | 1 | 0 | 48 | 0 | 1 | 0 | 48 | 0 | 1 | A | 0 | 4 |
| 189912 | A | G | 0 | 0 | 1.03 | G | 0 | 0 | 1 | G | 0 | 1 | 1.38 | G | 0 | 0 | 1 | G | 0 | 0 | 1 | G | 0 | 0 | 1 | G | 0 | 0 | 1 | G | 0 | 0 | 1 | A | 0 | 6 |
| 260148 | C | T | 0 | 0 | 1 | T | 0 | 0 | 1 | T | 0 | 0 | 1.32 | T | 0 | 0 | 1 | T | 0 | 0 | 1 | t | 0 | 0 | 1 | t | 0 | 0 | 1 | t | 6 | 0 | 1.3 | T | 0 | 5 |
| 271114 | C | A | 0 | 0 | 1 | A | 0 | 1 | 1.26 | A | 0 | 0 | 1.32 | A | 0 | 0 | 1 | A | 0 | 0 | 1.23 | A | 0 | 0 | 1 | A | 0 | 0 | 1 | a | 0 | 0 | 1.7 | C | a | 4 |
| 485976 | C | T | 0 | 0 | 1 | T | 0 | 0 | 1 | T | 0 | 0 | 1.12 | T | 0 | 0 | 1 | T | 0 | 0 | 1 | t | 1 | 0 | 1 | T | 0 | 0 | 1 | T | 0 | 0 | 1 | C | 0 | 6 |
| 557841 | C | T | 0 | 0 | 1 | T | 0 | 0 | 1.03 | T | 0 | 0 | 1.18 | T | 0 | 0 | 1 | T | 0 | 0 | 1 | t | 0 | 0 | 1 | T | 0 | 0 | 1 | T | 0 | 0 | 1 | C | 0 | 6 |
| 727741 | G | A | 0 | 0 | 1 | A | 0 | 0 | 1 | A | 0 | 0 | 1.49 | A | 0 | 0 | 1 | a | 0 | 0 | 1.28 | a | 0 | 0 | 1 | A | 0 | 0 | 1 | A | 0 | 0 | 1 | G | a | 6 |
| 779365 | C | T | 0 | 0 | 1 | T | 0 | 0 | 1 | T | 0 | 0 | 1.04 | T | 0 | 0 | 1 | T | 0 | 0 | 1 | T | 0 | 0 | 1 | T | 0 | 0 | 1 | T | 0 | 0 | 1 | C | 0 | 7 |
| 898980 | A | T | 0 | 0 | 1 | t | 0 | 0 | 1 | T | 0 | 0 | 1.31 | T | 0 | 0 | 1.12 | t | 0 | 0 | 1 | t | 0 | 0 | 1 | t | 0 | 0 | 1 | T | 0 | 0 | 1 | A | 0 | 6 |
| 1067966 | C | A | 0 | 0 | 1 | A | 0 | 0 | 1 | A | 0 | 0 | 1.10 | A | 0 | 0 | 1 | A | 0 | 0 | 1.09 | A | 0 | 0 | 1 | A | 0 | 0 | 1 | A | 0 | 0 | 1 | C | 0 | 6 |
| 1211729 | A | C | 0 | 0 | 1 | C | 0 | 0 | 1 | C | 0 | 0 | 1.15 | C | 0 | 0 | 1 | C | 0 | 0 | 1 | C | 0 | 0 | 1.08 | C | 0 | 0 | 1.04 | C | 0 | 0 | 1 | A | 0 | 5 |
| 1296743 | C | T | 0 | 0 | 1 | T | 0 | 0 | 1.08 | T | 0 | 0 | 1.33 | T | 0 | 0 | 1 | T | 0 | 0 | 1.06 | T | 0 | 0 | 1 | T | 0 | 0 | 1.09 | T | 0 | 0 | 1 | C | t | 3 |

| | | | | | | | | | | | | | | | | | | | | | | | | | | | | | | | | | | | | |
|---|---|---|---|---|---|---|---|---|---|---|---|---|---|---|---|---|---|---|---|---|---|---|---|---|---|---|---|---|---|---|---|---|---|---|---|---|
| 1387701 | C | T | 0 | 0 | 1 | t | 9 | 0 | 1 | T | 0 | 0 | 1.15 | t | 0 | 0 | 1 | T | 0 | 0 | 1 | T | 0 | 0 | 1 | T | 0 | 0 | 1 | 0 | 31 | 0 | 1 | c | 0 | 5 |
| 1387756 | A | G | 0 | 0 | 1 | 0 | 42 | 0 | 1 | G | 0 | 0 | 1.26 | g | 11 | 0 | 1 | G | 0 | 0 | 1 | 0 | 27 | 0 | 1 | 0 | 35 | 0 | 1 | g | 20 | 0 | 1 | A | 0 | 2 |
| 1413031 | C | A | 0 | 0 | 1 | A | 0 | 0 | 1 | A | 0 | 0 | 1.36 | a | 0 | 0 | 1 | A | 0 | 0 | 1 | a | 5 | 0 | 1 | a | 0 | 0 | 1 | a | 2 | 0 | 1 | C | 0 | 5 |
| 1434752 | C | A | 0 | 0 | 1.03 | a | 0 | 0 | 1 | A | 0 | 0 | 1.05 | A | 0 | 0 | 1 | A | 0 | 0 | 1 | A | 0 | 0 | 1 | A | 0 | 0 | 1 | A | 0 | 0 | 1 | C | 0 | 6 |
| 1489055 | C | T | 0 | 0 | 1 | T | 0 | 0 | 1 | T | 0 | 0 | 1.36 | T | 0 | 0 | 1 | T | 0 | 0 | 1 | T | 0 | 0 | 1 | T | 0 | 0 | 1 | N | 0 | 0 | 1 | C | t | 6 |
| 1530658 | C | A | 0 | 0 | 1 | A | 0 | 0 | 1 | A | 0 | 0 | 1.27 | A | 0 | 0 | 1 | a | 0 | 0 | 1 | a | 0 | 0 | 1 | a | 0 | 0 | 1 | a | 11 | 0 | 1 | C | 0 | 6 |
| 1609461 | T | C | 0 | 0 | 1 | C | 0 | 0 | 1.1 | C | 0 | 0 | 1.32 | C | 0 | 0 | 1 | C | 0 | 0 | 1 | C | 0 | 0 | 1 | C | 0 | 0 | 1 | C | 0 | 0 | 1 | T | 0 | 6 |
| 1754708 | C | T | 0 | 0 | 1 | T | 0 | 0 | 1 | T | 0 | 0 | 1.19 | T | 0 | 0 | 1 | 0 | 25 | 0 | 1 | T | 0 | 0 | 1 | t | 0 | 0 | 1 | T | 0 | 0 | 1 | C | 0 | 6 |
| 1868678 | G | T | 0 | 0 | 1 | T | 0 | 0 | 1.15 | T | 0 | 0 | 1.57 | T | 0 | 0 | 1 | T | 0 | 0 | 1 | T | 0 | 0 | 1 | T | 0 | 0 | 1 | T | 0 | 0 | 1 | G | 0 | 6 |
| 1956162 | T | C | 0 | 0 | 1 | 0 | 41 | 0 | 1 | C | 0 | 0 | 1.37 | C | 0 | 0 | 1 | C | 0 | 0 | 1 | C | 0 | 0 | 1 | C | 0 | 0 | 1 | 0 | 30 | 0 | 1 | T | c | 5 |
| 2092152 | C | T | 0 | 0 | 1 | t | 0 | 0 | 1 | T | 0 | 0 | 1.23 | T | 0 | 0 | 1 | 0 | 9 | 0 | 1 | t | 15 | 0 | 1 | t | 0 | 0 | 1 | 0 | 12 | 0 | 1 | C | 0 | 4 |
| 2097520 | G | T | 0 | 0 | 1 | T | 0 | 0 | 1 | T | 0 | 1 | 1.16 | T | 0 | 0 | 1 | t | 0 | 0 | 1 | T | 0 | 0 | 1 | T | 0 | 0 | 1 | N | 7 | 1 | 1 | G | t | 6 |
| 2352174 | T | G | 0 | 0 | 1 | G | 0 | 0 | 1 | g | 0 | 0 | 1.12 | G | 0 | 0 | 1 | G | 0 | 0 | 1 | G | 0 | 0 | 1 | G | 0 | 0 | 1 | G | 0 | 0 | 1 | T | 0 | 6 |
| 2419529 | G | A | 0 | 0 | 1 | A | 0 | 0 | 1 | a | 0 | 0 | 1.21 | A | 0 | 0 | 1 | A | 0 | 0 | 1 | A | 0 | 0 | 1 | A | 0 | 0 | 1.1 | A | 0 | 0 | 1 | G | a | 5 |
| 2495165 | C | A | 0 | 0 | 1 | 0 | 50 | 0 | 1 | A | 0 | 0 | 1.16 | a | 14 | 0 | 1 | A | 0 | 0 | 1 | a | 0 | 0 | 1 | 0 | 50 | 0 | NA | a | 12 | 0 | 1 | C | 0 | 3 |
| 2725715 | C | T | 0 | 0 | 1 | t | 14 | 0 | 1 | T | 0 | 0 | 1.27 | T | 0 | 0 | 1 | T | 0 | 0 | 1 | T | 0 | 0 | 1 | T | 0 | 0 | 1 | T | 0 | 0 | 1 | T | 0 | 6 |
| 2753572 | A | T | 0 | 0 | 1 | T | 0 | 0 | 1 | T | 0 | 0 | 1.09 | t | 0 | 0 | 1 | T | 0 | 0 | 1 | T | 0 | 0 | 1.07 | T | 0 | 0 | 1 | T | 0 | 0 | 1 | A | 0 | 6 |
| 2977542 | C | A | 0 | 0 | 1 | A | 0 | 0 | 1 | A | 0 | 1 | 1.23 | A | 0 | 0 | 1 | A | 0 | 0 | 1 | A | 0 | 0 | 1 | A | 0 | 0 | 1.08 | A | 0 | 0 | 1.1 | A | 0 | 5 |
| 3078807 | C | A | 0 | 0 | 1.05 | A | 0 | 0 | 1.03 | A | 0 | 0 | 1.04 | A | 0 | 0 | 1 | A | 0 | 0 | 1 | A | 0 | 0 | 1 | A | 0 | 0 | 1 | A | 0 | 0 | 1 | C | 0 | 5 |
| 3274298 | A | T | 0 | 0 | 1 | T | 0 | 0 | 1 | T | 0 | 0 | 1.60 | T | 0 | 0 | 1 | T | 0 | 0 | 1 | T | 0 | 0 | 1 | T | 0 | 0 | 1 | T | 0 | 0 | 1 | t | 0 | 7 |
| 3360963 | A | C | 0 | 0 | 1.08 | C | 0 | 0 | 1.27 | C | 0 | 0 | 1.64 | 0 | 16 | 0 | 1.56 | 0 | 43 | 0 | 1 | C | 0 | 0 | 1 | C | 0 | 0 | 1.07 | C | 0 | 0 | 1.1 | A | 0 | 1 |
| 3360984 | C | T | 0 | 0 | 1.06 | T | 0 | 0 | 1.28 | T | 0 | 0 | 1.58 | t | 16 | 0 | 1.63 | t | 22 | 0 | 1 | T | 0 | 0 | 1 | T | 0 | 0 | 1.17 | T | 0 | 0 | 1.1 | C | 0 | 1 |
| 3398153 | G | A | 0 | 0 | 1 | A | 0 | 0 | 1.05 | a | 0 | 0 | 1.47 | A | 0 | 0 | 1 | a | 1 | 0 | 1 | A | 0 | 0 | 1 | A | 0 | 0 | 1 | a | 0 | 0 | 1 | G | 0 | 5 |
| 3409414 | T | C | 0 | 0 | 1 | c | 6 | 0 | 1 | C | 0 | 0 | 1.01 | C | 0 | 0 | 1 | c | 7 | 0 | 1 | 0 | 26 | 0 | 1 | 0 | 26 | 0 | 1 | 0 | 12 | 0 | 1.1 | T | 0 | 2 |
| 3500922 | T | G | 0 | 0 | 1 | G | 0 | 0 | 1 | G | 0 | 0 | 1.49 | G | 0 | 0 | 1 | g | 0 | 0 | 1 | G | 0 | 0 | 1 | G | 0 | 0 | 1 | G | 0 | 0 | 1 | T | 0 | 7 |
| 3535148 | G | T | 0 | 0 | 1 | T | 0 | 0 | 1 | T | 0 | 0 | 1.00 | T | 0 | 0 | 1 | T | 0 | 0 | 1 | T | 0 | 0 | 1 | T | 0 | 0 | 1 | T | 0 | 0 | 1 | G | 0 | 7 |
| 3560088 | G | A | 0 | 0 | 1 | A | 0 | 0 | 1 | A | 0 | 0 | 1.13 | A | 0 | 0 | 1 | A | 0 | 1 | 1 | A | 0 | 0 | 1 | A | 0 | 0 | 1 | A | 0 | 0 | 1 | G | 0 | 6 |
| 3568597 | C | T | 0 | 0 | 1 | T | 0 | 0 | 1 | T | 0 | 0 | 1.22 | T | 0 | 0 | 1.05 | T | 0 | 0 | 1 | T | 0 | 0 | 1 | T | 0 | 0 | 1 | T | 0 | 0 | 1 | C | 0 | 6 |
| 3755861 | C | T | 0 | 0 | 1 | 0 | 29 | 0 | 1 | T | 0 | 0 | 1.01 | t | 15 | 0 | 1 | t | 7 | 1 | 1 | t | 1 | 0 | 1 | 0 | 37 | 0 | 1 | t | 22 | 0 | 1 | C | 0 | 1 |
| 3843195 | C | A | 0 | 0 | 1 | A | 0 | 0 | 1 | A | 0 | 1 | 1.14 | A | 0 | 0 | 1 | A | 0 | 0 | 1 | A | 0 | 0 | 1 | A | 0 | 0 | 1 | A | 0 | 0 | 1 | C | 0 | 7 |

| | | | | | | | | | | | | | | | | | | | | | | | | | | | | | | | | | | | | |
|---|---|---|---|---|---|---|---|---|---|---|---|---|---|---|---|---|---|---|---|---|---|---|---|---|---|---|---|---|---|---|---|---|---|---|---|---|
| 3892488 | C | T | 0 | 0 | 1 | T | 0 | 0 | 1 | T | 0 | 0 | 1.36 | T | 0 | 0 | 1 | T | 0 | 0 | 1 | T | 0 | 0 | 1.19 | T | 0 | 0 | 1 | T | 0 | 0 | 1 | C | 0 | 6 |
| 4066494 | C | T | 0 | 0 | 1 | t | 17 | 0 | 1.37 | T | 0 | 0 | 1.28 | T | 0 | 0 | 1 | t | 0 | 0 | 1 | 0 | 16 | 0 | 1 | T | 0 | 0 | 1 | T | 2 | 0 | 1.2 | C | 0 | 4 |
| 4307755 | G | A | 0 | 0 | 1 | A | 0 | 0 | 1 | A | 0 | 0 | 1.25 | A | 0 | 0 | 1 | A | 0 | 0 | 1 | A | 0 | 0 | 1 | A | 0 | 0 | 1 | A | 0 | 0 | 1 | G | 0 | 7 |
| 4423366 | G | A | 0 | 0 | 1 | A | 0 | 0 | 1.04 | A | 0 | 0 | 1.62 | A | 0 | 0 | 1 | A | 0 | 0 | 1 | A | 0 | 0 | 1.07 | A | 0 | 0 | 1 | A | 0 | 0 | 1 | G | 0 | 5 |
| 4460688 | C | T | 0 | 0 | 1 | T | 0 | 0 | 1 | T | 0 | 0 | 1.50 | T | 0 | 0 | 1 | T | 0 | 0 | 1 | T | 0 | 0 | 1 | T | 0 | 0 | 1 | T | 0 | 0 | 1 | T | 0 | 7 |
| 4465967 | C | A | 0 | 0 | 1.04 | A | 0 | 0 | 1 | A | 0 | 0 | 1.71 | A | 0 | 0 | 1.04 | A | 0 | 0 | 1 | A | 0 | 0 | 1 | A | 0 | 0 | 1 | A | 0 | 0 | 1 | c | 0 | 5 |
| 4628496 | C | A | 0 | 0 | 1 | A | 0 | 0 | 1 | A | 0 | 0 | 1.52 | A | 0 | 0 | 1 | a | 0 | 0 | 1 | A | 0 | 0 | 1 | A | 0 | 0 | 1 | A | 0 | 0 | 1 | C | 0 | 7 |
| 4629169 | G | A | 0 | 0 | 1.03 | A | 0 | 0 | 1 | A | 0 | 0 | 1.12 | A | 0 | 0 | 1 | A | 0 | 0 | 1.12 | A | 0 | 0 | 1 | A | 0 | 0 | 1 | A | 0 | 0 | 1.2 | G | 0 | 4 |

**Table S8:** List of all modern *Y. pestis* genomes with accession number, sample origin and the corresponding publication (for full references see main text).

| **Modern strains** | | | |
|---|---|---|---|
| **Strain ID** | **Accession No.** | **Origin** | **Publication** |
| 0.ANT1a_42013 | ADPG00000000 | China | Cui et al. 2013 |
| 0.ANT1b_CMCC49003 | ADQX00000000 | China | Cui et al. 2013 |
| 0.ANT1c_945 | ADPV00000000 | China | Cui et al. 2013 |
| 0.ANT1d_164 | ADOW00000000 | China | Cui et al. 2013 |
| 0.ANT1e_CMCC8211 | ADRD00000000 | China | Cui et al. 2013 |
| 0.ANT1f_42095 | ADPJ00000000 | China | Cui et al. 2013 |
| 0.ANT1g_CMCC42007 | ADQV00000000 | China | Cui et al. 2013 |
| 0.ANT1h_CMCC43032 | ADQW00000000 | China | Cui et al. 2013 |
| 0.ANT2_B42003004 | AAYU00000000 | China | Cui et al. 2013 |
| 0.ANT2a_2330 | ADOY00000000 | China | Cui et al. 2013 |
| 0.ANT3_231 | JMUF00000000 | FSU | Eroshenko et al. 2017 |
| 0.ANT3_790 | CP006806 | FSU | Zhgenti et al. 2015 |
| 0.ANT3_A-1486 | LYMP00000000 | FSU | Eroshenko et al. 2017 |
| 0.ANT3a_CMCC38001 | ADQU00000000 | China | Cui et al. 2013 |
| 0.ANT3b_A1956001 | ADPX00000000 | China | Cui et al. 2013 |
| 0.ANT3c_42082 | ADPH00000000 | China | Cui et al. 2013 |
| 0.ANT3d_CMCC21106 | ADQP00000000 | China | Cui et al. 2013 |
| 0.ANT3e_42091b | ADPI00000000 | China | Cui et al. 2013 |
| 0.ANT5_262 | QAGF00000000 | FSU | Kutyrev et al. 2018 |
| 0.ANT5_5M | LYMQ00000000 | FSU | Kutyrev et al. 2018 |
| 0.ANT5_A-1691 | LYMQ00000000 | FSU | Eroshenko et al. 2017 |
| 0.ANT5_A-1836 | LYOL00000000 | FSU | Eroshenko et al. 2017 |
| 0.PE2_1412 | CP006783 | FSU | Zhgenti et al. 2015 |
| 0.PE2_1413 | CP006762 | FSU | Zhgenti et al. 2015 |
| 0.PE2_14735 | AYLS00000000 | FSU | Zhgenti et al. 2015 |
| 0.PE2_1522 | CP006758 | FSU | Zhgenti et al. 2015 |
| 0.PE2_1670 | CP006806 | FSU | Zhgenti et al. 2015 |
| 0.PE2_3067 | CP006754 | FSU | Zhgenti et al. 2015 |
| 0.PE2_3544 | LZNH00000000 | FSU | Kutyrev et al. 2018 |
| 0.PE2_3551 | MBSJ00000000 | FSU | Kutyrev et al. 2018 |
| 0.PE2_3770 | CP006751 | FSU | Zhgenti et al. 2015 |
| 0.PE2_835_BPC | LYOJ00000000 | FSU | Kutyrev et al. 2018 |
| 0.PE2_8787 | CP006748 | FSU | Zhgenti et al. 2015 |
| 0.PE2_C-197 | LIYX00000000 | FSU | Kislichkina et al. 2015 |
| 0.PE2_C-235 | LIYY00000000 | FSU | Kislichkina et al. 2015 |
| 0.PE2_C-267 | LIYZ00000000 | FSU | Kislichkina et al. 2015 |
| 0.PE2_C-290 | LIYU00000000 | FSU | Kislichkina et al. 2015 |
| 0.PE2_C-291 | LIZC00000000 | FSU | Kislichkina et al. 2015 |
| 0.PE2_C-346 | LIZE00000000 | FSU | Kislichkina et al. 2015 |
| 0.PE2_C-359 | LIZB00000000 | FSU | Kislichkina et al. 2015 |
| 0.PE2_C-370 | MIDX00000000 | FSU | Kislichkina et al. 2017 |
| 0.PE2_C-535 | MIDY00000000 | FSU | Kislichkina et al. 2017 |
| 0.PE2_C-537 | LIYP00000000 | FSU | Kislichkina et al. 2015 |
| 0.PE2_C-590 | LIYQ00000000 | FSU | Kislichkina et al. 2015 |
| 0.PE2_C-666 | LIZF00000000 | FSU | Kislichkina et al. 2015 |
| 0.PE2_C-678 | MIDZ00000000 | FSU | Kislichkina et al. 2017 |
| 0.PE2_C-700 | MIEA00000000 | FSU | Kislichkina et al. 2017 |
| 0.PE2_C-712 | MTZW00000000 | FSU | Kislichkina et al. 2017 |
| 0.PE2_C-739 | MTZX00000000 | FSU | Kislichkina et al. 2017 |
| 0.PE2_C-741 | LPTX00000000 | FSU | Kutyrev et al. 2018 |
| 0.PE2_C-746 | MTZY00000000 | FSU | Kislichkina et al. 2017 |
| 0.PE2_C-824 | MTZZ00000000 | FSU | Kislichkina et al. 2017 |

| | | | |
|---|---|---|---|
| 0.PE2_KM874 | LZTG00000000 | FSU | Kutyrev et al. 2018 |
| 0.PE2_M-986 | LYMO00000000 | FSU | Kutyrev et al. 2018 |
| 0.PE2_PEST-F | NC_009381 | FSU | Cui et al. 2013 |
| 0.PE2b_G8786 | ADSG00000000 | FSU | Cui et al. 2013 |
| 0.PE4_5307-Gis | LIYS00000000 | FSU | Kislichkina et al. 2015 |
| 0.PE4_A-1804 | LIYW00000000 | FSU | Kislichkina et al. 2015 |
| 0.PE4_A-1807 | LIYT00000000 | FSU | Kislichkina et al. 2015 |
| 0.PE4_A-513 | LIZA00000000 | FSU | Kislichkina et al. 2015 |
| 0.PE4_I-3134 | LIYR00000000 | FSU | Kislichkina et al. 2015 |
| 0.PE4_I-3442 | NHYH00000000 | FSU | Kislichkina et al. 2018a |
| 0.PE4_I-3443 | MIED00000000 | FSU | Kislichkina et al. 2018a |
| 0.PE4_I-3446 | NHYI00000000 | FSU | Kislichkina et al. 2018a |
| 0.PE4_I-3447 | MIEE00000000 | FSU | Kislichkina et al. 2018a |
| 0.PE4_I-3455 | LIYV00000000 | FSU | Kislichkina et al. 2015 |
| 0.PE4_I-3515 | NHYJ00000000 | FSU | Kislichkina et al. 2018a |
| 0.PE4_I-3516 | NHMW00000000 | FSU | Kislichkina et al. 2018a |
| 0.PE4_I-3517 | NHMX00000000 | FSU | Kislichkina et al. 2018a |
| 0.PE4_I-3518 | NHMY00000000 | FSU | Kislichkina et al. 2018a |
| 0.PE4_I-3519 | NHMZ00000000 | FSU | Kislichkina et al. 2018a |
| 0.PE4_M0000002 | ADST00000000 | China | Cui et al. 2013 |
| 0.PE4_Microtus91001 | NC_005810 | China | Cui et al. 2013 |
| 0.PE4a_B1313 | LYMS00000000 | FSU | Kutyrev et al. 2018 |
| 0.PE4a_I-2751-55 | LYCL00000000 | FSU | Kutyrev et al. 2018 |
| 0.PE4a_I-2998 | LYMR00000000 | FSU | Kutyrev et al. 2018 |
| 0.PE4Aa_12 | ADOV00000000 | China | Cui et al. 2013 |
| 0.PE4Ab_9 | ADPT00000000 | China | Cui et al. 2013 |
| 0.PE4Ba_PestoidesA | ACNT00000000 | FSU | Cui et al. 2013 |
| 0.PE4Ca_CMCCN010025 | ADRT00000000 | China | Cui et al. 2013 |
| 0.PE4Cc_CMCC18019 | ADQO00000000 | China | Cui et al. 2013 |
| 0.PE4Cd_CMCC93014 | ADRM00000000 | China | Cui et al. 2013 |
| 0.PE4Ce_CMCC91090 | ADRJ00000000 | China | Cui et al. 2013 |
| 0.PE4h_A-1249 | LYMN00000000 | FSU | Eroshenko et al. 2017 |
| 0.PE4m_I-3086 | LZNY00000000 | Mongolia | Kutyrev et al. 2018 |
| 0.PE4t_A-1815 | LPTY00000000 | FSU | Eroshenko et al. 2017 |
| 0.PE5_I-2231 | PVLX00000000 | Mongolia | Kislichkina et al. 2018b |
| 0.PE5_I-2236 | PVLZ00000000 | Mongolia | Kislichkina et al. 2018b |
| 0.PE5_I-2238 | PVLX00000000 | Mongolia | Kislichkina et al. 2018b |
| 0.PE5_I-2239 | LIZD00000000 | Mongolia | Kislichkina et al. 2015 |
| 0.PE5_I-2422a | LIZG00000000 | Mongolia | Kislichkina et al. 2015 |
| 0.PE5_I-2422b | QANK00000000 | Mongolia | Kutyrev et al. 2018 |
| 0.PE5_I-2457 | PVMB00000000 | Mongolia | Kislichkina et al. 2018b |
| 0.PE5_I-3189 | LIYO00000000 | Mongolia | Kislichkina et al. 2015 |
| 0.PE5_I-3190 | PVLY00000000 | Mongolia | Kislichkina et al. 2018b |
| 0.PE7b_620024 | ADPM00000000 | China | Cui et al. 2013 |
| 1.ANT1_Antiqua | NC_008150 | Congo | Cui et al. 2013 |
| 1.ANT1_UG05-0454 | AAYR00000000 | Uganda | Cui et al. 2013 |
| 1.IN1a_CMCC11001 | ADQK00000000 | China | Cui et al. 2013 |
| 1.IN1b_780441 | ADPS00000000 | China | Cui et al. 2013 |
| 1.IN1c_K21985002 | ADSS00000000 | China | Cui et al. 2013 |
| 1.IN2a_CMCC640047 | ADRA00000000 | China | Cui et al. 2013 |
| 1.IN2b_30017 | ADPC00000000 | China | Cui et al. 2013 |
| 1.IN2c_CMCC31004 | ADQR00000000 | China | Cui et al. 2013 |
| 1.IN2d_C1975003 | ADPZ00000000 | China | Cui et al. 2013 |
| 1.IN2e_C1989001 | ADQB00000000 | China | Cui et al. 2013 |
| 1.IN2f_710317 | ADPP00000000 | China | Cui et al. 2013 |
| 1.IN2g_CMCC05013 | ADQF00000000 | China | Cui et al. 2013 |
| 1.IN2h_5 | ADPK00000000 | China | Cui et al. 2013 |
| 1.IN2i_CMCC10012 | ADQG00000000 | China | Cui et al. 2013 |

| | | | |
|---|---|---|---|
| 1.IN2j_CMCC27002 | ADQQ00000000 | China | Cui et al. 2013 |
| 1.IN2k_970754 | ADPW00000000 | China | Cui et al. 2013 |
| 1.IN2l_D1991004 | ADRX00000000 | China | Cui et al. 2013 |
| 1.IN2m_D1964002b | ADRV00000000 | China | Cui et al. 2013 |
| 1.IN2n_CMCC02041 | ADQC00000000 | China | Cui et al. 2013 |
| 1.IN2o_CMCC03001 | ADQD00000000 | China | Cui et al. 2013 |
| 1.IN2p_D1982001 | ADRW00000000 | China | Cui et al. 2013 |
| 1.IN2q_D1964001 | ADRU00000000 | China | Cui et al. 2013 |
| 1.IN3a_F1954001 | ADSC00000000 | China | Cui et al. 2013 |
| 1.IN3b_E1979001 | AAYV00000000 | China | Cui et al. 2013 |
| 1.IN3c_CMCC84038b | ADRF00000000 | China | Cui et al. 2013 |
| 1.IN3d_YN1683 | ADTD00000000 | China | Cui et al. 2013 |
| 1.IN3e_YN472 | ADTH00000000 | China | Cui et al. 2013 |
| 1.IN3f_YN1065 | ADTC00000000 | China | Cui et al. 2013 |
| 1.IN3g_E1977001 | ADRY00000000 | China | Cui et al. 2013 |
| 1.IN3h_CMCC84033 | ADRE00000000 | China | Cui et al. 2013 |
| 1.IN3i_CMCC84046 | ADRG00000000 | China | Cui et al. 2013 |
| 1.ORI1_CA88 | ABCD00000000 | USA | Cui et al. 2013 |
| 1.ORI1_CO92 | NC_003143 | USA | Cui et al. 2013 |
| 1.ORI1a_CMCC114001 | ADQL00000000 | China | Cui et al. 2013 |
| 1.ORI1b_India195 | ACNR00000000 | India | Cui et al. 2013 |
| 1.ORI1c_F1946001 | ADSB00000000 | China | Cui et al. 2013 |
| 1.ORI2_F1991016 | ABAT00000000 | China | Cui et al. 2013 |
| 1.ORI2a_YN2179 | ADTE00000000 | Myanmar | Cui et al. 2013 |
| 1.ORI2c_YN2551b | ADTF00000000 | China | Cui et al. 2013 |
| 1.ORI2d_YN2588 | ADTG00000000 | China | Cui et al. 2013 |
| 1.ORI2f_CMCC87001 | ADRH00000000 | China | Cui et al. 2013 |
| 1.ORI2g_F1984001 | ADSD00000000 | China | Cui et al. 2013 |
| 1.ORI2h_YN663 | ADTI00000000 | China | Cui et al. 2013 |
| 1.ORI2i_CMCCK100001a | ADRR00000000 | China | Cui et al. 2013 |
| 1.ORI2i_CMCCK110001b | ADRS00000000 | China | Cui et al. 2013 |
| 1.ORI3_IP275 | AAOS00000000 | Madagascar | Cui et al. 2013 |
| 1.ORI3_MG05-1020 | AAYS00000000 | Madagascar | Cui et al. 2013 |
| 1.ORI3a_EV76 | ADSA00000000 | Madagascar | Cui et al. 2013 |
| 2.ANT1_Nepal516 | ACNQ00000000 | Nepal | Cui et al. 2013 |
| 2.ANT1a_34008 | ADPD00000000 | China | Cui et al. 2013 |
| 2.ANT1b_34202 | ADPE00000000 | China | Cui et al. 2013 |
| 2.ANT2a_2 | ADOX00000000 | China | Cui et al. 2013 |
| 2.ANT2b_351001 | ADPF00000000 | China | Cui et al. 2013 |
| 2.ANT2c_CMCC347001 | ADQS00000000 | China | Cui et al. 2013 |
| 2.ANT2d_G1996006 | ADSE00000000 | China | Cui et al. 2013 |
| 2.ANT2e_G1996010 | ADSF00000000 | China | Cui et al. 2013 |
| 2.ANT2f_CMCC348002 | ADQT00000000 | China | Cui et al. 2013 |
| 2.ANT3_KM682 | LPVG00000000 | FSU | Kutyrev et al. 2018 |
| 2.ANT3a_CMCC92010 | ADRL00000000 | China | Cui et al. 2013 |
| 2.ANT3b_CMCC95001 | ADRN00000000 | China | Cui et al. 2013 |
| 2.ANT3c_CMCC96001 | ADRO00000000 | China | Cui et al. 2013 |
| 2.ANT3d_CMCC96007 | ADRP00000000 | China | Cui et al. 2013 |
| 2.ANT3e_CMCC67001 | ADRB00000000 | China | Cui et al. 2013 |
| 2.ANT3f_CMCC104003 | ADQH00000000 | China | Cui et al. 2013 |
| 2.ANT3g_CMCC51020 | ADQY00000000 | China | Cui et al. 2013 |
| 2.ANT3h_CMCC106002 | ADQI00000000 | China | Cui et al. 2013 |
| 2.ANT3i_CMCC64001 | ADQZ00000000 | China | Cui et al. 2013 |
| 2.ANT3j_H1959004 | ADSI00000000 | China | Cui et al. 2013 |
| 2.ANT3k_5761 | ADPL00000000 | Russia | Cui et al. 2013 |
| 2.ANT3l_735 | ADPR00000000 | Russia | Cui et al. 2013 |
| 2.MED0_C-627 | MBSI00000000 | FSU | Kutyrev et al. 2018 |
| 2.MED1_1045 | CP006794 | FSU | Zhgenti et al. 2015 |

| | | | |
|---|---|---|---|
| 2.MED1_1116-D | LPXS00000000 | FSU | Kutyrev et al. 2018 |
| 2.MED1_1240 | LZNI00000000 | FSU | Kutyrev et al. 2018 |
| 2.MED1_139 | QAPA00000000 | FSU | Kutyrev et al. 2018 |
| 2.MED1_173 | LQAZ00000000 | FSU | Kutyrev et al. 2018 |
| 2.MED1_1906 | LYOM00000000 | FSU | Kutyrev et al. 2018 |
| 2.MED1_244 | LZND00000000 | FSU | Kutyrev et al. 2018 |
| 2.MED1_261 | LZNG00000000 | FSU | Kutyrev et al. 2018 |
| 2.MED1_2944 | CP006792 | FSU | Zhgenti et al. 2015 |
| 2.MED1_44 | LZNF00000000 | FSU | Kutyrev et al. 2018 |
| 2.MED1_A-1763 | LQAW00000000 | FSU | Kutyrev et al. 2018 |
| 2.MED1_A-1809 | LYMF00000000 | FSU | Eroshenko et al. 2017 |
| 2.MED1_A-1825 | LYCM00000000 | FSU | Kutyrev et al. 2018 |
| 2.MED1_A-1920 | LYCO00000000 | FSU | Kutyrev et al. 2018 |
| 2.MED1_C-791 | LQAU00000000 | FSU | Kutyrev et al. 2018 |
| 2.MED1_K1973002 | AAYT00000000 | China | Cui et al. 2013 |
| 2.MED1_KIM10 | NC_004088 | Iran | Cui et al. 2013 |
| 2.MED1_KM816 | LPXU00000000 | FSU | Kutyrev et al. 2018 |
| 2.MED1_KM918 | LPQY00000000 | FSU | Kutyrev et al. 2018 |
| 2.MED1_M-1448 | LYCN00000000 | FSU | Kutyrev et al. 2018 |
| 2.MED1_M-1453 | LQAY00000000 | FSU | Kutyrev et al. 2018 |
| 2.MED1_M-1484 | LQAV00000000 | FSU | Kutyrev et al. 2018 |
| 2.MED1_M-1524 | LYCP00000000 | FSU | Kutyrev et al. 2018 |
| 2.MED1_M-1773 | LYMG00000000 | FSU | Kutyrev et al. 2018 |
| 2.MED1_M-1864 | LOHR00000000 | FSU | Kutyrev et al. 2018 |
| 2.MED1_M-519 | LQAX00000000 | FSU | Kutyrev et al. 2018 |
| 2.MED1_M-549 | LQBA00000000 | FSU | Kutyrev et al. 2018 |
| 2.MED1_M-595 | LYOH00000000 | FSU | Kutyrev et al. 2018 |
| 2.MED1_M-978 | LPXT00000000 | FSU | Kutyrev et al. 2018 |
| 2.MED1b_2506 | ADPA00000000 | China | Cui et al. 2013 |
| 2.MED1c_2654 | ADPB00000000 | China | Cui et al. 2013 |
| 2.MED1d_2504 | ADOZ00000000 | China | Cui et al. 2013 |
| 2.MED2b_91 | ADPU00000000 | China | Cui et al. 2013 |
| 2.MED2c_K11973002 | AAYT00000000 | China | Cui et al. 2013 |
| 2.MED2d_A1973001 | ADPY00000000 | China | Cui et al. 2013 |
| 2.MED2e_7338 | ADPQ00000000 | China | Cui et al. 2013 |
| 2.MED3a_J1963002 | ADSP00000000 | China | Cui et al. 2013 |
| 2.MED3b_CMCC125002b | ADQN00000000 | China | Cui et al. 2013 |
| 2.MED3c_I1969003 | ADSK00000000 | China | Cui et al. 2013 |
| 2.MED3d_J1978002 | ADSQ00000000 | China | Cui et al. 2013 |
| 2.MED3f_I1970005 | ADSL00000000 | China | Cui et al. 2013 |
| 2.MED3g_CMCC99103 | ADRQ00000000 | China | Cui et al. 2013 |
| 2.MED3h_CMCC90027 | ADRI00000000 | China | Cui et al. 2013 |
| 2.MED3i_CMCC92004 | ADRK00000000 | China | Cui et al. 2013 |
| 2.MED3j_I2001001 | ADSO00000000 | China | Cui et al. 2013 |
| 2.MED3k_CMCC12003 | ADQM00000000 | China | Cui et al. 2013 |
| 2.MED3l_I1994006 | ADSN00000000 | China | Cui et al. 2013 |
| 2.MED3m_SHAN11 | ADTA00000000 | China | Cui et al. 2013 |
| 2.MED3n_SHAN12 | ADTB00000000 | China | Cui et al. 2013 |
| 2.MED3o_I1991001 | ADSM00000000 | China | Cui et al. 2013 |
| 2.MED3p_CMCC107004 | ADQJ00000000 | China | Cui et al. 2013 |
| 3.ANT1a_7b | ADPN00000000 | China | Cui et al. 2013 |
| 3.ANT1b_CMCC71001 | ADRC00000000 | China | Cui et al. 2013 |
| 3.ANT1c_C1976001 | ADQA00000000 | China | Cui et al. 2013 |
| 3.ANT1d_71021 | ADPO00000000 | China | Cui et al. 2013 |
| 3.ANT2a_MGJZ6 | ADSX00000000 | Mongolia | Cui et al. 2013 |
| 3.ANT2b_MGJZ7 | ADSY00000000 | Mongolia | Cui et al. 2013 |
| 3.ANT2c_MGJZ9 | ADSZ00000000 | Mongolia | Cui et al. 2013 |
| 3.ANT2d_MGJZ11 | ADSU00000000 | Mongolia | Cui et al. 2013 |

| | | | |
|---|---|---|---|
| 3.ANT2e_MGJZ3 | ADSW00000000 | Mongolia | Cui et al. 2013 |
| 4.ANT_1454 | LZNC00000000 | FSU | Kutyrev et al. 2018 |
| 4.ANT_338 | LZNX00000000 | FSU | Kutyrev et al. 2018 |
| 4.ANT_517 | LYMH00000000 | FSU | Kutyrev et al. 2018 |
| 4.ANT_KM932 | LZNE00000000 | FSU | Kutyrev et al. 2018 |
| 4.ANT_M-1944 | LYOK00000000 | FSU | Kutyrev et al. 2018 |
| 4.ANT1a_MGJZ12 | ADSV00000000 | Mongolia | Cui et al. 2013 |
| **Ancient strains** | | | |
| **Strain ID** | **Accession No.** | **Origin** | **Publication** |
| DA101 | PRJEB25891 | Kyrgyzstan | Daamgard et al. 2018 |
| Altenerding | PRJEB14851 | Germany | Feldman et al. 2016 |
| 8124/8291/11972 | SRR341961, SRR341962, SRR341963 | United Kingdom | Bos et al. 2011 |
| Ellwangen | PRJEB13664 | Germany | Spyrou et al. 2016 |
| OBS107 | PRJEB12163 | France | Bos et al. 2016 |
| OBS110 | PRJEB12163 | France | Bos et al. 2016 |
| OBS116 | PRJEB12163 | France | Bos et al. 2016 |
| OBS124 | PRJEB12163 | France | Bos et al. 2016 |
| OBS137 | PRJEB12163 | France | Bos et al. 2016 |

**Table S8:** Table giving all unique and shared SNPs that were called in the re-analyzed Aschheim genome (A120), showing the classification as assessed to the other genomes. The classification is based on the preceding parameters regarding a 50 bp window surrounding the SNP: Only SNPs with no uncovered position, no heterozygous SNP and an equal mean coverage with high and low stringent mapping (ratio of 1.00) would be classified as true positive SNPs.

| Unique SNPs | | | | |
|---|---|---|---|---|
| Position | SNP coverage | Uncovered pos. | Heterozygous SNPs | Mean coverage ratio LS/HS |
| 22040 | 4 | 0 | 0 | 1.11 |
| 68401 | 4 | 6 | 1 | 1.11 |
| 90292 | 4 | 0 | 3 | 1.00 |
| 111916 | 3 | 24 | 0 | 1.27 |
| 118450 | 3 | 0 | 1 | 1.00 |
| 118467 | 5 | 0 | 1 | 1.06 |
| 119550 | 4 | 0 | 1 | 1.16 |
| 164957 | 4 | 5 | 1 | 1.24 |
| 206282 | 5 | 0 | 1 | 1.00 |
| 208068 | 4 | 0 | 2 | 1.00 |
| 208083 | 4 | 0 | 2 | 1.00 |
| 216157 | 8 | 0 | 1 | 3.93 |
| 221811 | 10 | 20 | 0 | 2.88 |
| 221849 | 11 | 0 | 2 | 2.76 |
| 222670 | 4 | 0 | 2 | 1.17 |
| 227200 | 4 | 0 | 2 | 1.09 |
| 228754 | 4 | 4 | 1 | 1.00 |
| 229308 | 4 | 8 | 0 | 2.08 |
| 229316 | 4 | 6 | 0 | 2.09 |
| 290016 | 4 | 0 | 2 | 1.00 |
| 318963 | 3 | 13 | 0 | 1.00 |
| 344159 | 4 | 0 | 0 | 1.00 |
| 352307 | 4 | 0 | 2 | 1.19 |
| 362357 | 4 | 17 | 0 | 5.11 |
| 363792 | 5 | 0 | 1 | 1.00 |
| 395774 | 37 | 0 | 0 | 4.65 |
| 439022 | 3 | 0 | 0 | 1.00 |
| 439036 | 3 | 0 | 0 | 1.00 |
| 473451 | 4 | 20 | 1 | 2.76 |
| 490767 | 4 | 7 | 1 | 1.24 |
| 500745 | 4 | 0 | 1 | 1.00 |
| 546706 | 3 | 0 | 0 | 1.03 |
| 567757 | 3 | 0 | 0 | 1.00 |
| 573139 | 4 | 0 | 1 | 0.99 |
| 573636 | 3 | 0 | 0 | 1.00 |
| 609900 | 5 | 13 | 1 | 2.54 |
| 610872 | 3 | 23 | 0 | 2.11 |
| 610908 | 6 | 19 | 1 | 2.13 |
| 705739 | 3 | 0 | 0 | 1.00 |
| 717675 | 4 | 24 | 2 | 1.44 |
| 723724 | 4 | 0 | 2 | 1.00 |
| 740842 | 3 | 6 | 0 | 1.15 |
| 788396 | 4 | 0 | 1 | 1.00 |
| 869369 | 4 | 0 | 2 | 1.26 |
| 912841 | 4 | 0 | 2 | 1.00 |
| 951452 | 4 | 0 | 1 | 1.00 |
| 1000454 | 4 | 2 | 1 | 1.00 |

| | | | | |
|---|---|---|---|---|
| 1017104 | 4 | 0 | 0 | 1.00 |
| 1029706 | 5 | 1 | 1 | 1.00 |
| 1179456 | 4 | 0 | 1 | 1.00 |
| 1278057 | 3 | 22 | 1 | 1.30 |
| 1287721 | 5 | 0 | 1 | 1.00 |
| 1308293 | 37 | 0 | 1 | 1.75 |
| 1349463 | 4 | 0 | 2 | 1.00 |
| 1349472 | 4 | 0 | 2 | 1.00 |
| 1365939 | 4 | 0 | 1 | 1.00 |
| 1371025 | 10 | 2 | 0 | 3.22 |
| 1389874 | 3 | 0 | 1 | 1.09 |
| 1421032 | 27 | 0 | 2 | 1.75 |
| 1444672 | 4 | 0 | 1 | 1.15 |
| 1451459 | 4 | 0 | 2 | 1.27 |
| 1487354 | 4 | 0 | 1 | 1.30 |
| 1498866 | 3 | 0 | 0 | 1.00 |
| 1534269 | 4 | 0 | 1 | 1.00 |
| 1543236 | 4 | 10 | 2 | 1.00 |
| 1555873 | 4 | 0 | 2 | 1.00 |
| 1572689 | 3 | 12 | 0 | 12.78 |
| 1610886 | 3 | 9 | 0 | 1.00 |
| 1711880 | 3 | 0 | 0 | 1.27 |
| 1715867 | 5 | 0 | 0 | 1.20 |
| 1722937 | 6 | 0 | 1 | 1.21 |
| 1745586 | 4 | 0 | 1 | 1.13 |
| 1761696 | 4 | 0 | 1 | 1.00 |
| 1806990 | 4 | 7 | 2 | 1.06 |
| 1854844 | 4 | 14 | 1 | 1.28 |
| 1864422 | 4 | 0 | 2 | 1.00 |
| 1927370 | 3 | 8 | 0 | 1.00 |
| 1943523 | 6 | 7 | 1 | 1.57 |
| 1959365 | 4 | 0 | 1 | 1.00 |
| 2009537 | 4 | 1 | 1 | 1.00 |
| 2072340 | 4 | 0 | 1 | 1.33 |
| 2072914 | 5 | 0 | 1 | 2.38 |
| 2084670 | 4 | 0 | 1 | 1.10 |
| 2336322 | 3 | 12 | 0 | 1.00 |
| 2345720 | 5 | 0 | 1 | 1.00 |
| 2350460 | 4 | 0 | 1 | 1.26 |
| 2361259 | 4 | 5 | 2 | 1.46 |
| 2415931 | 5 | 0 | 1 | 1.00 |
| 2417705 | 4 | 0 | 1 | 1.13 |
| 2459674 | 4 | 0 | 1 | 1.00 |
| 2513785 | 4 | 0 | 2 | 1.00 |
| 2586708 | 4 | 0 | 1 | 1.00 |
| 2591088 | 4 | 0 | 3 | 2.86 |
| 2664231 | 4 | 0 | 2 | 1.00 |
| 2714188 | 5 | 0 | 1 | 1.37 |
| 2836744 | 4 | 6 | 1 | 1.00 |
| 2847001 | 4 | 0 | 1 | 1.36 |
| 2861901 | 3 | 21 | 0 | 1.00 |
| 2861902 | 3 | 20 | 0 | 1.00 |
| 2863957 | 4 | 0 | 2 | 1.00 |
| 2865494 | 3 | 3 | 0 | 3.19 |
| 2972201 | 4 | 0 | 1 | 1.00 |
| 3070014 | 44 | 0 | 3 | 1.68 |
| 3070065 | 56 | 0 | 1 | 1.38 |
| 3074956 | 3 | 23 | 0 | 1.16 |

| | | | | |
|---|---|---|---|---|
| 3090759 | 4 | 0 | 1 | 1.00 |
| 3131243 | 3 | 15 | 0 | 1.00 |
| 3152585 | 4 | 0 | 3 | 1.39 |
| 3205018 | 4 | 0 | 3 | 1.15 |
| 3242622 | 3 | 2 | 0 | 1.21 |
| 3243987 | 4 | 0 | 1 | 1.41 |
| 3247821 | 4 | 0 | 0 | 1.00 |
| 3249954 | 4 | 1 | 2 | 1.00 |
| 3263455 | 4 | 0 | 1 | 1.24 |
| 3266505 | 3 | 10 | 1 | 1.31 |
| 3286005 | 3 | 0 | 1 | 1.00 |
| 3286017 | 4 | 0 | 1 | 1.00 |
| 3288015 | 4 | 0 | 2 | 1.00 |
| 3342782 | 4 | 6 | 1 | 1.00 |
| 3373238 | 5 | 0 | 2 | 1.00 |
| 3392897 | 18 | 2 | 1 | 3.16 |
| 3428541 | 4 | 0 | 1 | 1.00 |
| 3429285 | 6 | 0 | 1 | 3.25 |
| 3479274 | 4 | 0 | 2 | 1.00 |
| 3498406 | 4 | 0 | 1 | 1.03 |
| 3505837 | 4 | 0 | 1 | 1.00 |
| 3531163 | 4 | 0 | 1 | 1.40 |
| 3531197 | 4 | 0 | 1 | 1.14 |
| 3540557 | 4 | 0 | 3 | 1.87 |
| 3543444 | 3 | 0 | 1 | 1.00 |
| 3569690 | 4 | 0 | 1 | 1.25 |
| 3579698 | 4 | 0 | 1 | 1.32 |
| 3588681 | 3 | 8 | 0 | 1.00 |
| 3656870 | 4 | 0 | 1 | 1.11 |
| 3672205 | 8 | 0 | 1 | 1.35 |
| 3813424 | 7 | 23 | 0 | 2.51 |
| 3860098 | 4 | 0 | 1 | 1.07 |
| 3934112 | 3 | 7 | 0 | 1.03 |
| 3956001 | 9 | 0 | 2 | 2.96 |
| 3956018 | 4 | 0 | 5 | 3.46 |
| 4170791 | 45 | 0 | 1 | 1.80 |
| 4199220 | 30 | 0 | 0 | 6.01 |
| 4203620 | 23 | 0 | 1 | 6.18 |
| 4232217 | 3 | 8 | 0 | 1.00 |
| 4342798 | 4 | 12 | 1 | 1.78 |
| 4509055 | 26 | 0 | 0 | 2.46 |
| 4550310 | 3 | 0 | 1 | 1.26 |
| 4575345 | 33 | 0 | 0 | 4.52 |
| Shared SNPs | | | | |
| Position | SNP coverage | Uncovered pos. | Heterozygous SNPs | Mean coverage ratio LS/HS |
| 86824 | 2 | 0 | 0 | 1.00 |
| 189912 | 0 | 23 | 0 | 1.00 |
| 260148 | 0 | 34 | 0 | 2.39 |
| 271114 | 3 | 0 | 0 | 1.48 |
| 485976 | 3 | 0 | 0 | 1.00 |
| 549767 | 2 | 0 | 0 | 1.00 |
| 557841 | 1 | 0 | 0 | 1.80 |
| 727741 | 5 | 0 | 2 | 1.53 |
| 779365 | 2 | 0 | 0 | 1.00 |
| 898980 | 5 | 0 | 1 | 1.16 |
| 1067966 | 6 | 0 | 0 | 1.11 |
| 1211729 | 1 | 0 | 0 | 1.00 |
| 1296743 | 1 | 0 | 0 | 1.58 |

| | | | | |
|---|---|---|---|---|
| 1387701 | 2 | 0 | 0 | 1.00 |
| 1413031 | 0 | 24 | 0 | 4.23 |
| 1434752 | 2 | 6 | 1 | 1.00 |
| 1489055 | 2 | 1 | 0 | 1.42 |
| 1530658 | 1 | 16 | 0 | 1.38 |
| 1609461 | 2 | 0 | 2 | 1.29 |
| 1754708 | 2 | 0 | 1 | 1.50 |
| 1868678 | 3 | 0 | 1 | 1.00 |
| 1956162 | 5 | 0 | 1 | 1.24 |
| 2092152 | 2 | 0 | 0 | 1.85 |
| 2097520 | 1 | 0 | 0 | 1.00 |
| 2352174 | 1 | 0 | 0 | 1.21 |
| 2419529 | 0 | 24 | 0 | 1.00 |
| 2725715 | 3 | 0 | 0 | 1.00 |
| 2753572 | 1 | 0 | 0 | 1.00 |
| 2977542 | 0 | 15 | 0 | 1.00 |
| 3078807 | 1 | 18 | 0 | 1.48 |
| 3274298 | 3 | 0 | 1 | 1.00 |
| 3360963 | 0 | 41 | 0 | 4.04 |
| 3360984 | 0 | 50 | 0 | NA |
| 3398153 | 1 | 0 | 0 | 1.00 |
| 3409414 | 18 | 0 | 0 | 1.00 |
| 3500922 | 3 | 0 | 0 | 1.17 |
| 3535148 | 2 | 0 | 1 | 1.10 |
| 3560088 | 3 | 0 | 0 | 1.00 |
| 3568597 | 3 | 0 | 0 | 1.47 |
| 3750736 | 3 | 0 | 1 | 1.58 |
| 3843195 | 3 | 0 | 0 | 1.08 |
| 3892488 | 1 | 0 | 2 | 1.02 |
| 4066494 | 5 | 0 | 1 | 1.12 |
| 4307755 | 3 | 0 | 0 | 1.00 |
| 4412624 | 2 | 0 | 0 | 1.28 |
| 4423366 | 4 | 0 | 0 | 1.37 |
| 4460688 | 8 | 0 | 0 | 1.07 |
| 4465967 | 8 | 0 | 0 | 1.02 |
| 4628496 | 4 | 0 | 1 | 1.00 |
| 4629169 | 10 | 0 | 0 | 1.18 |

**Table S9:** Radiocarbon and archaeological dates of all individuals from plague-positive contexts in this study and of two published sites. Raw dates were calibrated with IntCal13 in OxCal v4.3.2. Archaeological dates are based on associated grave goods and cemetery stratigraphy. The probability plots for all dates are shown in Fig. S5.

| Site | Individual number | Lab ID | Skeletal element | Conventional Radiocarbon Age BP | $\delta^{13}C$ AMS [‰] | cal AD 2σ | Arch. Dating | Source |
|---|---|---|---|---|---|---|---|---|
| Previously published individuals | | | | | | | | |
| Altenerding | 1175 | AE1175 | Metacarpus | 1541±19 | -21.1 | 427-573 | 530-570 | Feldman et al. 2016 |
| Altenerding | 1176 | AE1176 | Metacarpus | 1563±20 | -22.9 | 425-546 | 530-570 | Feldman et al. 2016 |
| Aschheim | 58 | A58 | Rib | 1563±18 | -18.4 | 426-544 | 580-600 | Harbeck et al. 2013 |
| Aschheim | 76 | A76 | Rib | 1536±18 | -21.6 | 428-580 | 530-570 | Harbeck et al. 2013 |
| Aschheim | 120 | A120 | Rib | 1525±21 | -17.2 | 430-599 | 530-570 | Wagner et al. 2014 |
| Individuals identified in this study | | | | | | | | |
| Dittenheim | 18A | DIT003 | Rib | 1533±25 | -31.2 | 428-595 | 500-600 | This study |
| Edix Hill | Sk405 | EDI001 | - | - | - | - | 500-550 | This study |
| Lunel-Viel | 1 | LVC001 | Femur | 1492±23 | -13.1 | 539-632 | 400-600 | This study |
| Lunel-Viel | 2 | LVC002 | Femur | 1474±24 | -21.7 | 550-640 | 400-600 | This study |
| Lunel-Viel | 3A | LVC003 | Femur | 1471±23 | -19.6 | 555-641 | 400-600 | This study |
| Lunel-Viel | 4 | LVC005 | Femur | 1505+24 | -24.9 | 434-618 | 400-600 | This study |
| Lunel-Viel | 5 | LVC006 | Femur | 1485 ±24 | -17.6 | 543-635 | 400-600 | This study |
| Lunel-Viel | 6 | LVC007 | Femur | 1453±24 | -25.1 | 567-648 | 400-600 | This study |
| Petting | 378 | PET004 | Rib | 1517±17 | -22.3 | 434-600 | 570-600 | This study |
| Petting | 628 | PET005 | Rib | 1487±21 | -23.8 | 542-630 | 530-730 | This study |
| Petting | 630 | PET006 | Rib | 1551±20 | -19.6 | 427-558 | 570-600 | This study |
| Unterthürheim | 129 | UNT002 | Rib | 1555±20 | -19.0 | 427-553 | 550-600 | This study |
| Unterthürheim | 130 | UNT003 | Rib | 1520±17 | -21.9 | 433-599 | 550-600 | This study |
| Unterthürheim | 131 | UNT004 | Rib | 1587±20 | -20.9 | 419-537 | 550-600 | This study |
| Unterthürheim | 132 | UNT005 | Rib | 1505±24 | -24.6 | 434-618 | 550-600 | This study |
| Unterthürheim | 134 | UNT008 | Rib | 1576±23 | -23.2 | 421-542 | 550-600 | This study |
| Unterthürheim | 63 | UNT006 | Rib | 1507±20 | -22.7 | 437-611 | 550-600 | This study |
| Valencia | 41 | VAL001 | Tooth | 1512±22 | -18.2 | 432-610 | 500-700 | This study |
| Waging | 199 | WAG001 | Rib | 1547±16 | -15.7 | 429-562 | 600-700 | This study |
